# A finite element model of pregnancy derived from maternal sonography: effect of uterine and cervical structural properties on cervical mechanical loading

**DOI:** 10.64898/2026.06.22.733744

**Authors:** Erin M. Louwagie, Hannah Z. Haider, Camilo Duarte, Lei Shi, Mirella Mourad, Michael D. House, Helen Feltovich, Kristin M. Myers

## Abstract

Identification and treatment of pregnancies at risk for preterm birth is a central challenge in obstetric research. Many of the known causes of preterm birth originate from mechanical failure in reproductive tissues. To better understand the biomechanical environment of the gravid uterus and its potential contribution to preterm birth, this computational study presents a parametric method for modeling maternal reproductive anatomy during the early second trimester. A finite element modeling approach was built using existing sonographic measurements from early second-trimester maternal anatomy and material properties from published mechanical tests. We applied the same physiologically relevant intrauterine pressure to all models and quantified the resulting tissue stretch. The sensitivity of the stretch in the proximal cervix was explored by varying material properties and sonographic maternal anatomy dimensions. Cervical material properties, particularly the fiber stiffness modulus and ground substance Young’s modulus, were found to have the greatest effect on proximal cervix stretch compared to other material properties and sonographic dimensions. Among the sonographic dimension measurements, those defining the region surrounding the proximal cervix had the greatest effect on proximal cervix stretch, including the curvature of the posterior uterine wall and the thickness of the lower uterine segment. The computational modeling approach presented here enables future patient-specific studies of gravid reproductive tissues to elucidate differences between individuals who do and do not deliver preterm. Additionally, this study is foundational for building digital twins to support future virtual clinical studies on diagnostic and therapeutic device design to prevent preterm birth.

## 1 INTRODUCTION

Accurate prediction and effective prevention of preterm birth (PTB, birth before 37 weeks of gestation) have long eluded clinicians and researchers. PTB can be spontaneous (unplanned) or medically indicated (clinician initiated due maternal or fetal concerns), with 65–70% of PTB being spontaneous [1]. From 2014 to 2022, the combined spontaneous and medically indicated PTB rate in the United States in singleton gestation pregnancies rose 12% (from 7.74% to 8.67%), with the rate of early PTB (before 34 weeks gestation) concurrently rising 4% (from 2.07% to 2.16%) [2]. This is concerning, as PTB is associated with increased neonatal mortality and lifelong morbidity, with worse outcomes among the earliest PTBs [3]. Though there are many risk factors associated with PTB, including race, history of preterm birth, and vaginal infection, assessment of the cervix is a well established approach for PTB prediction [4]. The cervix is a collagenous, cylindrical shaped organ located at the base of uterus [5]. The cervix has a dual mechanical role during pregnancy: it must remain closed and long enough to retain the fetus as it develops, but soften and shorten to allow for vaginal delivery at term [6]. Cervical length (CL), measured via transvaginal sonography between 16 weeks and 23 weeks 6 days of gestation, is an established clinical method to assess PTB risk [4]. Pregnant individuals with CLs*<*25 mm between 16 weeks and 23 weeks 6 days of gestation are diagnosed with a short cervix and considered to be at increased risk of PTB [4]. However, in a study of singleton pregnancies that went on to deliver preterm, only 18% had CL*<*25 mm [7]. Thus, CL measurement alone does not accurately predict PTB.

To improve PTB prediction, a better understanding of the exact mechanisms that lead to PTB is necessary. There are various risk factors associated with PTB, with many of its cited causes stemming from tissue biomechanical failure [8]. The most often cited cause of PTB is spontaneous preterm labor (sPTL), where the uterus begins regular contractions that lead to cervical dilation before 37 weeks of gestation, preceding PTB in approximately 60% of cases [1]. In approximately 40% of all PTB, the fetal membrane (FM), a thin, bilayer tissue enclosing the fetus and amniotic fluid, breaks too soon, known as preterm prelabor rupture of membranes (PPROM) [1, 9]. The cervix itself may be unable to remain closed in the second trimester, even without uterine contractions and/or labor [10]. This inability of the cervix to retain the pregnancy is known as cervical insufficiency (CI), and it is estimated that 1% of pregnancies are affected by CI [10, 11]. Although the diagnosis of CI can be challenging, is known that preterm cervical shortening precedes cervical insufficiency in many cases [8]. Ultimately, these conditions lead to premature cervical dilation and support the concept that PTB is often a biomechanical failure of gravid tissues, with the biomechanical factors leading to these failures being poorly understood [8].

Human pregnancy is a protected environment, and concerns for fetal safety pose ethical, practical, and legal constraints for *in vivo* clinical studies [12]. Hence, *in silico* methods for studying pregnancy biomechanics and its relationship to PTB offer a promising framework for discovering disease mechanisms and designing and derisking biomedical devices. Initial *in silico* research into PTB biomechanics established methods to generate maternal geometry via three dimensional sonography [13, 14] and magnetic resonance images (MRI) [15]. In these studies, high-fidelity three dimensional solid models of pregnant maternal reproductive anatomy were generated via image segmentation [13, 14] and used in finite element (FE) models to study cervical funneling, uterine growth, and cervical deformation with fundal pressure [15]. Fernandez et al. continued this work, where MRI derived uterine and cervical geometries from patients with a long and short cervix were used to show that the FM, FM adhesion, uterine wall, and cervix all contribute to carrying the load of pregnancy [16]. Additionally, Mahmoud et al. developed a generalized two dimensional model of the uterus, cervix, and pelvic bone and were able to simulate cervical funneling with varying hydrostatic pressure and cervical stiffness [17]. Collectively, these studies provide methods to generate high-fidelity models of the pregnant anatomy and give valuable insights into possible mechanisms that could lead to PTB by showing the biomechanical loading state of pregnant reproductive tissues.

Parametric approaches to solid modeling of maternal anatomy using clinically implementable imaging modalities enable large-scale investigations of the relationship between pregnancy biomechanics and PTB outcomes. One approach to generate patient-specific geometries of the gravid uterus and cervix utilizes two-dimensional sonography. Two dimensional sonography is frequently used in prenatal care and presents an opportunity to develop parametric patient-specific solid models of maternal anatomy, where dimension measurements of key anatomical features are collected and used to build a simplified geometry. Sokolowski et al. modeled the uterus as an ellipsoid based on the anterior-posterior, transverse, and longitudinal diameters of the uterus with a uniform wall thickness collected in the anterior uterine wall [18]. Using this method, Sokolowski et al. measured the uterine size of 320 pregnancies at 4 gestational ages to estimate progressive uterine wall tension [18]. Similarly, Westervelt et al. used an ellipsoid to model the geometry of the gravid uterus, but also included the cervix, FM, and a surrounding abdomen [19]. The parametric framework outlined by Westervelt et al. was used to measure and model one pregnant individual from two dimensional sonography images, and explored FE model sensitivity to select parameters, such as cervical stiffness, CL, and intrauterine pressure [19].

Building from previous parametric computational studies, we developed an improved pregnancy finite element analysis (FEA) model based on the sonographic maternal anatomy scans presented in Westervelt et al. The new method built maternal anatomy [20] into a pipeline to computationally simulate tissue deformation under applied intrauterine pressure. The developed pipeline was then implemented in sensitivity studies across all dimensions and uterine and cervical material properties that serve as inputs to the simulations. We present here 1) material models for the uterus and cervix based on existing uniaxial tension and spherical indentation mechanical testing data [21, 22], 2) a solid modeling method incorporating posterior uterine wall curvature to build parametric maternal anatomy for FE simulation, 3) a quantitative approach to compare cervical deformation spatially between models of different sizes, and 4) the sensitivity of the resulting stretch in the proximal cervix to varied tissue material properties and sonographic maternal dimensions.

## 2 METHODOLOGY

### 2.1 Overview

The finite element analysis (FEA) of maternal anatomy required the development of nonlinear tissue material models and a parametric solid modeling approach. Fiber composite constitutive material models were used to describe the material behavior of the uterus and cervix, with material property values determined via inverse FEA (IFEA) from existing uterine and cervical tissue mechanical data. The solid model of pregnant maternal anatomy was based on existing parametric approaches for generating the uterus, cervix, fetal membrane (FM), and abdomen from sonographic dimension measurements. Proximal cervix stretch under applied intrauterine pressure were quantified at varying tissue material property values and maternal sonographic dimension measurements.

### 2.2 Material models

The uterus, cervix, and FM were modeled as a solid mixture of a continuously distributed fiber network embedded in a compressible neo-Hookean ground substance [23, 24, 21, 25, 22, 26, 27]. These fiber-based models capture the material’s asymmetric tension and compression responses, which are critical for capturing the complex three-dimensional stress and stretch at the uterocervical junction and for reducing bending rigidity in the FM [16, 19, 27, 26]. The most recent cervical and uterine material models include features of the collagen fiber crosslinking that help in quantifying cervical remodeling during gestation [21, 22]. These entropic fiber models are computationally expensive. Hence, in this work focusing on structural features of maternal anatomy, we present a phenomenological based fiber strain energy density and corresponding material model fits as described here and previously [23, 25, 24]. The general total strain energy density (Ψ*^T^ ^otal^*) for the uterus, cervix, and FM is given by equation 1:

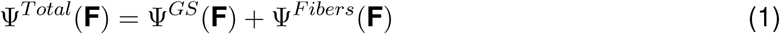

where Ψ*^GS^* is the free-energy density of the neo-Hookean ground substance, Ψ*^F^ ^ibers^* is the free-energy density of the embedded fibers, and **F** is the deformation gradient. The ground substance was modeled as an isotropic, compressible neo-Hookean material, with the strain energy density Ψ*^GS^* given by equation 2 [28]:

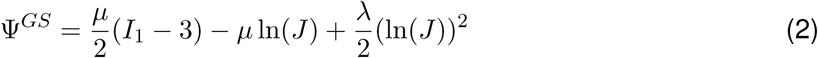

where *I*_1_ is the first invariant of the right Cauchy-Green deformation tensor (**C** = **F***^T^* **F**), *J* is the determinant of **F**, and *µ* and *λ* are the Lamé constants [29]. The Lamé constants are related to Young’s modulus (*E*) and Poisson’s ratio (*ν*) following equations 3 and 4 [29]:

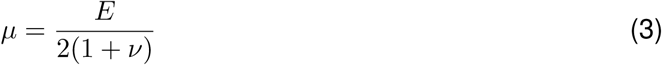

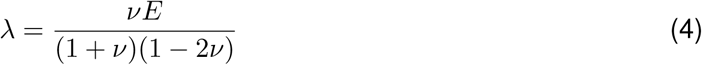

The fibers were modeled as a continuous distribution of fiber orientations, with the general strain energy function for an unconstrained free-energy formulation (Ψ*_r_*) given by equation 5 [30, 31]:

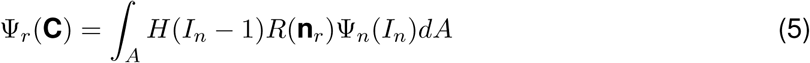

where *H* is the Heaviside unit step function, ensuring the fibers only hold tension. *I_n_* is the normal component of the right Cauchy-Green deformation tensor along the fiber direction unit vector in the reference configuration (**n***_r_*), with *I_n_* = **n***_r_* · **C** · **n***_r_*. The fiber density distribution function, *R*(**n***_r_*), dictates the spatial fractional fiber distribution, and Ψ*_n_* is the strain energy density of the fiber bundle, which is given by equation 6 for a fiber with an exponential-power law [32, 31]:

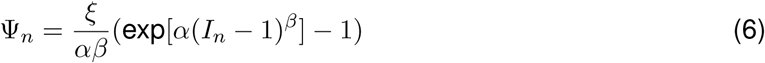

where *ξ* represents the fiber stiffness, *α* is the coefficient of the exponential argument, and *β* is the power of the exponential argument.

#### 2.2.1 Uterus and cervix

The uterus is a muscular, thick-walled organ. The uterus comprises three layers: the innermost endometrium, middle myometrium, and outermost perimetrium. The myometrium is considered to dominate the uterine mechanical response and is composed mainly of smooth muscle cells [33]. The smooth muscle cells of the myometrium are densely packed, and run parallel to a complexly dispersed collagen network [34]. In contrast to the uterus, the cervix is a collagen-dense, cylindrical organ located at the base of the uterus. The cervical stroma makes up the bulk of the cervical tissue and is hypothesized to dominate the cervical mechanical response due to its high levels of collagen [35]. The “three-zone” theory of collagen structure in the cervix proposes an inner and outer zone with longitudinally aligned collagen fibers sandwiching a middle zone with circumferentially aligned fibers, though it is known that these fibers are dispersed around these directions and vary depending on location within the cervix [36, 37, 8]. Additionally, the internal os region of the cervix is comprised of 50-60% smooth muscle cells, indicating that contractility at the internal os is possible [8]. Both the uterus and cervix exhibit active and poro-viscoelastic mechanical behavior [38, 39, 40, 41, 42]. This work will focus on the passive, equilibrium mechanical behavior of uterine and cervical tissue to provide a fundamental basis for future work exploring contractility and tissue material property heterogeneity.

A spherical fiber distribution was used in the uterus and cervix, where fibers rotate and stretch according to principal directions. For a spherical fiber distribution, 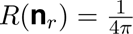 [43]. The exponent *β* was set equal to 2 in accordance with existing work modeling the material behavior of the uterus and cervix [24, 25]. Thus, the final form of the fiber strain energy density for the uterus and cervix (Ψ*^UtCxF^ ^ibers^*) is given by equation 7:

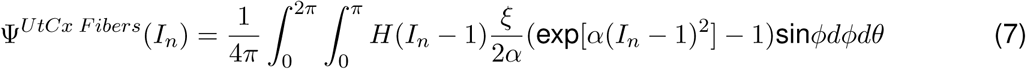

Equation 1 was used in IFEA to fit values for *E*, *ν*, *ξ*, and *α* in simulation to existing force and displacement data from the term pregnant (PG) and non-pregnant (NP) uterus and cervix [21, 22] with Ψ*^F^ ^ibers^* set equal to Equation 7 (full material parameter fitting approach and results presented in Appendix A). The early second-trimester material property values were taken to be the average of the NP and PG values, and are given in Table 1. This assumption is reasonable for mid-gestation cervical properties based on evidence of the non-human primate cervix across gestation [44, 41]. This remodeling trend has also been observed in humans, where the cervix softens most between the late first and middle second trimester [20]. For the sensitivity studies, a small fiber stiffness modulus (*ξ* = 5 kPa) was used in the cervix to mimic a cervix that would be at increased risk of PTB [45, 46]. The uterocervical transition region was assigned a linear gradient of material property values for *E*, *ν*, and *ξ*, with *α* assigned as the average between the uterine and cervical values (1.87).

**Table 1.** Material property values for the cervix, uterus, and fetal membrane in second trimester models as found from mechanical tests of non-pregnant and pregnant human uterus [21] and cervix [22] tissue, and delivered at-term fetal membranes [49]. In the sensitivity studies, a cervical fiber stiffness modulus of 5 kPA was used to mimic a cervix at high-risk for PTB.

| Material Property | Uterus | Cervix | Fetal Membrane |
| --- | --- | --- | --- |
| Neo-Hookean ground substance |  |  |  |
| Young's modulus ($E$, [kPa]) | 1.5 | 1.0 | 1.21 |
| Poisson's Ratio ($\nu$) | 0.27 | 0.225 | 0.38 |
| Embedded fibers |  |  |  |
| Fiber distribution | spherical | spherical | circular |
| Fiber stiffness modulus ($\xi$, [kPa]) | 2.7 | 104.7 (5.0) | 2222 |
| Exponential coefficient ($\alpha$) | 0.74 | 3 | 0.28 |
| Power ($\beta$) | 2 | 2 | 3.33 |

#### 2.2.2 Fetal membrane

The FM is a thin, bilayer tissue comprised of the thick, cellular chorion and the thin, collagen-dense amnion, which is believed to dominate the FM’s mechanical behavior [47]. In this work, the solid model of the FM was built as a monolayer structure, fusing the chorion and the amnion. A previously described material modeling approach was used, with a randomly distributed planar fiber network [27]. A circular fiber distribution was assigned, where 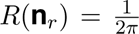 [48]. Thus, the final form of the fiber strain energy density for the FM (Ψ*^F^ ^MF^ ^ibers^*) is given by equation 8:

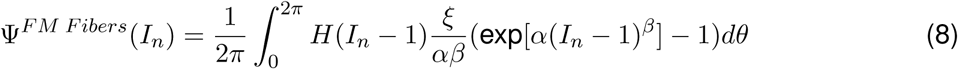

Westervelt used equation 1 in IFEA to fit existing mechanical data of the at-term delivered amnion [49] with Ψ*^F^ ^ibers^* set equal to equation 8 [27]. The resulting at-term parameter values were used in the FM of the early-second trimester model 1.

#### 2.2.3 Abdomen

The abdomen was given a compressible neo-Hookean material model, as described in Equation 2. A Young’s modulus (*E*) of 100 kPa and Poisson’s ratio (*ν*) of 0.4 were assigned based on previous work and sensitivity studies on abdomen stiffness [27, 50].

### 2.3 Patient-specific finite element modeling approach

With the tissue material properties defined, the maternal geometry was built using an existing parametric approach to generate solid models of the uterus and cervix [20], with the FM, abdomen, and a uterocervical transition zone added. The solid geometries were then discretized into elements. Boundary, contact, and loading conditions were defined to mimic the pregnant physiological state.

#### 2.3.1 Parametric geometry

Three-dimensional parametric solid models of gravid human anatomy were built in Solidworks 2019 (Dassault Systémes, Vélizy-Villacoublay, France), including the uterus, cervix, uterocervical transition zone, FM, and surrounding abdomen (Fig. 1) [51]. A *Default Configuration* was first established, where the geometric relations between solid model components were defined, and a *Design Table* was created so that parametric solid models with varying sonographic measurements could be automatically generated. For this study, median sonographic dimension mesurements obtained in the supine position from 29 pregnancies at low-risk for PTB in the early second trimester were used to determine the baseline geometry for comparison [52]. The uterus and cervix were built using our previously published solid modeling approach with parametric measurements collected from two-dimensional sonographic images (listed in Tab. 2 and shown in Fig. 1) [52, 20].

**Fig. 1.**
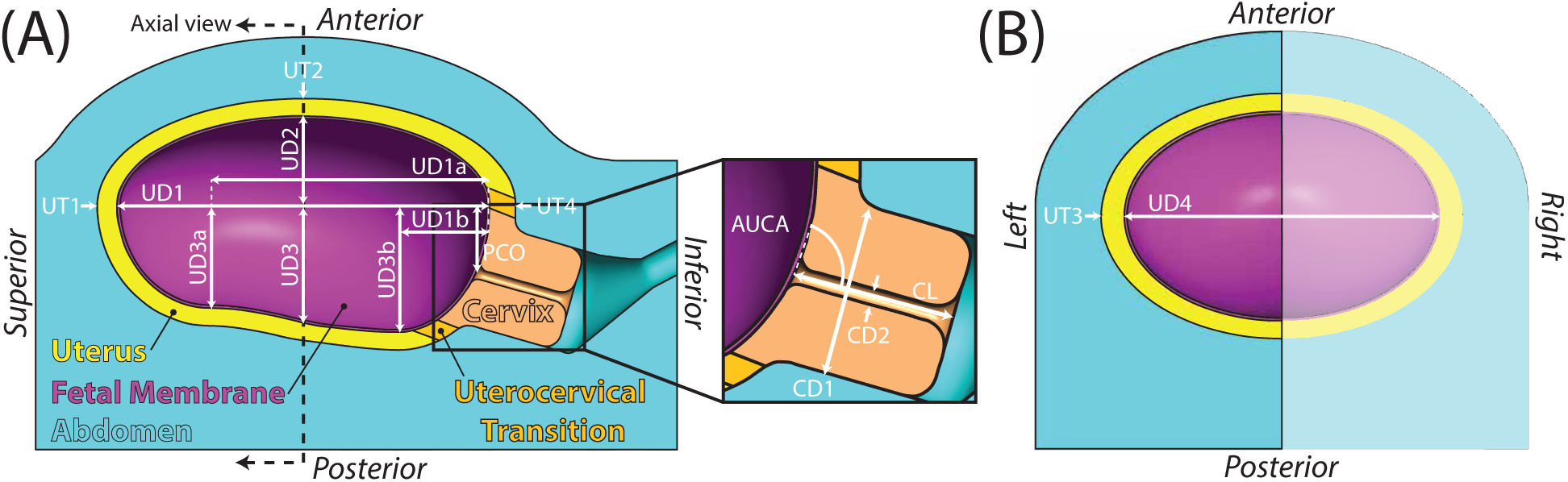
Parametric solid model of human pregnancy from median sonographic measurements in the early second trimester. Sonographic dimensions measurements from the (A) sagittal anatomical plane and (B) axial anatomical plane, as presented in our previous work [52, 20]. The solid model includes, the uterus, cervix, uterocervical transition, fetal membrane, and a surrounding abdomen. Symmetry across the sagittal plane was assumed, and only the left half of the anatomy was used in finite element analysis.

**Table 2.** Definitions of maternal sonographic dimensions used to build the parametric solid models of maternal anatomy, grouped by category.

| Category | Maternal sonographic dimension | Physical meaning |
| --- | --- | --- |
| Intrauterine diameters | UD1 | Longest inferior-superior intrauterine diameter |
|  | UD2 | Anterior intrauterine diameter measured from the midpoint of UD1 |
|  | UD3 | Posterior intrauterine diameter measured from the midpoint of UD1 |
|  | UD4 | Longest left-right intrauterine diameter |
| Posterior uterine dimensions | UD3a | Posterior intrauterine diameter measured at UD1a |
|  | UD1a | Locator dimension for UD3a (75% of UD1 from the inferior) |
|  | UD3b | Posterior intrauterine diameter measured at UD1b |
|  | UD1b | Locator dimension for UD3b (25% of UD1 from the inferior) |
|  | PCO | Perpendicular cervical offset (perpendicular distance between the cervical internal os and UD1) |
| Uterine wall thicknesses | UT1 | Fundal uterine wall thickness |
|  | UT2 | Anterior uterine wall thickness |
|  | UT3 | Left/right uterine wall thickness |
|  | UT4 | Lower uterine segment thickness |
| Cervical dimensions | CL | Cervical length |
|  | CD1 | Outer cervical diameter |
|  | CD2 | Cervical canal diameter |
|  | AUCA | Anterior uterocervical angle (angle between lower uterine segment and cervical canal) |

First, the entire maternal reproductive anatomy (combined uterus and cervix) was built as one solid part, and then the cervical and uterine regions were defined (Fig. 1). For this sensitivity study, the proximal end of the cervical tissue was defined as the outer edge of the fillet between the cervix and uterus, marking the cervical boundary with the uterocervical transition zone (Fig. 1a). This line was turned into a surface using the *Extruded Surface* tool, and its angle in relation to the cervix was set to be as parallel as possible to the cervical canal. The end of the uterocervical transition zone was defined 6 mm perpendicular to this surface using the *Offset Surface* tool. This value was chosen based on existing measurements of isthmus length in nonpregnant uteri, which fall between 5-10 mm [53, 54, 55]. The uterus, uterocervical transition zone, and cervix were separated using these surfaces as the *Trim Tools* in the *Split* function.

The FM was created using the *Thicken* tool, where the intrauterine surface was selected as the surface to thicken. The FM was set to be 0.79 mm thick, which is the mean reported *in vivo* FM thickness in the second trimester measured via sonography for term births (0.79±0.23 mm for term births, 0.77±0.27 mm for preterm births) [56]. For the abdomen, the anterior abdomen was created using the *Lofted Boss/Base* tool, with *Profiles* defined by the anterior uterus with an added abdomen thickness of 20 mm, and a *Guide Curve* defined through the anterior points. Inferiorly, the abdomen was drawn to extend 20 mm plus the CL past the inferior-most point of the outer uterus and posteriorly was drawn to extend from the inferior-superior axis as the sum of the largest posterior intrauterine diameter (UD3, UD3a, or UD3b), UT2, and CL. A vaginal canal was added to connect the cervix to the inferior face of the abdomen. Then, to finish the *Default Configuration*, the entire geometry was cut in the sagittal plane for computational efficiency, retaining the left side of the geometry (Fig. 1b). The sensitivity of the solid modeling approach to various build assumptions, such as abdomen thickness, cervical internal os (IO) fillet radius, and cervical external os (EO) fillet radius was explored, with the modeling approach described here chosen to minimize their effects on simulation results [50].

#### 2.3.2 Finite Element Discretization

Geometries were discretized into elements in Hypermesh 2021.1 (Altair, Troy, MI, U.S.A.) [57]. The meshing scheme was based on a mesh convergence study (Appendix B). The final meshing approach resulted in errors of less than 1% change in strain in the regions of interest, specifically proximal cervix strain and regions close to the cervical IO for the cervix and the FM. The element type and count for the median geometry are given in Table 3.

**Table 3.** Element type and count for each part in the median geometry.

| Part | Element Type | Count |
| --- | --- | --- |
| Cervix | Quadratic tetrahedral | 108740 |
| Fetal membrane | Quadratic hexahedral | 4593 |
| Uterus | Linear hexahedral | 33290 |
| Uterocervical transition | Linear tetrahedral | 22420 |
| Abdomen | Linear tetrahedral | 120688 |

#### 2.3.3 Boundary, contact, & loading conditions

The boundary conditions, contacts, and load were prescribed to reflect the pregnant physiology (Fig. 2). Boundary conditions were assigned such that the superior, inferior, posterior, and mid-sagittal faces were fixed in the normal direction, allowing only in-plane movement. Contact conditions between tissues were assigned to mimic the gravid environment. In uncomplicated pregnancies, it is thought that the entire FM outer surface is tied to the intrauterine surface, and that disruption to the uterine-FM interface could lead to PTB [58]. The detection of specific biomarkers, such as fetal fibronectin (fFN) and phosphorylated insulin-like growth factor binding protein-1 (phIGFBP-1), in cervicovaginal fluid as a clinical test to detect the disruption of the uterine-FM interface has been proposed, but none have shown promise for accurate PTB prediction [59, 60, 61, 62]. To capture the possible loss of FM adhesion in the model, the superior 85% of the FM was tied to the intrauterine wall, while the lower 15% was allowed to slide on the uterus, uterocervical transition zone, and cervix. *In vivo* support to the uterus and cervix is provided by the cardinal and uterosacral ligaments, which connect the cervix to the lateral pelvic wall and sacrum, respectively, represented in the model by a tied contact between the outer cervix and abdomen [63]. In contrast, there are no ligaments that greatly restrict the movement of the uterus, so the outer uterus and uterocervical transition zone were allowed to slide on the abdomen. Because the cervix, uterocervical transition zone, and uterus are one continuous entity, the cervix was fully tied to the uterocervical transition zone, which was fully tied to the uterus (Fig. 2). The effect of varying the contact conditions on the simulation results was explored and accounted for to minimize their impact [50].

**Fig. 2.**
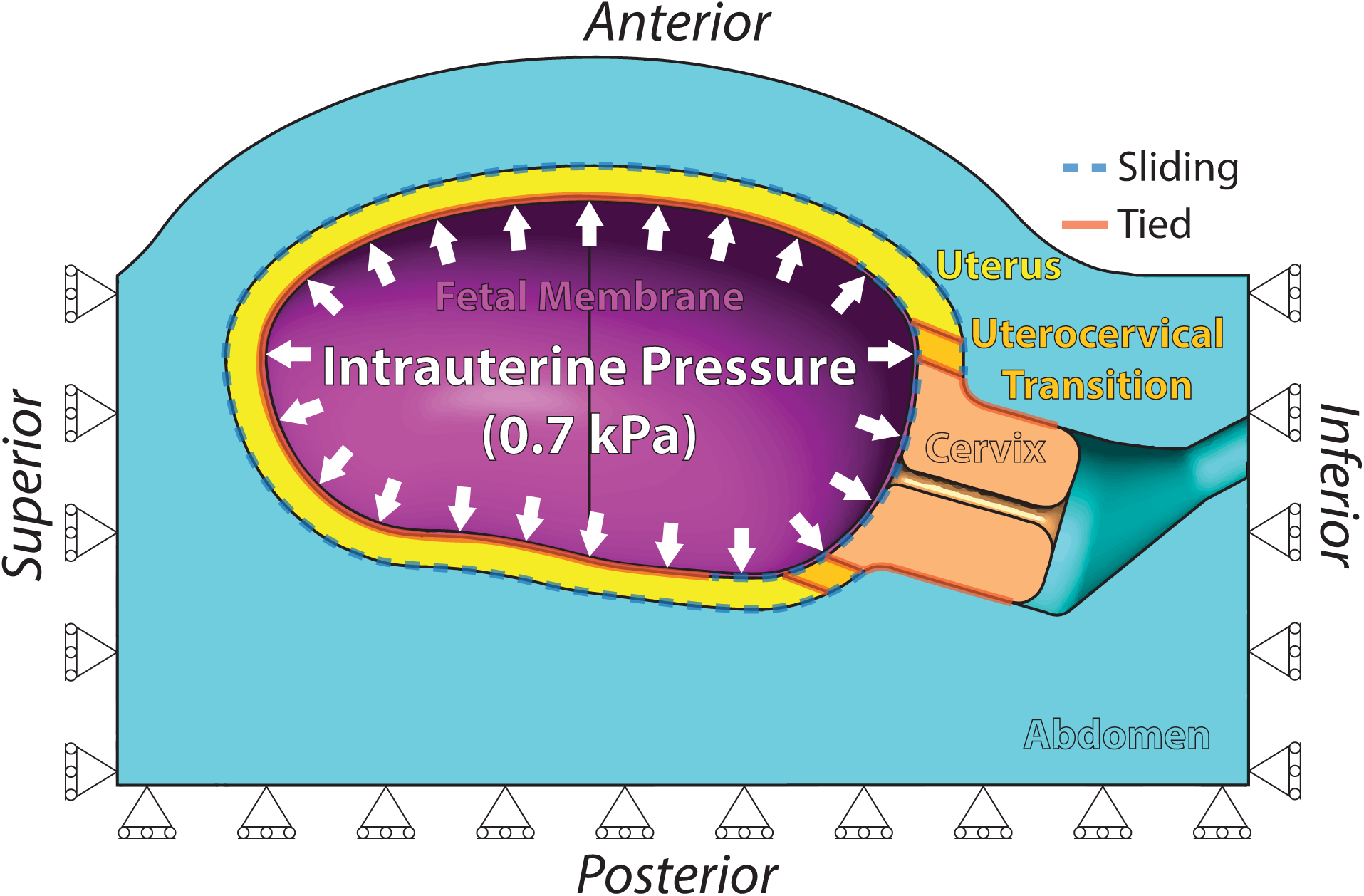
Finite element analysis model boundary conditions and external loading for human pregnancy in the early second trimester (14–17 weeks of gestation). The superior, inferior, and posterior faces were fixed in the normal direction, as well as the sagittal symmetry plane (not shown). Physiologically motivated contact conditions were assigned between tissues, and a gestational age-appropriate intrauterine pressure was applied to the inner fetal membrane surface [64].

The intrauterine pressure (IUP) was assigned to be equal to the amniotic fluid pressure at 16 weeks gestation, given by Equation 9 from Fisk et al. [64]:

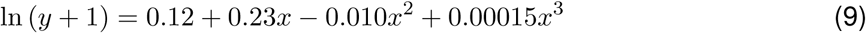

where *y* is the IUP in mmHg and *x* is the gestational age in weeks. This yields an IUP of 0.7 kPa for 16 weeks of gestation.

### 2.4 Stretch analysis

Tissue stretch from specified regions within the maternal anatomy models was investigated (Fig. 3a). First, the maximum 1st principal stretch (*λ*_1_) in the cervical IO was assessed. Then, local cylindrical stretch across the proximal cervix face was calculated, such that the local surface radial, circumferential, and normal stretches could be reported. Regions of interest (ROI) from the mid-stromal region of the anterior, left, and posterior proximal cervix face were defined, and mean stretch within those regions was reported.

**Fig. 3.**
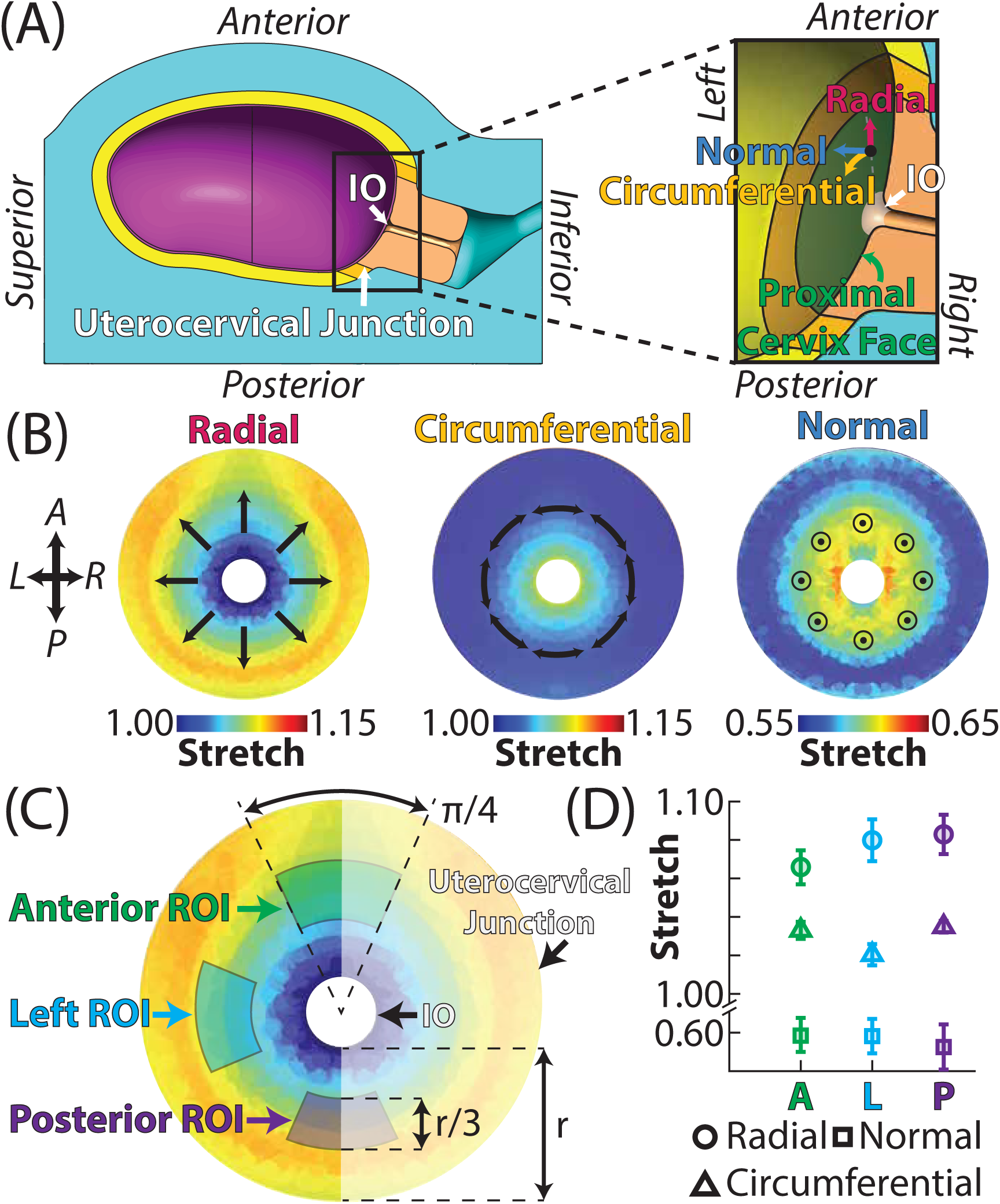
Tissue stretch analysis anatomical locations in the proximal cervix mid-stroma. (A) The proximal cervix face was defined as the nodes between the outer edge of the cervical internal os (IO) fillet and uterocervical transition. The surface radial, circumferential, and normal stretch were calculated for each node in the proximal cervix face. Nodes from the IO were selected to compare maximum *λ*1. The uterocervical junction is located at the interface between the cervix and uterocervical transition. (B) Example heat maps of surface radial, circumferential, and normal stretch in the proximal cervix. (C) Regions of interest (ROI) were selected in the anterior, left, and posterior proximal cervix mid-stroma to span a circumferential distance of *π/*4 and radial distance (*r*) of the middle third (*r/*3) spanning the IO and uterocervical junction boundaries. (D) The average and standard deviation of the surface radial, circumferential, and normal stretch were calculated for each ROI: (A)nterior, (L)eft, and (P)osterior. (R)ight side of proximal cervix is symmetrical to left.\

#### 2.4.1 Cervical internal osj

The cervical IO is a location of interest in studying pregnancy biomechanics, as the opening of the cervical IO is a part of the cervical funneling process [65]. Therefore, the maximum *λ*_1_ in the cervical IO was evaluated to compare the effects of varying material and geometric model inputs (Fig. 3a).

#### 2.4.2 Proximal cervix face

The local surface radial, circumferential, and normal stretch direction was calculated based on the local curvature in radial segments of the proximal cervix face using a custom script in MATLAB 2025 (MathWorks Inc., Natick, MA) [66] (Fig. 3) (URL provided upon publication). The difference in the local surface radial, circumferential, and normal stretch fields was then calculated between models to understand the effects of material and maternal dimension model inputs on cervical stretch under membrane pressurization (Fig. 3a). Heat maps of the local surface radial, circumferential, normal and stretch in the proximal cervix were generated (Fig. 3b), as well as difference heat maps to compare FEA results. ROIs from the anterior, left, and posterior proximal cervix mid-stroma were each defined as a radial segment of width *π/*4 with a length of one-third the distance from the internal boundary to the external boundary (Fig. 3c). The mean and standard deviation of the local surface radial, circumferential, and normal stretches within each mid-stroma ROI were calculated and plotted against the varied model inputs (Fig. 3d). The full process for calculating stretch and generating local cylindrical stretch heat maps is described in Appendix B.

### 2.5 Sensitivity studies

The sensitivity of the FEA model results to changes in cervical and uterine material properties and sonographic dimension measurement values was explored. In all cases, a one-at-a-time approach was followed. The baseline material properties values were based on Tab. 1 and the sonographic dimension measurements were based on existing data from pregnancies in the second trimester [52, 56]. Of note, the calculation to obtain the median early second trimester geometry was based on a dataset that omitted one participant, so the actual median values are slightly different than those reported and used in this work, though all values between the actual and used median were within 1 mm, and no notable differences in the simulation results were found between the actual and used median FEA models (Appendix C Figure 1).

#### 2.5.1 Material properties

The sensitivity of tissue stretch in the cervical IO and proximal cervix face was investigated via a one-at-a-time approach with varying material parameters in the uterus and cervix. The simulation range for each material parameter was based on the nonpregnant (NP) and term pregnant (PG) values, with the lower bound being set as one standard deviation below the mean for the smaller of the NP/PG values and the upper bound as one standard deviation above the mean for the larger of the NP/PG values (Tab. 4). The PG value was smaller than the NP value for every material property except uterine *E*. Exceptions to the use of the standard deviation to find the sensitivity study bounds were made for the *α* lower bound in the uterus and cervix, which was set to zero (*α* cannot be negative), and the *ξ* lower bound in the cervix, which was set to 1 kPa (*ξ* cannot be negative and the smallest fit cervical *ξ* value was 1.52kPa) (Appendix A). The range of stretch values in the proximal cervix produced by varying the material properties from the smallest to largest value was found. A linear regression was individually fit to the cylindrical stretches for all nodes in the anterior, left, and posterior proximal cervix mid-stroma ROIs against the normalized material property value from the smallest (0) to largest (100) material property value in Table 4 (eq. 10, where *X* is the material property value). The same was done for the maximum *λ*_1_ in the cervical IO. This slope was used to evaluate the sensitivity of the model outcome, tissue stretch in the proximal cervix, to each variable. The *p*-value for the slope is reported, with the null hypothesis rejected (slope of 0) when *p <* 0.05.

**Table 4.** Ranges of material property values used in the material property sensitivity study.

| Material property | Uterus range | Cervix range |
| --- | --- | --- |
| Neo-Hookean ground substance |  |  |
| Young's modulus ($E$, [kPa]) | 0.5–2.5 | 0.05–3.5 |
| Poisson's Ratio ($\nu$) | 0.16–0.37 | 0.135–0.285 |
| Embedded fibers |  |  |
| Fiber stiffness modulus ($\xi$, [kPa]) | 0.8–6 | 1–400 |
| Exponential coefficient ($\alpha$) | 0–2.5 | 0–7.7 |

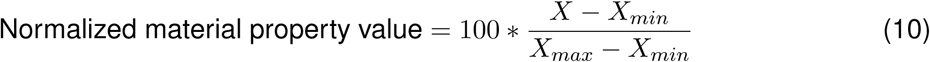

#### 2.5.2 Sonographic dimensions

A one-at-a-time approach was also employed to investigate the effects of varying sonographic maternal anatomy dimension values on the stretch in the cervical IO and the proximal cervix face. The median early second trimester uterine and cervical geometry in the supine position was found using our publicly available dataset of uncomplicated pregnancies, and testing ranges found to cover the majority of the cohort (25th-75th percentiles) and measurement error between sonographers (75th percentile) [52]. The effect of FM thickness was also explored, though it was not collected as a maternal sonographic dimension in our existing cohort. The FM thickness during gestation has been measured via sonographic images, with values from 0.50–1.94 mm reported [67, 56, 68]. However, the material properties used in the FM were collected from amnion samples at term, which has been reported to be approximately 0.1 mm thick [49]. Because the material model used was fit to only the amnion, we modeled FM thicknesses from 0.1–2.00 mm. For all sonographic dimensions, the range of stretch values in the proximal cervix produced by varying the sonographic dimensions from the smallest to largest value was found. A linear regression was individually fit to the cylindrical stretches in the anterior, left, and posterior proximal cervix mid-stroma ROIs against the percent change from the smallest to largest sonographic dimension value in Table 5 and smallest to largest FM thickness value (eq. 11, where *X* is the sonographic dimension). The same was done for the maximum *λ*_1_ in the cervical IO. This slope was used to evaluate the sensitivity of the model outcome, tissue stretch in the proximal cervix, to each variable. The *p*-value for the slope is reported, with the null hypothesis rejected (slope of 0) when *p <* 0.05.

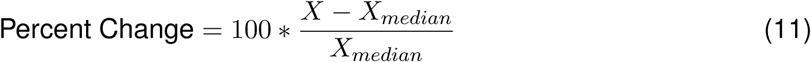

**Table 5.** Sonographic dimension measurement median and 25th-75th %ile values, 75th %ile measurement error value, and range used in sonographic dimension sensitivity study, based on early second trimester anatomy (14-17 weeks gestation) [52].

| Dimension | Median (25<sup>th</sup>-75<sup>th</sup> %ile) | 75<sup>th</sup> %ile Error | Sensitivity Range |
| --- | --- | --- | --- |
| UD1 [mm] | 120.70 (110.73–128.68) | 13.44 | 108.63 (-10%) – 132.76 (+10%) |
| UD2 [mm] | 28.95 (23.36–37.33) | 11.36 | 23.16 (-20%) – 34.74 (+20%) |
| UD3 [mm] | 38.08 (33.55–41.22) | 11.92 | 30.46 (-20%) – 45.69 (+20%) |
| UD4 [mm] | 100.9 (90.76–111.28) | 7.49 | 90.81 (-10%) – 110.99 (+10%) |
| UD3a [mm] | 33.32 (29.18–39.16) | 10.68 | 26.66 (-20%) – 39.98 (+20%) |
| UD1a [mm] | 90.52 (N/A) | N/A | 81.47 (-10%) – 99.57 (+10%) |
| UD3b [mm] | 41.21 (35.82–46.43) | 12.94 | 32.97 (-20%) – 49.45 (+20%) |
| UD1b [mm] | 30.17 (N/A) | N/A | 24.14 (-20%) – 36.21 (+20%) |
| PCO [mm] | 22.01 (15.76–28.39) | 18.15 | 11.00 (-50%) – 33.01 (+50%) |
| UT1 [mm] | 6.48 (5.42–7.93) | 2.94 | 5.18 (-20%) – 7.78 (+20%) |
| UT2 [mm] | 5.76 (4.95–7.20) | 2.75 | 4.61 (-20%) – 6.91 (+20%) |
| UT3 [mm] | 7.31 (6.40–8.16) | 3.22 | 5.84 (-20%) – 8.77 (+20%) |
| UT4 [mm] | 8.51 (6.33–16.01) | 5.31 | 4.25 (-50%) – 17.01 (+100%) |
| CL [mm] | 32.31 (29.88–36.44) | 8.38 | 12.92 (-60%) – 38.77 (+20%) |
| CD1 [mm] | 33.43 (30.71–36.36) | 7.48 | 30.09 (-10%) – 36.77 (+10%) |
| CD2 [mm] | 3.11 (2.59–3.95) | 2.11 | 1.56 (-50%) – 4.67 (+50%) |
| AUCA [°] | 81.16 (60.21–92.37) | 27.36 | 64.92 (-20%) – 97.39 (+20%) |

## 3 RESULTS

The sensitivity of the FEA model results to varying material properties and sonographic dimension values was determined by analyzing stretch in the mid-stromal regions of interest (ROI) in the proximal cervix (anterior, left, and posterior) and maximum 1st principal stretch (*λ*_1_) the cervical internal os (IO). The range of stretch in the proximal cervix produced by varying each material and sonographic parameter was calculated and compared. Linear regressions were fit to the resulting stretch in these regions, with the slopes, R2, and significance reported in Appendix C Tables 1 and 2 for the material property and sonographic dimension values, respectively. The stretch values were visualized against the normalized material property value and the percent change from the median sonographic dimension value. Additionally, heat maps of stretch in the proximal cervix for sonographic dimensions are presented to visualize changes to stretch magnitude and pattern.

### 3.1 Tissue stretch in the proximal cervix is sensitive to uterine and cervical tissue properties

The cervical ground substance Young’s modulus *E* had the largest effect on radial, circumferential, and normal stretch in the proximal cervix mid-stroma and maximum *λ*_1_ in the cervical IO (Figs. 4, 5, and 6, Appendix C Table 1). After cervical *E*, cervical fiber stiffness *ξ* had the next largest effect on radial and circumferential stretch in proximal cervix and maximum *λ*_1_ in the cervical IO (Figs. 4 and 6, Appendix C Table 1). Cervical Poisson’s ratio (*ν*) had the next largest effect on normal stretch in the proximal cervix and maximum *λ*_1_ in the cervical IO after cervical *E* (Figs. 4 and 6, Appendix C Table 1). Uterine *ν* had the third largest impact on radial and circumferential stretch in the proximal cervix of the uterine and cervical material properties (Fig. 5, Appendix C Table 1). The smallest *E* values tested in the cervix (*E* = 0.05 kPa) and uterus (*E* = 0.5) did not fully converge, but were still considered as the 0% normalized material property value for the FEA model without reporting stretch at these points. Heat maps for stretch and difference from baseline in the proximal cervix face for all of the varied material properties are presented in Appendix C.

**Fig. 4.**
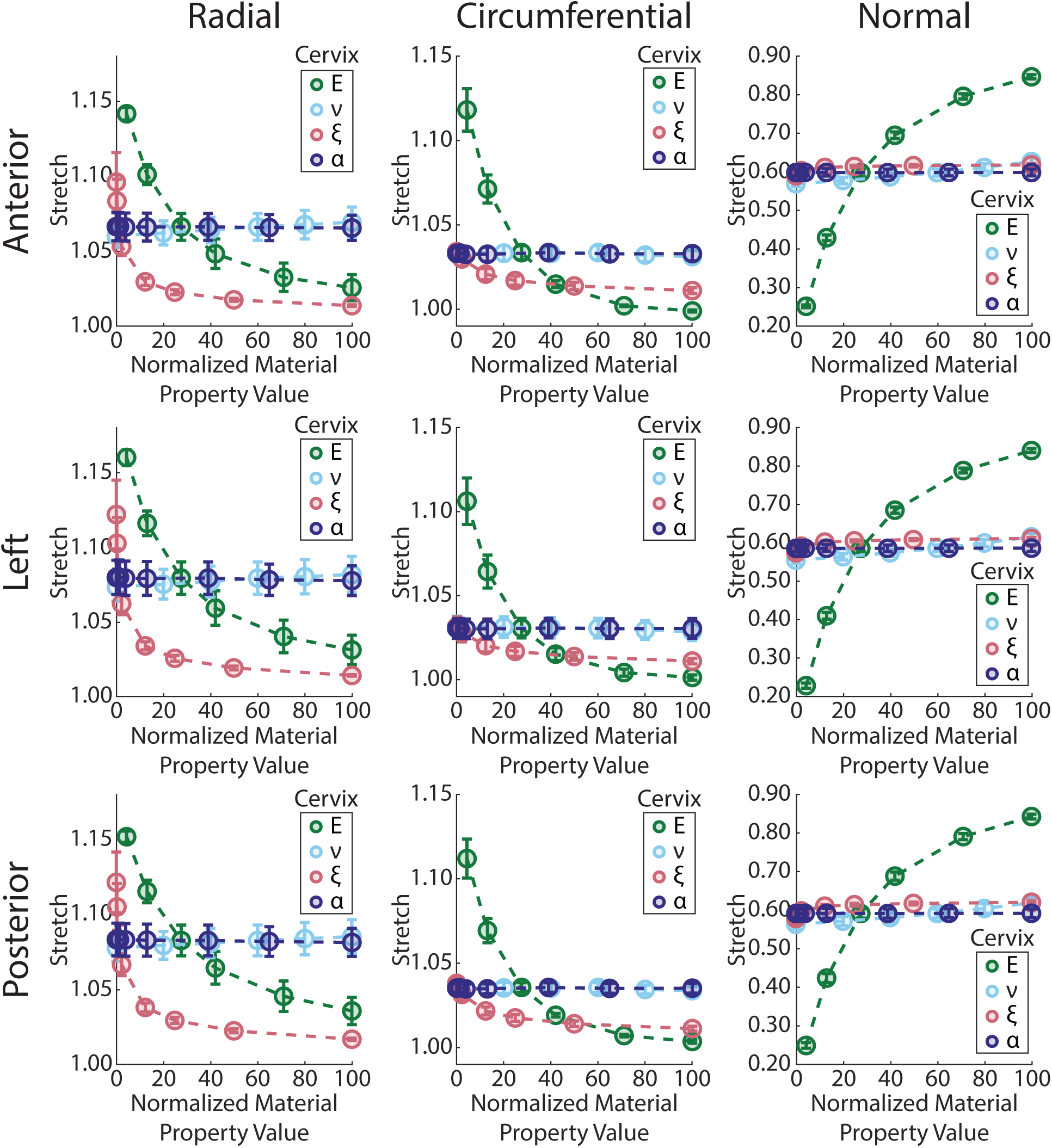
Proximal cervix mid-stroma stretch was highly sensitive to cervical. *E* **and** *ξ*. Stretch in the anterior (row 1), left (row 2), and posterior (row 3) ROIs of the proximal cervix mid-stroma with varying cervical material property values in the surface radial (column 1), circumferential (column 2), and normal (column 3) directions (*E* is the ground substance Young’s modulus, *ν* is the ground substance Poisson’s ratio, *ξ* is the fiber stiffness modulus, and *α* is the fiber exponential coefficient). Normalized material property value indicates the cervical material property’s position between the lower and upper bound tested for the given cervical material property (Tab. 4).

**Fig. 5.**
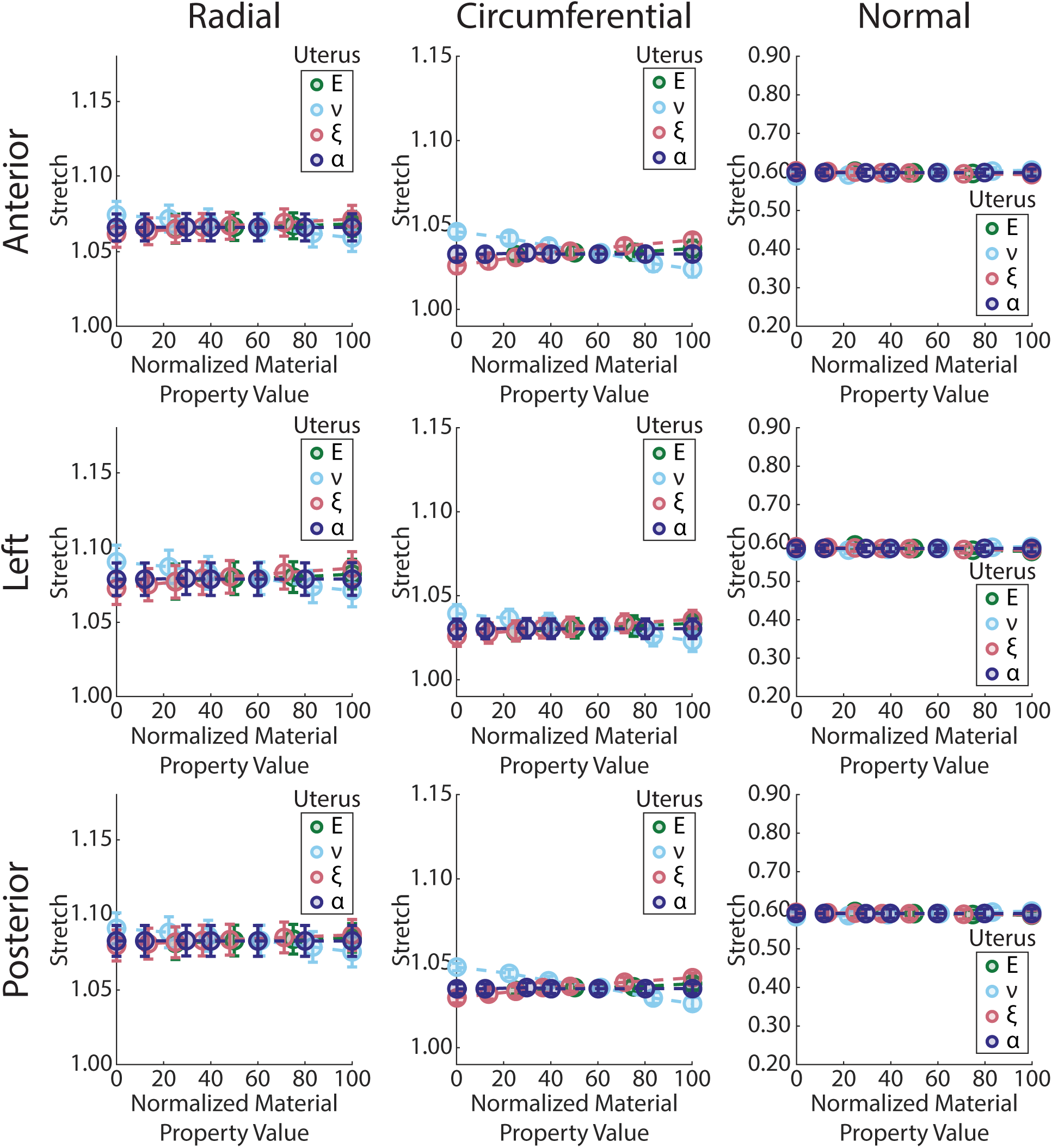
Proximal cervix mid-stroma stretch was not sensitive to uterine material properties. Stretch in the anterior (row 1), left (row 2), and posterior (row 3) ROIs of the proximal cervix mid-stroma with varying uterine material property values in the surface radial (column 1), circumferential (column 2), and normal (column 3) directions. Normalized material property value indicates the uterine material property value’s position between the lower and upper bound tested for the given uterine material property (Tab. 4).

**Fig. 6.**
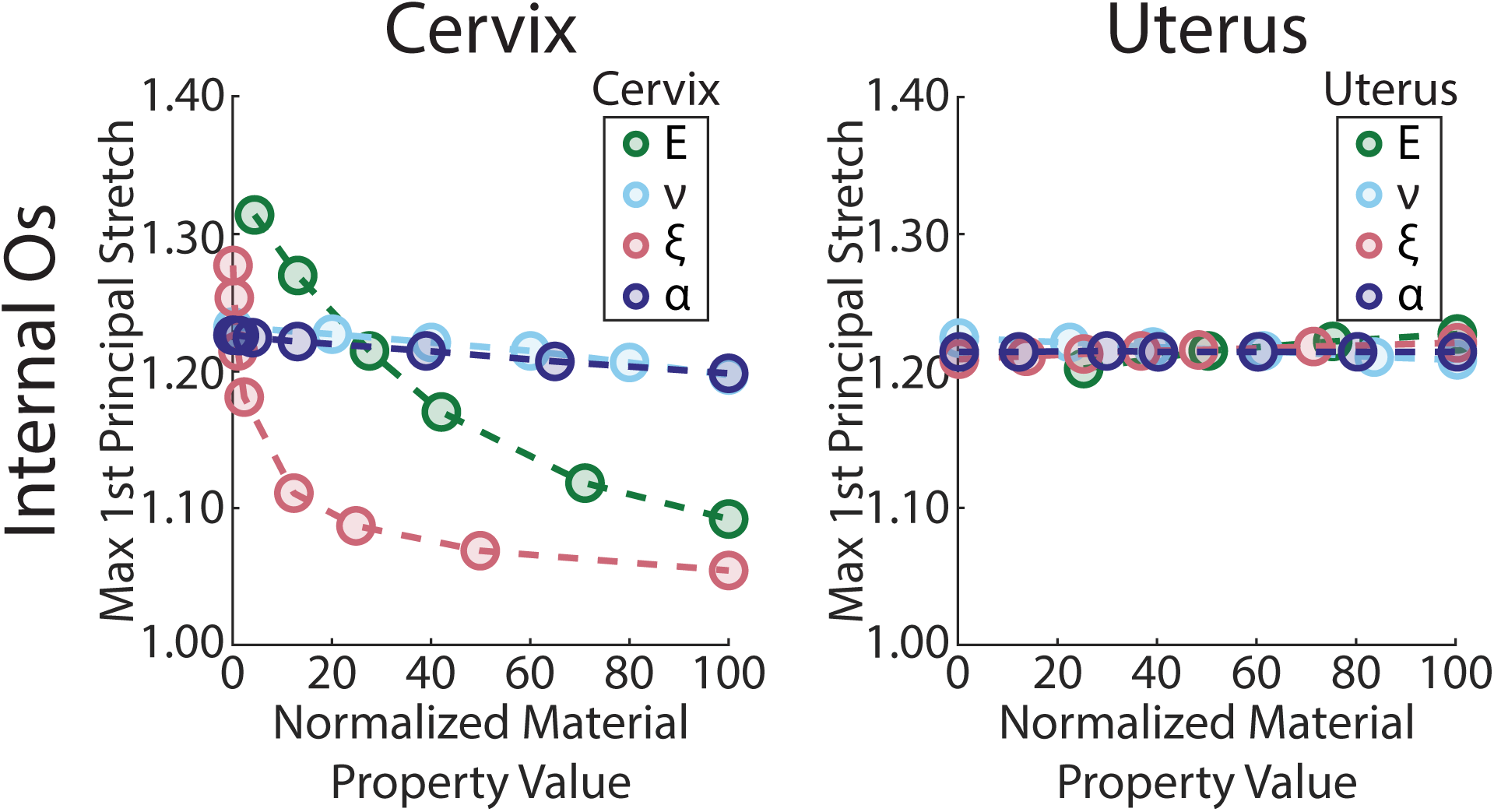
Cervical *E* and *ξ* have the greatest effect on maximum *λ*_1_ in the cervical internal os. Maximum 1st principal stretch (*λ*1) in the cervical internal os with varying cervical (column 1) and uterine (column 2) material property values (*E* is the ground substance Young’s modulus, *ν* is the ground substance Poisson’s ratio, *ξ* is the fiber stiffness modulus, and *α* is the fiber exponential coefficient). Normalized material property value indicates the cervical or uterine material property’s position between lower and upper bound tested for the given cervical or uterine material property (Tab. 4).

#### 3.1.1 Tissue stretch in the proximal cervix is highly sensitive to cervical ground substance and fiber stiffness

Cervical *E* and *ξ* produced greater changes to stretch in the anterior, left, and posterior proximal cervix mid-stroma than cervical *ν* and *α* (Fig. 4, Appendix C Figures 2–5, Appendix C Table 1). In the radial direction, increasing cervical *E* from 0.2–3.5 kPa produced a similar magnitude decrease in stretch in the anterior, left, and posterior mid-stroma (−0.12, −0.13, and −0.12, respectively) and slightly smaller magnitude decreases in stretch were found with cervical *ξ* from 1–400 kPa (−0.08, −0.11, and −0.10, respectively). In the circumferential direction, increasing cervical *E* from 0.1–3.5 kPa produced a similar magnitude change in the anterior, left, and posterior mid-stroma (−0.12, −0.10, and −0.11, respectively) and much smaller magnitude decreases were found with cervical *ξ* from 1–400 kPa (−0.02, −0.02, and −0.03, respectively). In the normal direction, increasing cervical *E* from 0.2–3.5 kPa produced similar magnitude change in the anterior, left, and posterior mid-stroma (0.60, 0.61, and 0.59, respectively).

Cervical *E* and *ξ* also produced greater changes to maximum *λ*_1_ in the cervical IO than cervical *ν* and *α* (Fig. 6, Appendix C Table 1). Increasing cervical *E* from 0.2–3.5 kPa produced a decrease of −0.22 in maximum *λ*_1_ in the cervical IO. Increasing cervical *ξ* from 1–400 kPa also produced a decrease of −0.22 in maximum *λ*_1_ in the cervical IO. Comparatively minimal changes to maximum *λ*_1_ in the cervical IO were found with increasing cervical *ν* from 0.135–0.285 and cervical *α* from 0–7.7 (−0.04 and −0.03, respectively).

#### 3.1.2 Tissue stretch in the proximal cervix is less sensitive to varying uterine material properties

Compared to the cervix, uterine material properties did not produce large changes to stretch in the anterior, left, and posterior proximal cervix mid-stroma (Fig. 5, Appendix C Figures 6–9, Appendix C Table 1). Of the uterine material properties, the greatest changes to proximal cervix stretch were found with uterine *ν* and *E*. In the radial direction, increasing uterine *ν* from 0.16–0.34 produced a similar magnitude decrease in stretch in the anterior, left, and posterior mid-stroma (−0.02, −0.02, and −0.02, respectively). In the circumferential direction, increasing uterine *ν* from 0.16–0.34 produced a similar magnitude decrease in stretch in the anterior, left, and posterior mid-stroma (−0.02, −0.02, and −0.02, respectively). In the normal direction, increasing uterine *ν* from 0.16–0.34 produced a similar magnitude increase (less compression) in the anterior, left, and posterior mid-stroma (0.02, 0.01, and 0.01, respectively), and uterine *E* from 1–2.5 kPa produced a slightly larger magnitude change in normal stretch magnitude than *ν* in the left mid-stroma (−0.02).

**Fig. 7.**
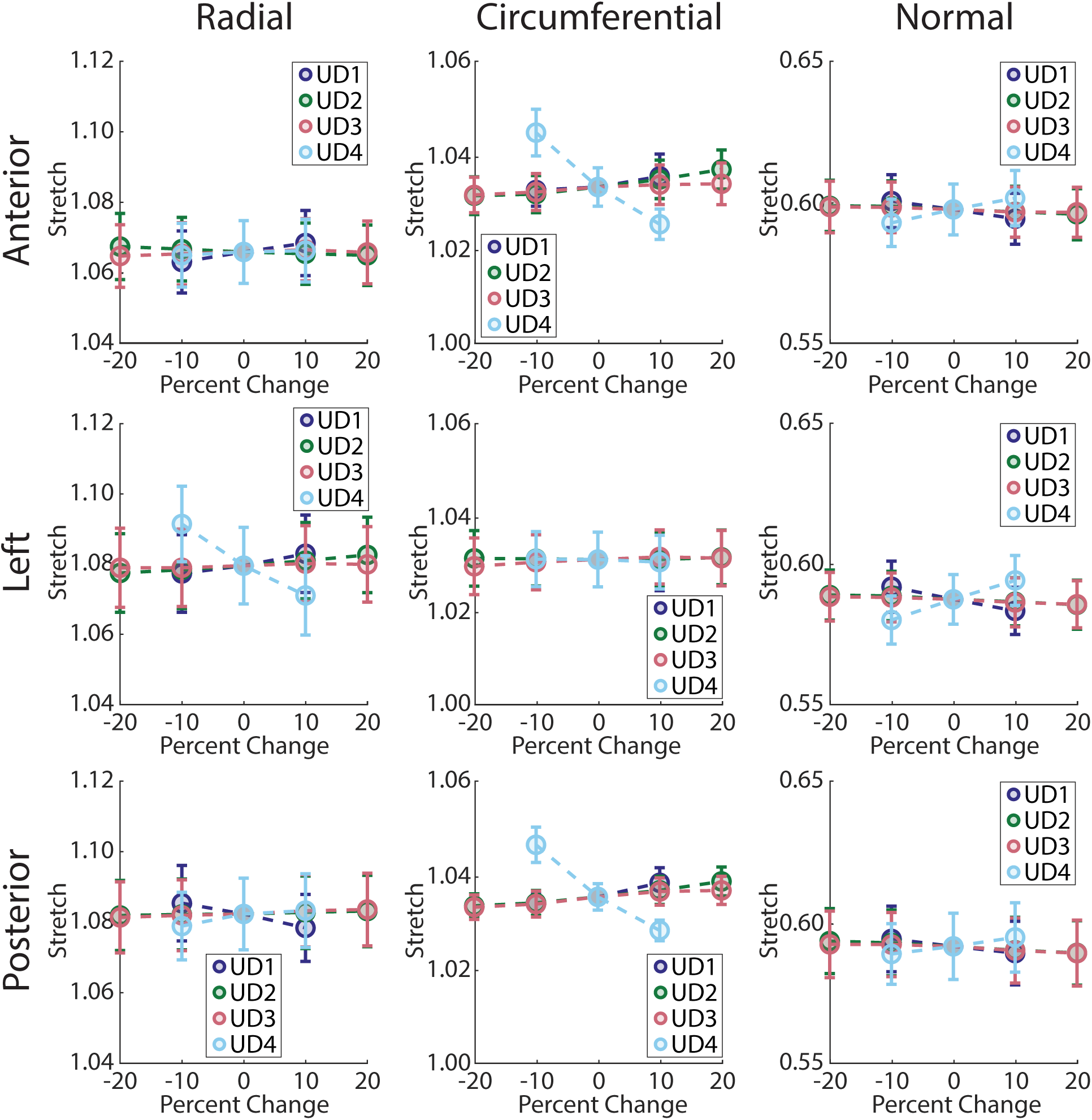
Of the intrauterine diameters, stretch in the proximal cervix mid-stroma was most sensitive to the left-right intrauterine diameter (UD4). Stretch in the anterior (row 1), left (row 2), and posterior (row 3) ROIs of the proximal cervix mid-stroma with varying uterine diameter value in the surface radial (column 1), circumferential (column 2), and normal (column 3) directions. Percent change indicates the percentage change from median for the given intrauterine dimeter (Tab. 5).

**Fig. 8.**
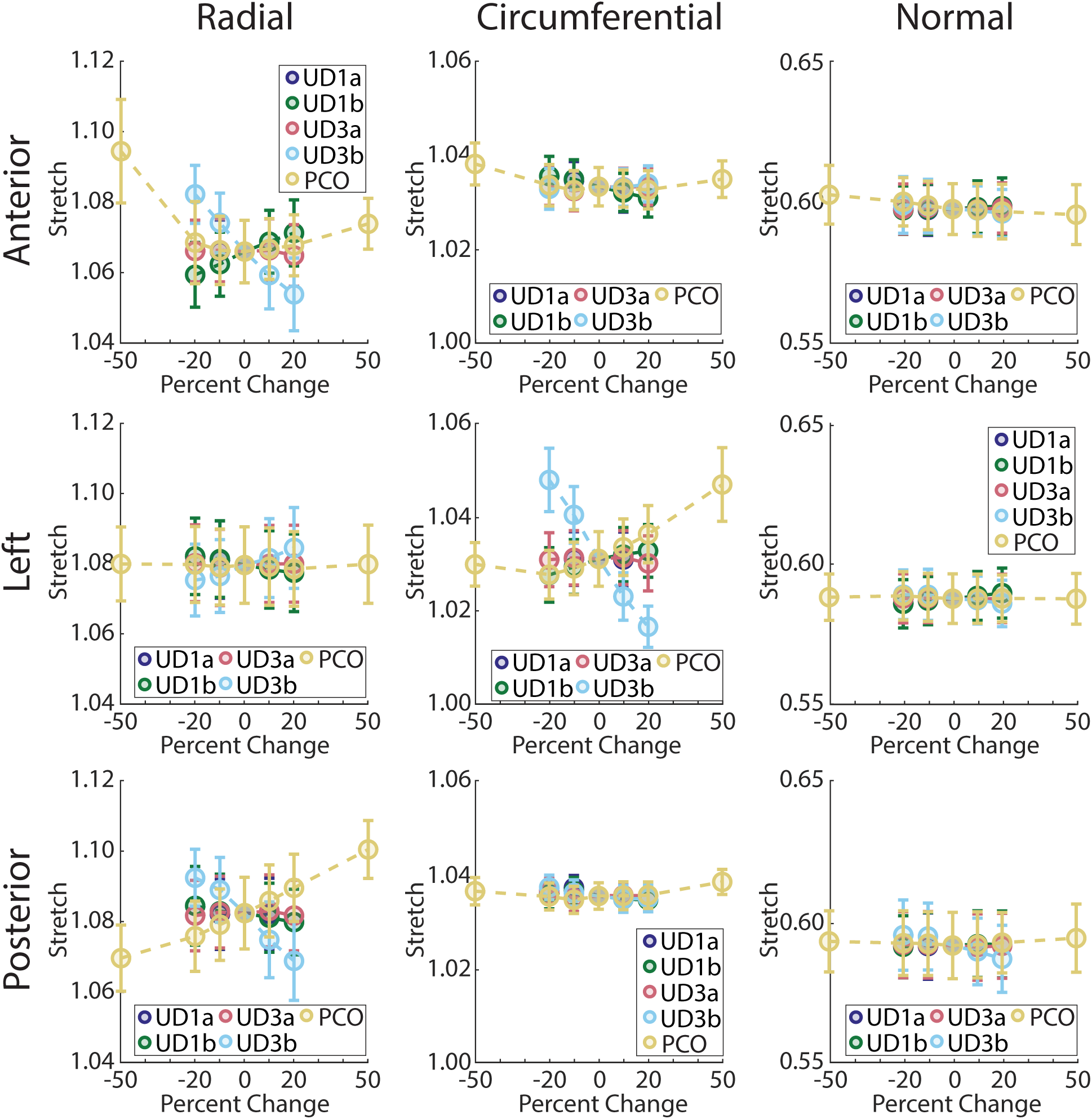
Of the posterior uterine dimensions, stretch in the proximal cervix mid-stroma was most sensitive to the posterior uterine diameter on the inferior half of the uterus (UD3b) and the perpendicular cervical offset (PCO). Stretch in the anterior (row 1), left (row 2), and posterior (row 3) ROIs of the proximal cervix mid-stroma with varying posterior dimension value from median in the surface radial (column 1), circumferential (column 2), and normal (column 3) directions. Percent change indicates the percentage change from median for the given posterior uterine dimension (Tab. 5).

**Fig. 9.**
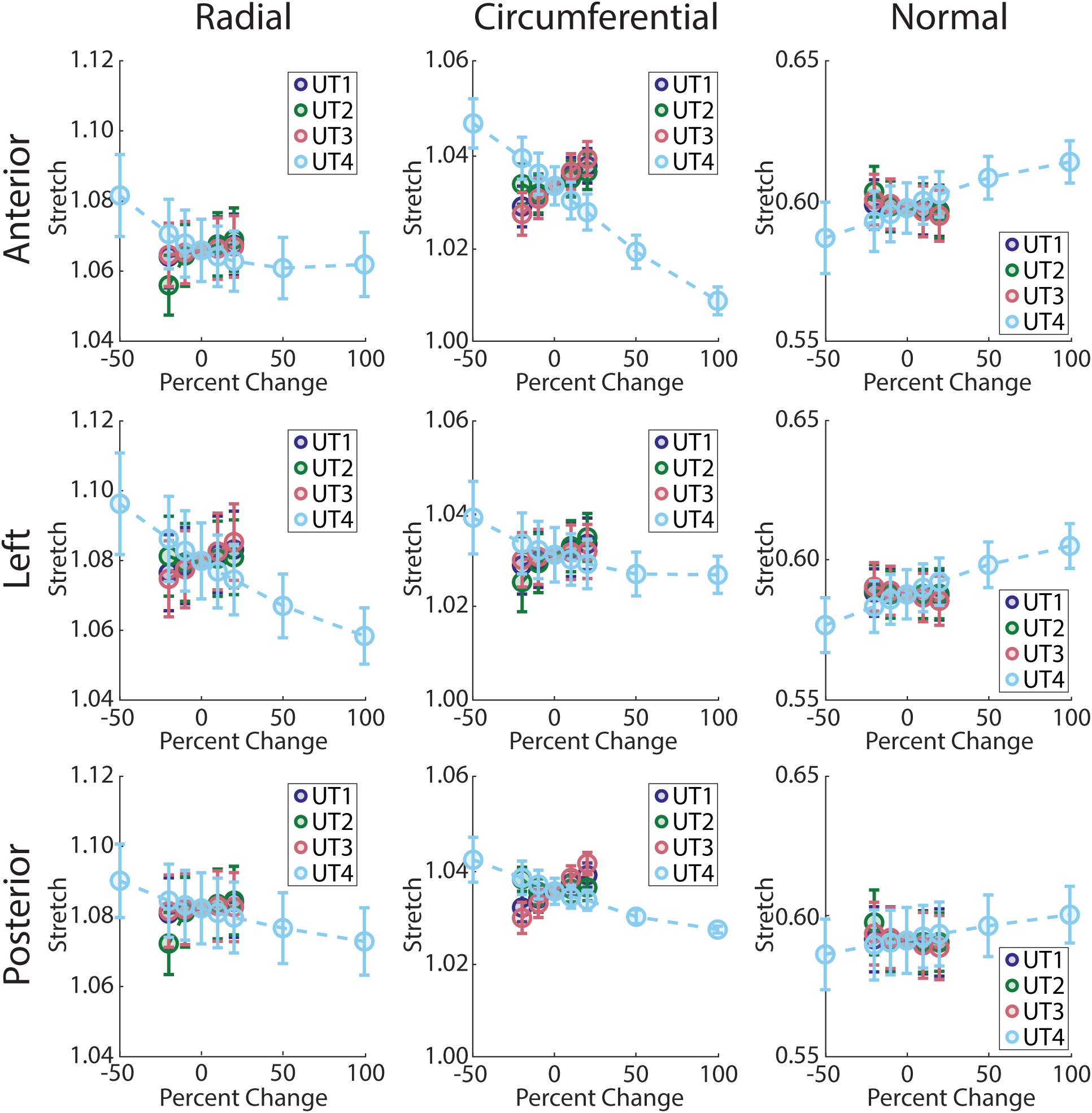
Of the uterine wall thicknesses, stretch in the proximal cervix was most sensitive to the lower uterine segment thickness (UT4). Stretch in the anterior (row 1), left (row 2), and posterior (row 3) ROIs of the proximal cervix mid-stroma with varying uterine wall thickness value in the surface radial (column 1), circumferential (column 2), and normal (column 3) directions. Percent change indicates the percentage change from median for the given uterine wall thickness (Tab. 5).

Uterine *E* produced greater changes to maximum *λ*_1_ in the cervical IO than uterine *ν*, *ξ*, and *α* (Fig. 6, Appendix C Table 1). Increasing uterine *E* from 1–2.5 kPa produced an increase of 0.03 in maximum *λ*_1_ in the cervical IO. Comparatively smaller changes to maximum *λ*_1_ in the cervical IO were found with increasing uterine *ν* from 0.16–0.34, uterine *ξ* from 0.8–6 kPa, and uterine *α* from 0-2.5 (−0.02, 0.01, and 0.00).

### 3.2 Tissue stretch in the proximal cervix is sensitive to maternal anatomical sonographic dimensions

Maternal anatomical sonographic dimensions that describe the region surrounding the proximal cervix had the greatest effect on radial, circumferential, and normal stretch in the proximal cervix and maximum 1st principal stretch (*λ*_1_) in the cervical internal os (IO) (Figs. 7, 8, 9, 10, and 11, Appendix C Table 2). Of the intrauterine diameters, the left-right intrauterine diameter (UD4) had the greatest effect on stretch in the proximal cervix mid-stroma, while the posterior intrauterine diameter (UD3) had the greatest effect on stretch in the cervical IO. Of the posterior intrauterine dimensions, the perpendicular cervical offset (PCO) and the posterior uterine dimension on the inferior half of the uterus (UD3b) had the greatest effect on stretch in the proximal cervix mid-stroma, while PCO produced slightly greater changes to stretch in the cervical IO. Of the uterine wall thicknesses, the lower uterine segment thickness (UT4) produced the greatest changes to stretch in the proximal cervix and cervical IO. Of the cervical dimensions, the cervical length (CL) produced the greatest change to stretch in the proximal cervix mid-stroma. Similar magnitude changes to stretch in the cervical IO were found for all uterine wall thicknesses. The changes to stretch magnitude induced by varying maternal sonographic dimensions were generally not uniform across the anterior, left, and posterior mid-stroma, resulting in an overall change to stretch pattern with varying maternal sonographic dimensions. Stretch in the proximal cervix mid-stroma and cervical IO were also sensitive to varied FM thickness (Fig. 12, Appendix C Table 2). The smallest FM thickness model attempted (0.1 mm) did not converge fully.

**Fig. 10.**
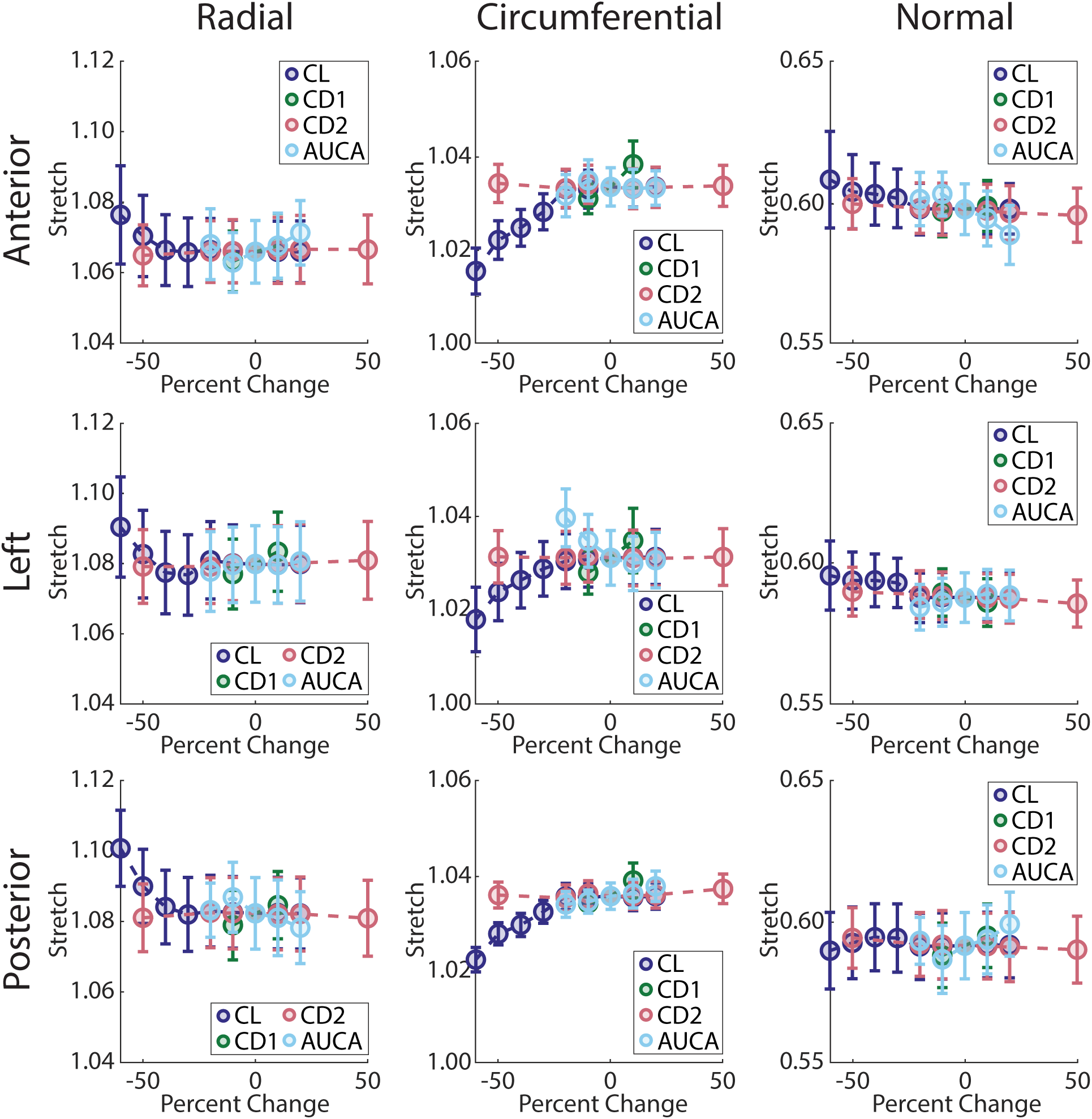
Of the cervical dimensions, stretch in the proximal cervix was most sensitive to the cervical length (CL). Stretch in the anterior (row 1), left (row 2), and posterior (row 3) ROIs of the proximal cervix mid-stroma with varying cervical dimension value in the surface radial (column 1), circumferential (column 2), and normal (column 3) directions. Percent change indicates the percentage change from median for the given cervical dimension (Tab. 5).

#### 3.2.1 Tissue stretch in the proximal cervix is not highly sensitive to intrauterine diameters

Of the intrauterine diameters, the left-right intrauterine diameter (UD4) produced greater changes to stretch in the anterior, left, and posterior proximal cervix mid-stroma than the inferior-superior (UD1), anterior (UD2), and posterior (UD3) intrauterine diameters (Figs. 7 and 13, Appendix C Figures 10–12, Appendix C Table 2). In the radial direction, increasing UD4 from 90.81–110.99 mm produced a −0.02 decrease in stretch in the left mid-stroma. In the circumferential direction, increasing UD4 from 90.81–110.99 mm produced a similar magnitude decrease in stretch in the anterior and posterior mid-stroma (−0.02 and −0.02, respectively). In the normal direction, increasing UD4 from 90.81–110.99 mm and increasing UD1 from 108.63–132.76 mm produced magnitude changes in stretch of approximately 0.01 in the anterior, left, and posterior mid-stroma (less compression with increased UD4, more compression with increased UD1). Mid-stromal changes in stretch are reflected in the heat maps of stretch with varied UD4 (Fig. 13). Radial stretch decreased with increasing UD4 in the left and right proximal cervix. Circumferential stretch decreased with increasing UD4 in the anterior and posterior proximal cervix. Normal stretch increased (less compression) with increasing UD4 across the proximal cervix.

**Fig. 11.**
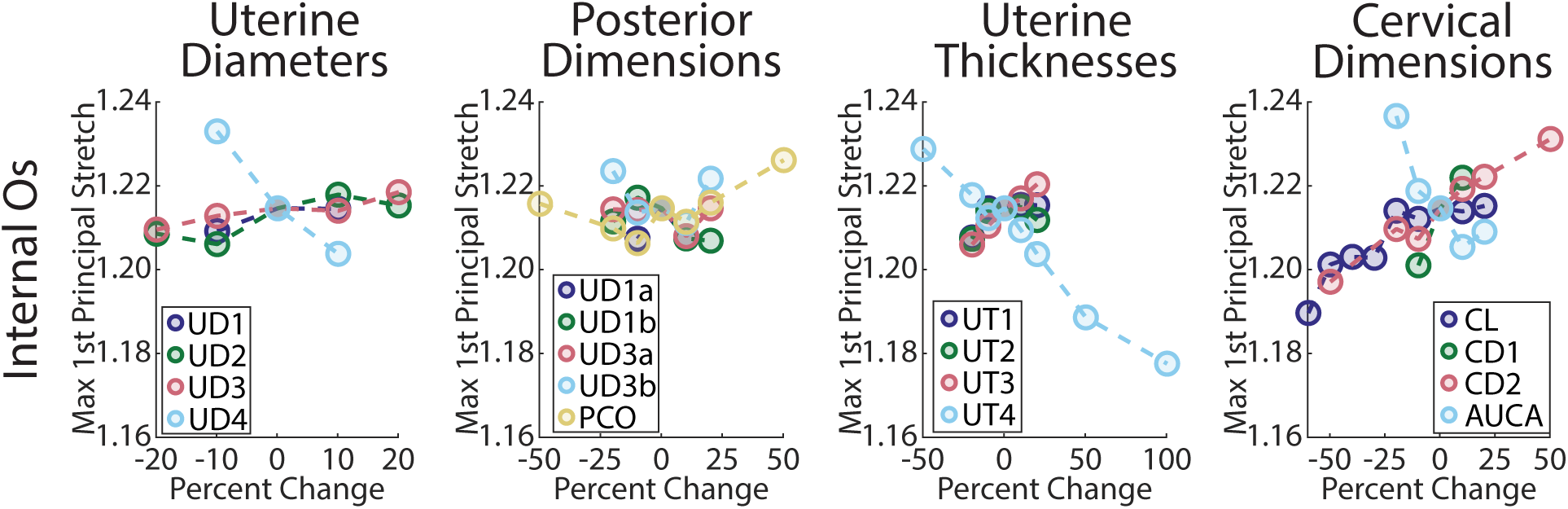
Stretch in the cervical internal os was sensitive to maternal sonographic dimensions. Maximum 1st principal stretch (*λ*1) in the cervical internal os (row 2) with varying uterine diameters (column 1), posterior uterine dimensions (column 2), uterine wall thicknesses (column 3), and cervical dimensions (column 4). Percent change indicates the percentage change from median for the given sonographic dimension (Tab. 5).

**Fig. 12.**
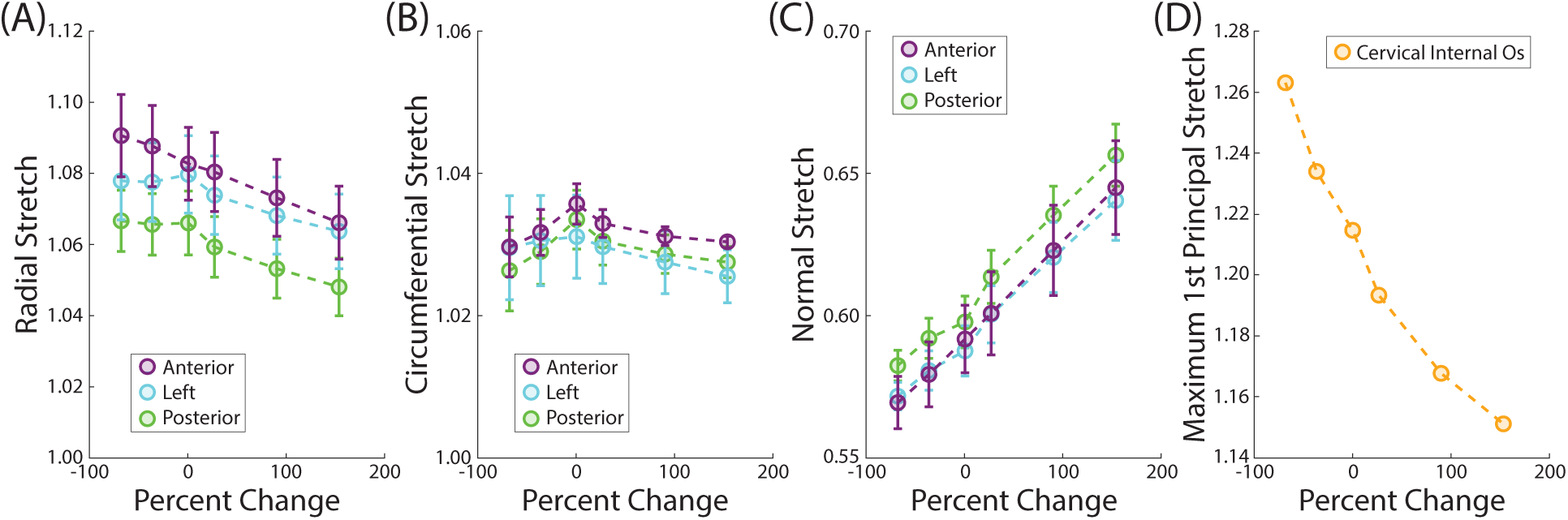
Stretch in the proximal cervix mid-stroma and the cervical internal os (IO) were sensitive to changes in FM thickness. Stretch in the (A) radial, (B) circumferential, and (C) normal directions in the ROIs of proximal cervix mid-stroma. (D) Maximum 1st principal stretch (*λ*1) in the cervical internal os. Percent change indicates the percentage change from the used FM thickness in the second trimester (0.79 mm). [56].

**Fig. 13.**
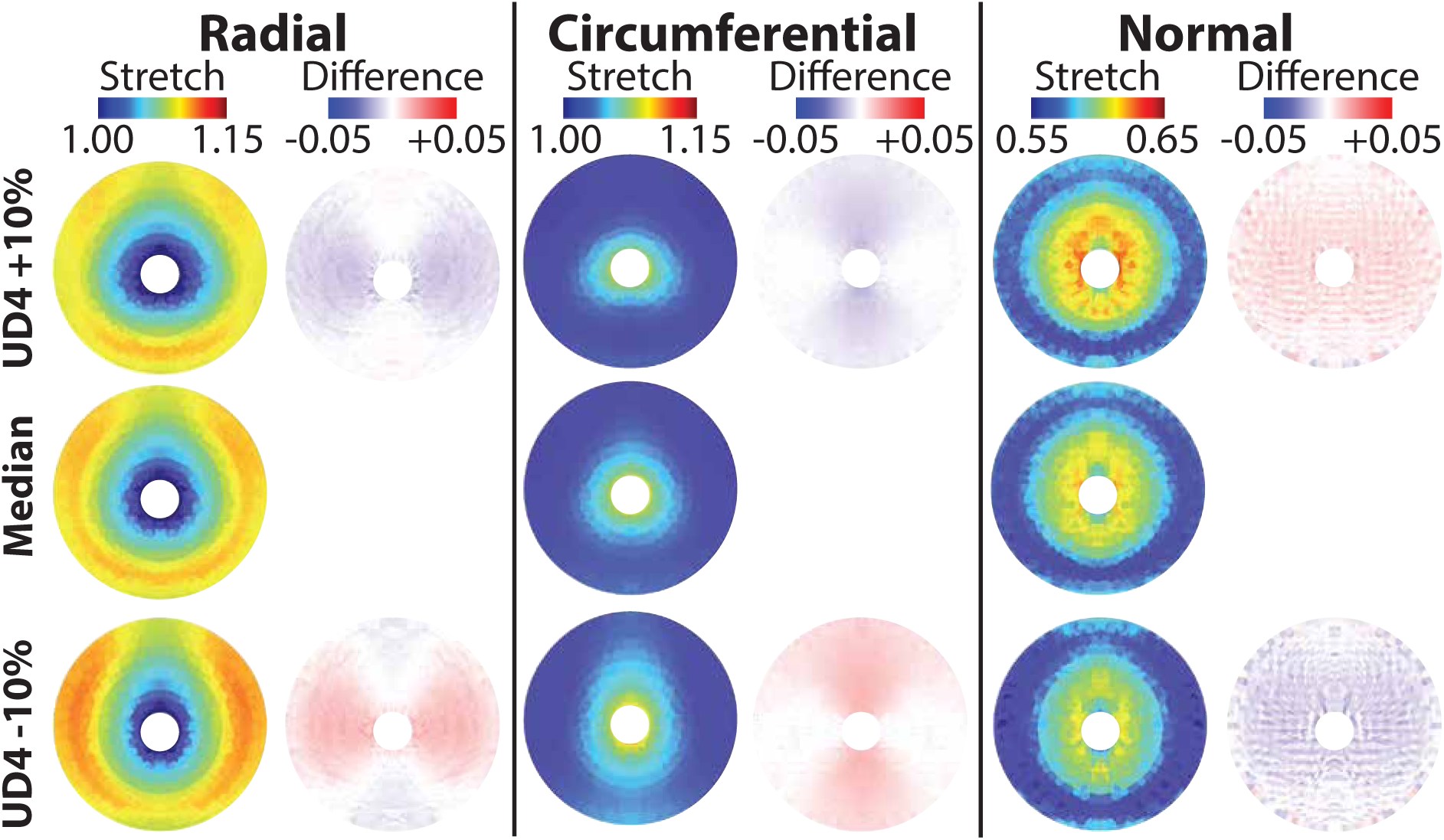
Increasing UD4 produced a decrease in overall proximal cervix deformation. Heat maps of proximal cervix face stretch and difference from median in the surface radial, circumferential, and normal direction for varied left-right intrauterine diameter (UD4).

UD4 produced greater changes to maximum *λ*_1_ in the cervical IO than UD1, UD2, and UD3 (Fig. 11, Appendix C Table 2). Increasing UD4 from 90.81–110.99 mm produced a decrease of −0.03 in maximum *λ*_1_ in the cervical IO. The same magnitude change in maximum *λ*_1_ (0.01) in the cervical IO was found with varied UD1 (108.63–132.76 mm), UD2 (23.16–34.74 mm), and UD3 (30.46–45.69 mm).

#### 3.2.2 Tissue stretch in the proximal cervix is sensitive to posterior uterine dimensions near the cervix

Of the posterior uterine dimensions, which control the curvature of the posterior uterine wall (UD1a, UD1b, UD3a, and UD3b) and the placement of the cervix upon it (PCO), the posterior uterine dimension on the inferior half of the uterus (UD3b) and the perpendicular cervical offset (PCO) had the greatest effect on stretch in the proximal cervix mid-stroma (Figs. 8 and 14, Appendix C Figures 13–16, Appendix C Table 2). In the radial direction, increasing UD3b from 32.97–49.45 mm produced changes in stretch in the anterior, left, and posterior mid-stroma (−0.03, 0.01, and −0.02, respectively), and changes in stretch were also found with PCO from 11.00–33.01 mm (−0.03, 0.00, and 0.03, respectively, not monotonic in the anterior mid-stroma with increasing PCO). In the circumferential direction, notable changes to stretch in the left mid-stroma were produced with UD3b from 32.97–49.45 mm (−0.03) and PCO from 11.00–33.01 mm (0.02, not monotonic with increasing PCO). Only marginal changes to stretch in the normal direction were found for all posterior uterine dimensions (change in stretch was *<*0.01). Mid-stromal changes in stretch are reflected in the heat maps of stretch with varied UD3b (Fig. 14). Radial stretch decreased in the anterior and posterior proximal cervix with increasing UD3b, with marginal increases in the left and right proximal cervix. Circumferential stretch decreased in the left and right proximal cervix with increasing UD3b, with marginal increases in stretch in the anterior region. Marginal changes to normal stretch in the proximal cervix were observed with increasing UD3b.

**Fig. 14.**
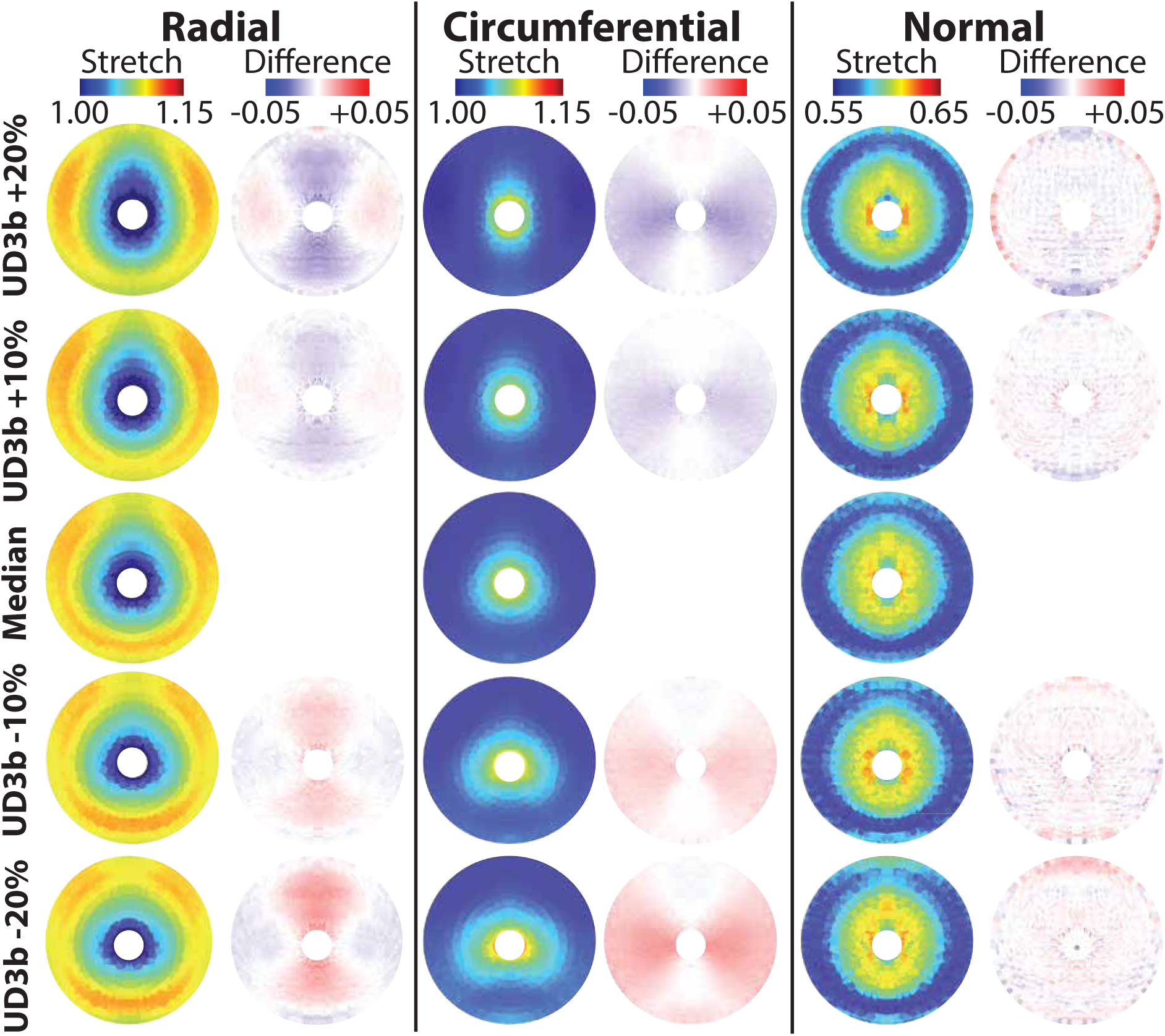
Increasing UD3b produced an overall decrease in radial and circumferential stretch in the proximal cervix. Heat maps of proximal cervix face stretch and difference from median in the surface radial, circum-ferential, and normal direction for varied posterior uterine diameter on the inferior half of the uterus (U3b).\

**Fig. 15.**
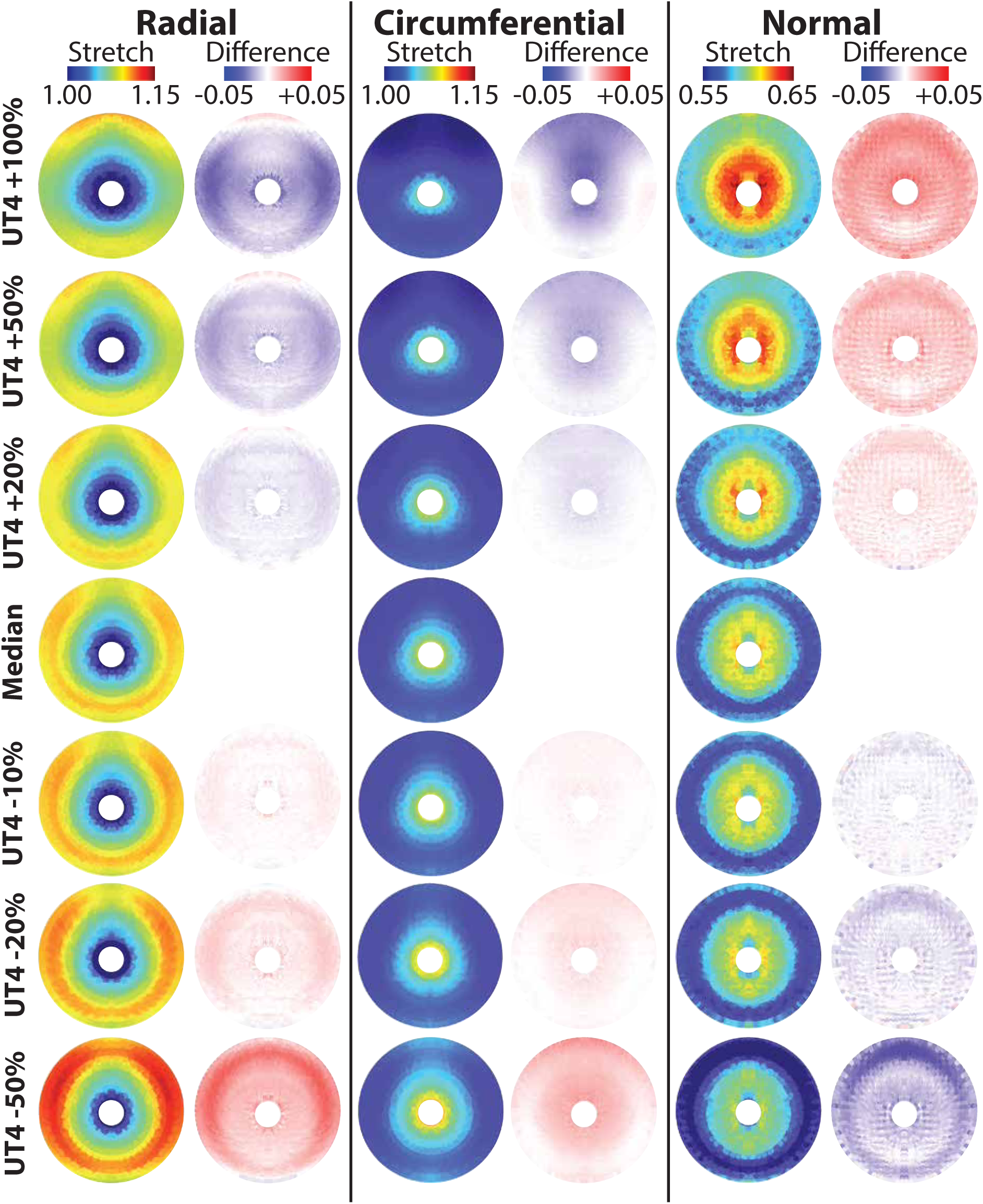
Increasing UT4 produced an overall decrease in stretch in the proximal cervix. Heat maps of proximal cervix face stretch and difference from median in the surface radial, circumferential, and normal direction for varied lower uterine segment wall thickness (UT4).

**Fig. 16.**
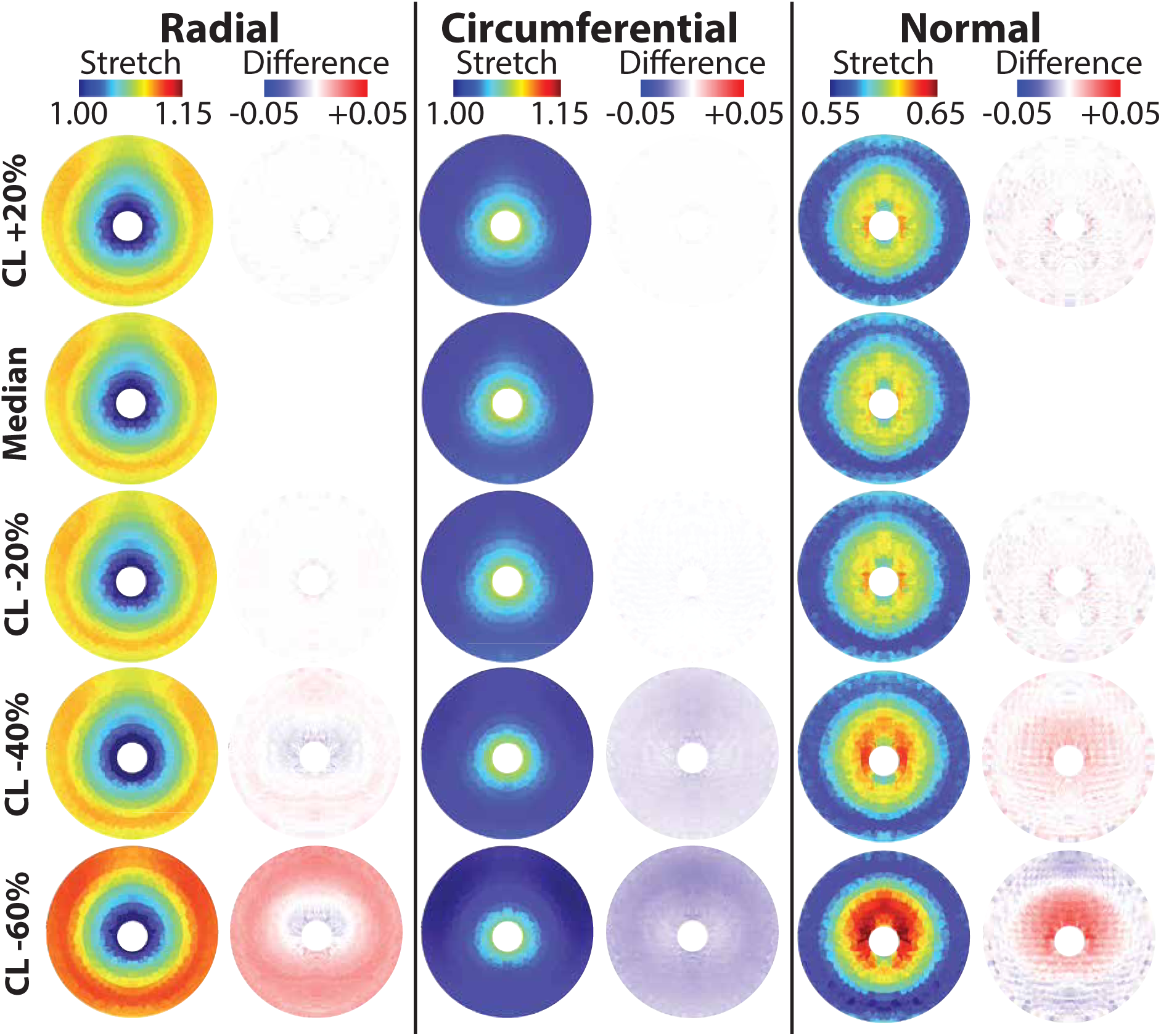
Stretch in the proximal cervix was sensitive to CL. *<* **25 mm.** Heat maps of proximal cervix stretch and difference from median in the surface radial, circumferential, and normal direction for varied cervical length (CL).

PCO produced marginally greater changes to maximum *λ*_1_ in the cervical IO than UD3b, the posterior uterine dimension on the superior half of the uterus (UD3a), and their locator dimensions along UD1 (UD1b and UD1a, respectively) (Fig. 11, Appendix C Table 2). Increasing PCO from 11.00–33.01 mm produced a change of magnitude 0.02 in maximum *λ*_1_ in the cervical IO (not monotonic with increasing PCO). A magnitude change of 0.01 in maximum *λ*_1_ in the cervical IO was found with increasing UD3a from 26.66–39.98 mm, UD1a from 81.47–99.57 mm, UD3b from 32.97–49.45 mm, and UD1b from 24.14–36.21 mm (not monotonic with increasing UD3a, UD1a, UD3b, or UD1b).

#### 3.2.3 Tissue stretch in the proximal cervix is sensitive to uterine wall thicknesses

The fundal (UT1), anterior (UT2), left/right (UT3), and lower uterine segment (UT4) uterine wall thicknesses produced notable effect on stretch in the mid-stroma of the proximal cervix, with the greatest magnitude change typically observed for UT4 (Figs. 9 and 15, Appendix C Figures 17–19, Appendix C Table 2). In the radial direction, increasing UT4 from 4.25–17.01 mm produced a decrease in stretch in the anterior, left, and posterior mid-stroma (−0.02, −0.04, and −0.02). Increasing UT1 from 5.18–7.78 mm and increasing UT3 from 5.84–8.77 mm produced an increase of 0.01 in radial stretch in the left mid-stroma, and increasing UT2 from 4.61–6.91 mm produced an increase of 0.01 in radial stretch in the anterior and posterior mid-stroma. In the circumferential direction, increasing UT4 from 4.25–17.01 mm produced a decrease in stretch in the anterior, left, and posterior mid-stroma (−0.04, −0.01, and −0.02, respectively). Increasing UT1 from 5.18–7.78 mm and increasing UT3 from 5.84–8.77 mm produced an increase of 0.01 in circumferential stretch in the anterior and posterior mid-stroma, and increasing UT2 from 4.61–6.91 mm produced an increase of 0.01 in circumferential stretch in the left mid-stroma. In the normal direction, increasing UT4 from 4.25–17.01 mm produced an increase in normal stretch (less compression) in the anterior, left, and posterior mid-stroma (0.03, 0.03, and 0.01, respectively), and increasing UT3 from 5.84–8.77 mm decreased stretch (more compression) in these regions (−0.01 in anterior, left, and posterior mid-stroma). Increasing UT2 from 4.61–6.91 mm decreased stretch by −0.01 in both the anterior and posterior mid-stroma. Mid-stromal changes in stretch are reflected in the heat maps of stretch with varied UT4 (Fig. 15). Radial stretch decreased with increasing UT4, particularly towards the uterocervical junction. Circumferential stretch decreased with increasing UT4, particularly in the anterior proximal cervix. Normal stretch increased (less compression) in the proximal cervix with increasing UT4, particularly towards the anterior uterocervical junction.

UT4 produced the greatest change to maximum *λ*_1_ in the cervical IO of the uterine wall thicknesses (Fig. 11, Appendix C Table 2). Increasing UT4 from 4.25–17.01 mm produced a decrease of 0.05 in maximum *λ*_1_ in the cervical IO. An increase of 0.01 in maximum *λ*_1_ in the cervical IO was produced with increasing UT1 from 5.18–7.78 mm, UT2 from 4.61–6.91 mm, and UT3 from 5.84–8.77 mm.

#### 3.2.4 Tissue stretch in the proximal cervix is affected by cervical dimensions

The cervical length (CL) had more of an effect on stretch in the mid-stroma of the proximal cervix than outer cervical diameter (CD1), cervical canal diameter (CD2), and anterior uterocervical angle (AUCA) (Figs. 10 and 16, Appendix Figures 20–23, Appendix C Table 2). In the radial direction, increasing CL from 12.92–38.77 mm produced decreases in stretch in the anterior, left, and posterior mid-stroma (−0.01, −0.01, and −0.02, respectively). In the circumferential direction, increasing CL from 12.92–38.77 mm produced increases in stretch in the anterior, left, and posterior mid-stroma (0.02, 0.01, and 0.01, respectively), and increasing AUCA from AUCA from 64.93– 97.39◦ decreased circumferential stretch in the left mid-stroma by −0.01. In the normal direction, increasing CL from 12.92–38.77 mm produced decreases of −0.01 in stretch (more compression) in the anterior and left mid-stroma, and increasing AUCA from 64.93–97.39◦ produced changes in normal stretch in the anterior and posterior mid-stroma (−0.01 and 0.01, respectively, not monotonic with increasing AUCA). Mid-stromal changes in stretch are reflected in the heat maps of stretch with varied CL (Fig. 16). Radial stretch in the proximal cervix increased in the mid-stroma to uterocervical junction for the shortest CLs, with a slight decrease in radial stretch towards the cervical IO. Circumferential stretch decreased in the proximal cervix for the shortest CLs. Normal stretch in the proximal cervix increased (less compression) towards the cervical IO for the shortest CLs, with a slight decrease in normal stretch (more compression) towards the uterocervical junction.

Similar changes to maximum *λ*_1_ in the cervical IO were found with varying the cervical dimensions (Fig. 11 and Appendix C Table 2). A magnitude change of 0.03 in maximum *λ*_1_ in the cervical IO was found with increasing CL from 12.92–38.77 mm, CD2 from 1.56–4.67 mm, and AUCA from 64.93–97.39◦ (not monotonic with increasing CL, CD2, or AUCA). Increasing CD1 from 30.07–36.77 mm produced an increase of 0.02 in maximum *λ*_1_ in the cervical IO (not monotonic with increasing CD1).

#### 3.2.5 Tissue stretch in the proximal cervix is sensitive to fetal membrane thickness

The FM thickness affected stretch in the mid-stroma of the proximal cervix, with similar changes to stretch observed in the anterior, left, and posterior mid-stroma (Fig 12a–c, Appendix C Figure 24, Appendix C Table 2). In the radial direction, increasing FM thickness from 0.25–2 mm decreased radial stretch in the anterior, left, and posterior mid-stroma (−0.02, −0.01, and −0.02, respectively). In the circumferential direction, increasing FM thickness from 0.25–2 mm produced an overall change in stretch in the anterior, left, and posterior mid-stroma (0.01 in all regions, not monotonic with increasing FM thickness). In the normal direction, increasing FM thickness from 0.25–2 mm produced an overall increase in stretch (less compression) in the anterior, left, and posterior mid-stroma (0.07, 0.07, and 0.08, respectively). Varying FM thickness also caused changes to maximum *λ*_1_ in the cervical IO (Fig. 12d). A decrease of −0.11 in maximum *λ*_1_ in the cervical IO was produced with FM thickness increased from 0.25–2 mm.

## 4 DISCUSSION

We built a digital twin simulation of human pregnancy, based on 2D maternal sonography and fiber-based soft-tissue material properties, to quantify the magnitude and pattern of cervical distensibility to intrauterine pressure at the time of second trimester cervical length screening. The purpose of this digital twin finite element model is two-fold. First, the simulations provide an *in silico* method to structurally interrogate the biomechanical function of the cervix, given its anatomical shape/size, tissue material properties, and boundary conditions. The proximal cervical stretch maps provide a functional readout of *in vivo* cervical deformation, with larger stretches reflecting a more compliant structure. Second, these structural evaluations provide the basis for developing a PTB risk diagnostic tool beyond cervical length alone, where the digital twin integrates multiple anatomical and tissue features. Beyond this study, the digital twin has the capacity to integrate other pertinent features, such as bodily biofluid markers and intrauterine contents (placenta, amniotic fluid, and fetus). Here, we provide foundational finite element modeling methods and input sensitivity analysis to develop a functional measurement of cervical structural performance at the time of routine second trimester anatomy survey in pregnancy.

Finite element simulations show that the Young’s modulus (*E*) of the cervical ground substance and the fiber stiffness (*ξ*) drove tissue stretch magnitudes in the cervical internal os (IO) and in the proximal cervix face. Maternal sonographic dimensions drove stretch patterns in the proximal cervix under simulated intrauterine pressurization. We found these maternal loading patterns through our updated approach to generating patient-specific parametric computational models of the pregnant maternal anatomy, building on the work of Westervelt et al. [19]. We used a previously described design table driven approach to generate solid models of the uterus and cervix from sonographic dimension measurements, adding the FM, a surrounding abdomen, and a uterocervical transition zone [52, 20]. Material models based on existing equilibrium mechanical test data of the uterus and cervix were chosen to mimic the physiological tissue behavior, and appropriate second trimester material parameter values were determined [21, 22]. The developed approach serves as a robust *in silico* mechanical test of second trimester maternal anatomy, where the deformation response of the FM and cervix can be quantified under an applied intrauterine pressure.

The computational model developed in this work is intended for implementation in the second trimester. During gestation, the cervix softens most in the first trimester, as observed by clinical physical examination, elastography, and tissue aspiration, which is partially attributed to the remodeling of the cervical collagen fibers [69, 23, 70, 71, 20]. We hypothesize collagen fiber remodeling to be captured by the cervical fiber stiffness modulus (*ξ*) in the material model. Of the uterine and cervical material properties, stretch in the cervical IO and the proximal cervix mid-stroma were most sensitive to changes in the cervical stiffness moduli (*E* and *ξ*). The cervical ground substance stiffness (*E*) had the largest overall effect on resultant stretch. The greatest change in stretch in the proximal cervix mid-stroma occurred in the normal direction across the range of tested values for cervical *E*, with dramatically less compression in the proximal cervix face with increasing cervical *E*. The reduction in cervical compression with increased cervical *E* in turn caused the reduction in the tensile radial and circumferential stretch. After cervical *E*, cervical fiber stillness modulus (*ξ*) had the largest effect on radial and circumferential stretch in the proximal cervix mid-stroma and maximum *λ*_1_ in the cervical IO. The minimal effect of cervical *ξ* on surface normal stretch in the proximal cervix likely stems from the fibers’ ability to bear only tensile loads. Westervelt et al. also investigated the effect of cervical *ξ* on resultant cervical deformation to applied intrauterine pressure, quantifying the percentage of elements above an effective stretch threshold of 1.05, with a nonlinear decrease from 93% of cervical elements above this threshold for cervical *ξ* = 1.71 kPa and 25% of cervical elements above this threshold for cervical *ξ* = 769 kPa [19].

Radial stretch in the proximal mid-stroma and maximum *λ*_1_ in the cervical IO decreased rapidly as cervical *ξ* increased from 1–50 kPa (0 to 25% normalized material property value), with minor changes to stretch as cervical *ξ* increased beyond this point (Fig. 4 and 6). Given that cervical stiffness rapidly decreases in the first to second trimesters, it is likely that patient-specific cervical *ξ* would exist in the range where the model is most sensitive (1–50 kPa) [69, 20]. On the other hand, it is not clear that cervical *E* is affected by gestational age. Where cervical *ξ* from mechanical testing IFEA was significantly different between the term pregnant (PG) and nonpregnant (NP) tissue (*p* = 0.03), cervical *E* only trended towards significance (*p* = 0.08, Appendix A Figure 4). Other works have also found the stiffness of the cervical ground substance at equilibrium to be similar between PG and NP cervical tissue. In paired uniaxial tension and spherical indentation tests of PG and NP cervical tissue from humans, the bulk modulus *κ*, which is hypothesized to be related to the non-fibrous cervical components, was not significantly different between PG and NP cervical tissue samples from comparable cervical locations [22]. In non-human primate cervical tissue, fitted cervical *E* from across gestation was also not significantly different between gestational time points [41]. Therefore, cervical *E* may not significantly change during gestation, though it may vary between individuals. Given the importance of cervical *ξ* and *E* to resultant stretch in the proximal cervix, ideally both of these material parameters would be included as patient-specific inputs in future studies using the presented modeling pipeline in cohort studies of pregnancy biomechanics.

Varying uterine material properties produced some minimal changes to tissue stretch in the proximal cervix, though not to the extent of cervical *E* and *ξ*. The uterine parameters that produced the most change to proximal cervix mid-stroma and cervical IO stretch were uterine *E* and the ground substance Poisson’s ratio (*ν*). The stretch in the proximal cervix mid-stroma was more sensitive to uterine *ν* than any other uterine material parameter, whereas maximum *λ*_1_ in the cervical IO was marginally more sensitive to uterine *E*. In fitting the material models to existing equilibrium mechanical tests of human uterine tissues via IFEA, we found a significant difference in uterine *E* for NP and PG samples (*p* = 0.03) and no significant difference was found for uterine *ν* (*p* = 0.52, Appendix A Figure 3). Previous work fitting a similar material model to spherical indentation tests on the same uterine samples used in this work found no significant difference in uterine *E* and *ν* in NP and PG uterine tissue samples at equilibrium [24]. Microindentation tests on myometrial tissue from NP and PG human uteri [40] and across gestation in the non-human primate uterus [72] also showed no significant difference in uterine *E*. Therefore, we do not believe patient-specific values for uterine *E* and *ν* are necessary in future implementation of this modeling pipeline to investigate equilibrium cervical stretch. However, future studies that investigate cervical stretch under a larger applied intrauterine pressure or with increasing gestational age would likely be affected by uterine material properties.

Considered together, the effects of varied uterine and cervical material properties on stretch in the proximal cervix affected stretch magnitude rather than stretch pattern (Appendix C Figures 2–9). A notable exception to this observation is cervical *ξ*, where radial stretch magnitude increased most near the uterocervical junction and less towards the cervical IO for decreasing cervical *ξ* (Appendix C Figure 4). Still, there was radial symmetry about the cervical canal in the difference heat maps of the proximal cervix with varying cervical *ξ*. When maternal sonographic dimensions were varied, the magnitude of stretch difference was of a smaller magnitude than when cervical *E* or cervical *ξ* were varied, but there were often observed differences in stretch pattern.

Most intrauterine diameters (UD1, UD2, and UD3) did not greatly alter stretch in the proximal cervix, with the exception of the left-right intrauterine diameter (UD4). UD4 controls the left-right width of the intrauterine cavity, and the changes observed in stretch in the proximal cervix were consistent with changes in the left-right geometry. The radial stretch decreased in the left (and by symmetry right) proximal cervix as UD4 increased, and circumferential stretch in the anterior and posterior proximal cervix decreased with increased UD4 (Fig. 13). Similarly, as UD4 increased the circumferential stretch decreased in the anterior and posterior proximal cervix, where circumferential stretch is more aligned with the left-right direction. In the normal direction, decreased compressive deformation was observed in the proximal cervix face as UD4 increased. Taking the radial, circumferential, and normal stretches together, there was less deformation in the proximal cervix face as UD4 increased, particularly in stretch aligned in the left-right direction. Changes to UD4 affect the coronal curvature of the uterine wall near the proximal cervix, which has the effect of focusing the intrauterine load on the proximal cervix as UD4 narrows and dispersing load from the proximal cervix as UD4 widens.

The curvature of the posterior uterine wall in the sagittal plane and its positioning relative to the cervix are defined by the posterior uterine dimensions (UD1a, UD1b, UD3a, UD3b, and PCO). The posterior uterine dimensions that define the uterine wall near the proximal cervix (UD3b and PCO) had the greatest effect on stretch in the proximal cervix. This is exemplified by surface radial and circumferential stretch in the proximal cervix face, which varied drastically with changes to UD3b. Similarly to UD4, UD3b exhibited regional trends in proximal cervix stretch change, with radial stretch increasing in the anterior and posterior and circumferential stretch increasing in the left (and by symmetry right) proximal cervix as UD3b increased. These changes to stretch are a consequence of the geometry which UD3b alters, where a larger UD3b creates less drastic curvature around the proximal cervix face in the anterior-posterior direction.

Previously, Westervelt et al. found larger PCO values to be associated with greater levels of stretch in the cervix [19]. The results from this study do not directly dispute these findings, but they do reveal a more nuanced effect, where the region of maximum radial and circumferential stretch in the proximal cervix shifts with varying PCO. Regarding PCO and UD3b in the context of future clinical and computational studies, it was previously shown that both UD3b and PCO have low agreement between observers [52]. Because they are on the posterior side of the uterus, sonographic waves must travel farther from the anteriorly placed probe and through more tissues to reach the posterior uterine wall, making them more difficult to visualize. Additionally, both PCO and UD3b depend on the initial placement of UD1, which may vary between observers. Given the importance of the curvature of the uterus near the cervix on stretch in the proximal cervix, future studies should explore approaches to better capture uterine wall curvature in this region.

Uterine wall thicknesses (UT1, UT2, UT3, and UT4) had varied effects on stretch in the proximal cervix. Deformation in the proximal cervix face and in the cervical IO increased with increasing uterine wall thickness for UT1, UT2, and UT3, but decreased with decreasing UT4. UT4 is measured in the lower uterine segment anterior to the proximal cervix, with increased UT4 resulting in an increased volume of tissue near the proximal cervix. With thicker UT4s, the deformation of the applied intrauterine pressure is felt across larger volume of tissue in the lower uterine segment resulting in overall less deformation of the tissue near the intrauterine surface. With thinner UT4s, the intrauterine pressure causes deformation of the relatively soft proximal cervix rather than the relatively stiff surrounding abdomen, particularly near the anterior uterocervical junction, resulting in larger levels of deformation at the intrauterine surface. Practical differences also exist between UT1–3 and UT4 outside of location. UT1–3 have poor reproducibility between observers, whereas UT4 has excellent reproducibility and more variability between individuals [52]. UT4 has been shown to decrease significantly across gestation, with marked decreases occurring between the late second and middle third trimesters [52, 20]. Yapan et al. have reported lower uterine segment thickness to be associated with PTB risk, with thinner lower uterine segment thicknesses associated with higher PTB risk [73]. From the results of this sensitivity study, we hypothesize that the increased stretch in the proximal cervix due to the thinning of UT4 generates a biomechanical risk of PTB, but further study is needed to confirm this hypothesis.

Of all dimensions investigated in this study, maximum *λ*_1_ in the cervical IO was most sensitive to FM thickness, as well as normal stretch in the proximal cervix. However, varied FM thickness did not produce radial and circumferential stretch changes of greater magnitude than other sonographic dimensions, such as PCO, UD3b, and UT4. Unlike the uterine and cervical dimensions investigated in this study, FM thickness is not a dimension we have collected in our previous longitudinal studies, and FM thickness may not be well captured by the resolution available on most sonography machines [52, 20, 74]. Therefore, FM thickness is a dimension that will likely be given an assumed value in future work rather than measured as a patient-specific value. The findings from this sensitivity study show that while stretch magnitude in the cervical IO is highly sensitive to FM thickness, the magnitude and pattern of stretch in the proximal cervix is not. The material model used for the FM was fit to uniaxial mechanical tests of the at-term amnion, which has been reported to have a thickness *<*0.1 mm [49, 27, 74]. Westervelt et al. used an FM thickness of 0.1 mm in their initial implementation of this parametric model of pregnancy, with an Ogden material model fit to biaxial tests of the at-term amnion [19, 49]. Later, the fiber composite model was used to capture the bending behavior of the FM, and a 1 mm FM thickness was assumed in IFEA for fitting the material model and in the maternal geometry [27]. In this work, we chose an FM thickness that is representative of the second trimester combined amnion and chorion (0.79 mm), and we attempted to vary it from values representative of the at-term amnion (0.1 mm) to the largest reported second trimester combined amnion chorion thickness ( 2 mm) [56, 74, 68]. We found FM thicknesses smaller than 0.79 mm to have convergence issues, as was found by Westervelt, and could not get full convergence for the 0.1 mm FM thickness model [27]. Ultimately, the amnion thickness and FM material properties in the second trimester remain unknown, and currently there are no *in vivo* approaches to reliably measure these parameters. Therefore, though the FM thickness and material parameter values used in this model may not be representative of the second trimester maternal anatomy, we will acknowledge these as modeling assumptions that are used in future models of maternal anatomy.

Short cervical length (CL) is a known risk factor for PTB, with cervices less than 25 mm considered at increased risk for PTB [4]. CL is currently the only imaging biomarker used to for PTB prediction, but it has a reported positive predictive value ranging from 6–76%, depending on patient risk factors [75]. In this computational study, we observed elevated radial stretch in the proximal cervix when CL was below 20% of the median (*<*25.85mm). Thus, short CL may result in increased radial stretch in the proximal cervix *in vivo*, which itself may be a precursor to premature cervical dilation. Similarly, Westervelt et al. found a decrease in proximal cervix stretch with increased CL, though CL*<*25 mm was not included in the study [19]. In addition to CL, AUCA has also been studied as a possible predictor of PTB, with no consensus on its advantage over or in combination with CL [76, 77, 78, 79, 80, 81, 82, 83]. For studies that show significant differences in AUCA between term and preterm births, larger AUCAs are associated with PTB [76, 79, 83]. In this study, increasing AUCA produced an increase in radial stretch in the anterior mid-stroma and decrease in radial stretch in the posterior mid-stroma. However, increasing AUCA produced a decrease in radial stretch in the anterior proximal cervix at the uterocervical junction. This is similar to what was found for CL and UT4, where radial stretch in the proximal cervix mid-stroma to uterocervical junction (particularly in the anterior region) increased with decreasing CL and UT4. These findings indicate that the biomechanical consequence of these risk factors (short CL, thin UT4, and large AUCA) is an increase in tension between the anterior cervix and uterus. In addition, the increase in tensile stretch between the cervix and uterus elicited by a short CL, thin UT4, and large AUCA supports a combination of these dimensions as a clinically useful biomarker of PTB. Westervelt et al. also found minor differences in stretch with varied AUCA [19]. These findings highlight the importance of maternal sonographic anatomical measurements that define the region surrounding the proximal cervix in the resulting pattern of cervical deformation.

### 4.1 Limitations

Though the methods and results in this work on modeling patient-specific maternal anatomy in the second trimester present many future research opportunities, they are not without assumptions and limitations. One limitation is the basis of the models, which were built using measurements from already pressurized pregnant anatomy. Thus, the geometry used in simulation was not actually in a reference state when the load was applied. The results of these models can therefore only be considered an *in silico* mechanical test of maternal anatomy, and the resulting stretch magnitude is not representative of *in vivo* loading. Future work should consider the growth and remodeling of these tissues to find more physiologically relevant deformation levels in the cervix. In future studies, we plan to utilize this computational approach to model maternal anatomy in the early to mid-second trimester. Intrauterine pressure (IUP) in the early to mid-second trimester, or 14–24 weeks gestation, was reported to be 0.3–1.3 kPa by Fisk et al. [64]. We chose to use the IUP in Equation 9 as it had been used in a previous parametric model of pregnancy [19, 64], though Sideris and Nicolaides found an IUP of 1.5 kPa at 16 weeks, with a range of 0.7–2.1 kPa from 14–24 weeks gestation [84]. It is probable that proximal cervix stretch is highly sensitive to IUP, but because the measurement of intrauterine pressure is an invasive procedure, it is not expected that IUP can feasibly be a patient-specific parameter. Therefore, a constant intrauterine pressure of 0.7 kPa was used in all simulations. Additionally, the material property values in the cervix and uterus were based on averaging equilibrium mechanical test results from non-pregnant and at-term tissues, and not on actual second-trimester tissues. Though this approach is supported by mechanical data from non-human primate cervical tissue across gestation, given that cervical *E* and *ξ* have the greatest effect on resulting stretch, it is imperative to have patient-specific measures of cervical stiffness in future patient-specific efforts.

Material models were chosen to represent the most salient features of uterine and cervical equilibrium mechanical response (tension/compression asymmetry and fibers that could orient to the direction of greatest tensile stretch). A spherical fiber distribution was assigned in both the uterus and cervix, but it is well established that both have sophisticated fiber structures that evolve throughout pregnancy. The uterus is composed primarily of densely packed smooth muscle cell fascicles, which are sheathed in a collagenous extracellular matrix, where smooth muscle fascicles and collagen fibers run parallel to each other [5, 34, 24]. The cervix is primarily composed of preferentially aligned collagen fibers. The “three-zone” theory describes the cervical collagen orientation, with an inner and outer zone of longitudinally aligned collagen fibers sandwiching a middle zone, with circumferentially aligned fibers, with considerable dispersion about these primary direction [36, 8]. Cervical fiber direction and dispersion were previously explored, but due to the high levels of fiber dispersion in the uterus and cervix and the inability to determine fiber dispersion on a patient-specific basis, a spherical fiber distribution was ultimately used, allowing fiber realignment to the direction of greatest simulated deformation [50]. Previous research has also found that the remodeling of the collagen fibers themselves has a greater impact on cervical mechanical response than fiber dispersion, supporting our decision to use a spherical fiber distribution [23].

A simplistic material and geometry approach was used to model the FM. The FM is a bi-layer tissue composed of two layers, the amnion and the chorion [85]. The chorion is significantly thicker than the amnion, with term thicknesses of 0.431±0.113 mm and 0.111±0.078 mm, respectively [85]. In our solid FM model, we used a monolayer structure for the FM with a total thickness representative of the FM in the second trimester (0.79mm) [56]. The chorion and amnion also have distinct microstructures, with the chorion being highly cellular and the amnion being highly collagenous [86]. More detailed models of the FM have been developed, including distinct FM layers in solid models with specified material models for each layers [47, 26]. We modeled the FM material as a neo-Hookean ground substance with an embedded fiber distribution, which was fit to mechanical tests of the at-term amnion [27, 49]. Because the goal of this work is to develop a clinically implementable patient-specific *in-silico* mechanical test, these simplifications are justified. However, future work to develop true digital twins of pregnancy should consider higher fidelity modeling techniques for the FM material and geometry. There is also ambiguity in defining where cervical tissue ends and uterine tissue begins, with the transition zone between them referred to as the isthmus in nonpregnancy, which is pulled open by the stretched uterus during pregnancy to become the lower uterine segment [87]. The actual location and geometry of the uterocervical transition zone are likely dependent on both individual variation and gestational age, and the effects of the uterocervical transition zone geometry should be explored in future study.

The study design used in the sensitivity studies was a one-at-a-time approach, which allows for an initial understanding of the sensitivity of the model stretch results. However, it does not allow for a quantification of expected error in future patient-specific models which would have uncertainties in all material and sonographic values. Thus, future work involving a probabilistic approach to quantifying sensitivity must be undertaken. Also, the ranges used to vary the ultrasonic dimensions were not constant between dimensions, so comparing stretch ranges does not provide an equal comparison approach. However, the ranges chosen for each maternal sonographic dimension are reflective of the ranges we’ve observed for variation between individuals and measurement error between observers, making them relevant for understanding relative importance for implementation in future studies. Finally, though useful for gaining an understanding of how deformation in the proximal cervix changes with varying parameters, the approach of comparing the stretch change ranges and linear stretch slopes in the defined ROIs only provide a rough measure of sensitivity, as many parameters did not exhibit a linear response and it is unknown if these regions are important to the prediction of PTB. That being said, this work does provide a holistic starting point for future work, as well as a clinically implementable method to be used in future patient-specific studies.

## 5 CONCLUSION

This work presents a method to generate patient-specific finite element models of pregnant maternal anatomy in the early second trimester. The potential cross-talk between humans and computational models will accelerate PTB investigation, which could meaningfully inform clinical management. We investigated the sensitivity of the resultant model stretch to various parameters, including material property and sonographic maternal dimensions. Overall, tissue material properties primarily affected the magnitude of proximal cervix stretch, whereas sonographic maternal dimensions primarily affected stretch pattern. The cervical ground substance Young’s modulus and fiber stiffness modulus had the greatest effect on stretch in the proximal cervix when compared to other material properties and all of the sonographic dimension measurements. The sonographic dimension measurements that control the region surrounding the proximal cervix had the greatest effect on its deformation (e.g., UD3b, PCO, UT4). The research presented here is a necessary step towards future digital twin efforts, which will further our understanding of pregnancy biomechanics to better predict and prevent PTB.

## ACKNOWLEDGMENT

The authors would like to thank Jada Hinds, Miccaela Lejwa, Arielle Feder, and Colin Edwards for their assistance in the early stages of this study. We acknowledge computing resources from Columbia University’s Shared Research Computing Facility project, which is supported by NIH Research Facility Improvement Grant 1G20RR030893-01, and associated funds from the New York State Empire State Development, Division of Science Technology and Innovation (NYSTAR) Contract C090171, both awarded April 15, 2010

## FUNDING

Jada Hinds and Miccaela Lejwa were supported by the Columbia-Amazon Summer Undergraduate Research Experience program. This work is sponsored by the Eunice Kennedy Shriver Institute of Child Health & Human Development under Award R01HD091153 to KMM. The content is solely the responsibility of the authors and does not necessarily represent the official views of the National Institutes of Health.

## Notes

### Competing Interest Statement

We disclose that Dr. Myers has industry-sponsored research with GE Healthcare to develop ultrasound-based digital twins for preterm birth risk assessment and CX Therapeutics. Dr. House is the president of CX Therapeutics, a company developing cervical closure devices for improved PTB therapies.

