## Appendix A for "A finite element model of pregnancy derived from maternal sonography: effect of uterine and cervical structural properties on cervical mechanical loading"

### Appendix A: Uterine and cervical material property values from inverse finite element analysis

**4 Conclusion on material properties** **5**

### 1 METHODS

#### 1.1 Mechanical Testing Data

##### *1.1.1 Uterus*

The uterine mechanical testing data used in this work were collected and analyzed for previously published work, from which a subset was included in this analysis, specifically those analyzed in both Fang et al. 2021 and 2025 [1, 2]. Fundal human uterine tissue specimens were collected from seven non-pregnant and seven pregnant patients. The non-pregnant patients were premenopausal ($<$50 years old) and undergoing total hysterectomy, and the pregnant patients were undergoing a cesarean section hysterectomy in late pregnancy. Two samples of the fundal myometrium were mechanically tested from each patient, giving 14 non-pregnant and 14 pregnant samples. Each sample was mechanically tested in compression via spherical indentation and in tension. In the indentation test, a ramp-hold protocol was followed at displacements of 15%, 30%, and 45% of the specimen thickness. In the tension test, a ramp-hold protocol was followed at displacements of 15%, 30%, and 45% of the gauge-to-gauge distance. Force-displacement-time data was recorded, along with measurements of sample size to evaluate the first and second principal Lagrange strains. Full details on tissue collection, preparation, and mechanical testing can be found in Fang et al. 2021 and 2025 [1, 2].

##### *1.1.2 Cervix*

Existing mechanical testing data of the cervix were used in this work [3]. Human cervical tissue specimens were collected from four non-pregnant and four pregnant patients. The non-pregnant patients were premenopausal ($<$50 years old) and undergoing total hysterectomy. The pregnant patients were undergoing cesarean hysterectomy. Slices of cervical tissue were collected at the internal and external os, giving a total of 8 non-pregnant and 8 pregnant samples. Each sample was first tested under indentation, during which the alignment of cervical collagen fibers was deduced. Two strips of tissue were then cut from the sample: one perpendicular and one parallel to the fiber direction. In the indentation test, a ramp-hold protocol was followed at displacements of 12.5%, 25%, 37.5%, and 50% of the specimen thickness. In the tension test, a several-stage ramp-hold was conducted, with maximum tensile strains chosen based on previous experimental results [3, 4]. Full details on tissue collection, preparation, and mechanical testing can be found in [3]. In this work, only the indentation and parallel-to-fiber tensile tests were used.

#### 1.2 Material Model

A phenomenological material modeling approach was pursued for computational efficiency purposes. Uterine and cervical tissue were modeled as a continuously distributed fiber composite embedded in a neo-Hookean ground substance. This material model was chosen because it allows for the reorientation and alignment of the fibers in the principal stretch direction and captures uterine and cervical tissues' asymmetric and non-linear tension-compression response well.

#### 1.3 Inverse Finite Element Analysis

##### *1.3.1 Uterus*

Inverse finite element analysis on the uterine mechanical testing data was performed in FEBio Studio v1.3.0 with the approach established by Shuyang Fang [5, 6]. A customized MATLAB (MathWorks, Natick, MA) code implemented an established genetic-based algorithm to test and assess material property value fits while automatically generating the sample geometry (Fig. 1)

[7]. Using the force-strain data, the best-fit parameter values for Young's modulus ($E$), Poisson's ratio ($\nu$), fiber stiffness modulus ($\xi$), and coefficient of the exponential argument ($\alpha$) were found by minimizing the error given by equation 1 [6]:

$$\text{error} = \frac{1}{8}\sum_{i=1}^{n}\left|\frac{e_1^{FEA^i} - e_1^{EXP^i}}{e_{norm}^{EXP^n}}\right| + \frac{1}{8}\sum_{i=1}^{n}\left|\frac{e_2^{FEA^i} - e_2^{EXP^i}}{e_{norm}^{EXP^n}}\right| + \frac{1}{4}\sum_{i=1}^{n}\left|\frac{F_I^{FEA^i} - F_I^{EXP^i}}{F_I^{EXP^n}}\right| + \frac{1}{2}\sum_{i=1}^{n}\left|\frac{F_T^{FEA^i} - F_T^{EXP^i}}{F_T^{EXP^n}}\right| \quad (1)$$

where $i$ is the $i^{th}$ ramp-hold level, $n$ is the number of total ramp-holds, $FEA$ is the force/strain from finite element analysis, $EXP$ is the force/strain from the mechanical test, $F_I$ is indentation force, $F_T$ is tension force, $e$ is the principal strain of the sample during indentation, and $e_{norm}^{EXP^n}$ is the Euclidean norm of $e_1^{EXP^n}$ and $e_2^{EXP^n}$. A Welch's t-test was performed between the non-pregnant and pregnant parameter results.

###### *1.3.2 Cervix*

Inverse finite element analysis on the cervical mechanical testing data was performed in FEBio Studio v1.3.0 with the approach established by Shi et al. [5, 3]. A customized MATLAB (MathWorks, Natick, MA) code implemented an established genetic-based algorithm to test and assess material property value fits while automatically generating the sample geometry (Fig. 2) [7]. Using the force-strain data, the best-fit parameter values for Young's modulus ($E$), Poisson's ratio ($\nu$), fiber stiffness modulus ($\xi$), and coefficient of the exponential argument ($\alpha$) were found by minimizing the error given by equation 2:

$$\text{error} = \sum_{i=1}^{n}\left|\frac{F^{FEA^i} - F^{EXP^i}}{F^{EXP^n}}\right| \quad (2)$$

where $i$ is the $i^{th}$ ramp-hold level, $n$ is the number of total ramp-holds, $FEA$ is the force/strain from finite element analysis, $EXP$ is the force/strain from the mechanical test, and $F$ is force from the indentation/tension test. A Welch's t-test was performed between the non-pregnant and pregnant parameter results.

#### 2 RESULTS

##### 2.1 Uterus

The best-fit combination of parameters for all uterine samples is given in table 1, with the average values and standard deviations in the non-pregnant and pregnant state shown. Boxplots of best-fit uterine parameters in the non-pregnant and pregnant states are shown in fig. 3, along with *P*-values from Welch's t-tests. Young's modulus ($E$) was significantly smaller in the non-pregnant uterine samples compared to the pregnant uterine samples (1.13 ± 0.68 kPa vs. 1.76 ± 0.75, p = 0.03). The coefficient of the exponential argument ($\alpha$) was significantly larger in the non-pregnant uterine samples compared to the pregnant uterine samples (1.237 ± 1.226 vs. 0.440 ± 0.565, p = 0.04). No significant differences were found between the non-pregnant and pregnant values for the uterine Poisson's ratio ($\nu$) and the uterine fiber stiffness modulus ($\xi$).

##### 2.2 Cervix

The best-fit combination of parameters for all cervical samples, with the average values and standard deviations in the non-pregnant and pregnant state, is given in table 2. Boxplots of best-fit cervical parameters in the non-pregnant and pregnant states are shown in fig. 4, along with *P*-values from Welch's t-tests. The cervical fiber stiffness modulus ($\xi$) was significantly larger in the non-pregnant samples compared to the pregnant samples (200.22 $\pm$ 199.00 kPa vs. 9.22 $\pm$ 9.34, p = 0.03). The coefficient of the exponential argument ($\alpha$) was significantly larger in the non-pregnant cervical samples compared to the pregnant cervical samples (5.765 $\pm$ 1.938 vs. 0.083 $\pm$ 0.078, p = 7e-5). No significant differences were found between the non-pregnant and pregnant values for the cervical Young's modulus ($E$) and cervical Poisson's ratio ($\nu$).

#### 3 DISCUSSION

##### 3.1 Uterus

The inverse finite element analysis results from existing mechanical data on non-pregnant and pregnant human uterine tissue align with the existing literature, with modest changes in uterine material properties observed between the pregnant and non-pregnant states. Fang et al. investigated changes to uterine tissue material properties for one pregnant and one non-pregnant patient under spherical indentation, and did not find statistically different changes in Young's modulus ($E$), Poisson's ratio ($\nu$), or fiber stiffness modulus ($\xi$) [1]. The average material parameter values for the non-pregnant and pregnant uterine tissue found in Fang et al. 2021 are similar to those found in this work, except for the fiber stiffness modulus, which in this work was found to be an order of magnitude larger than in Fang et al. 2021 [1]. This difference may be due to the lack of tensile data in Fang et al., where the tensile mechanical response is dominated by the fiber material properties [1]. Additionally, $\alpha$ was set to be 5.7 and not considered as a fitted parameter [1].

Fodera et al. performed microindentation tests on non-pregnant and pregnant uterine tissue, assessing mechanical differences in the endometrial/decidual, myometrial, and perimetrial uterine layers [8]. Significant differences in Young's modulus ($E$) were found between layers, with the endometrial/decidual layer being less stiff than the myometrial layer in the non-pregnant and pregnant state. For the myometrium alone, no significant differences were found in Young's modulus between the non-pregnant and pregnant samples [8]. Young's moduli ranged from about 0.3kPa to 2.5kPa, which is similar to the values reported in this work.

##### 3.2 Cervix

The inverse finite element analysis results, based on existing mechanical data from non-pregnant and pregnant human cervical tissue, align with the literature, showing large decreases in tissue stiffness in the pregnant state compared to the non-pregnant state. Myers et al. fit ellipsoidally distributed fibers to existing uni-axial tension and compression data of non-pregnant and pregnant cervices [9]. Similar values for Young's modulus ($E$), Poisson's ratio ($\nu$), and fiber stiffness modulus ($\xi$) were found in the non-pregnant and pregnant tissues as found here (tab. 4). In addition to including fiber directionality, Myers et al. differed from the approach presented in the material model by setting $\alpha$ to 0, thereby using a fiber-power law and fitting $\beta$. The largest difference observed between the results in Myers et al. and this work is in the fiber stiffness modulus for pregnant cervices, which was found to be an order of magnitude larger in this work (0.7 kPa in Myers et al., 9.22 kPa in this work).

Shi et al. investigated cervical tissue material properties in seven non-pregnant cervices [7]. Cervical tissue samples were subjected to indentation tests at the internal and external os at locations in the anterior, left, posterior, and right stroma [7]. Material properties were found using inverse finite element analysis and a similar strain-energy density function was assumed [7]. The only difference between the model used by Shi et al. and the model used in this work is that Shi et al. used a von-Mises fiber distribution and the concentration parameter $b$ was left as a fitting parameter [7]. The best-fit parameters for the non-pregnant cervices found by Shi et al. are given in tab. 5. Similar values were found for Young's modulus ($E$), Poisson's ratio ($\nu$), and the exponential coefficient ($\alpha$). However, the fiber stiffness modulus ($\xi$), had a difference of two orders of magnitude. This difference may be due to the lack of tensile data in Shi et al., which is dominated by the fiber material properties [7].

#### 4 CONCLUSION ON MATERIAL PROPERTIES

The overall goal of this work is to model pregnant human reproductive tissues in the second trimester. Therefore, the values obtained directly from this inverse finite element analysis study are not appropriate for the intended use. To model pregnant human reproductive tissues in the second trimester, interpolation is required to obtain acceptable material values. Longitudinal studies on non-human primate cervical tissues have been published, which found that most of the change in material parameter value occurs between the non-pregnant and early third trimester [10]. Thus, values approximately halfway between those found for the non-pregnant and pregnant uterus will be assumed for uteri (tab. 6) and cervices (tab. 7) in the second trimester.

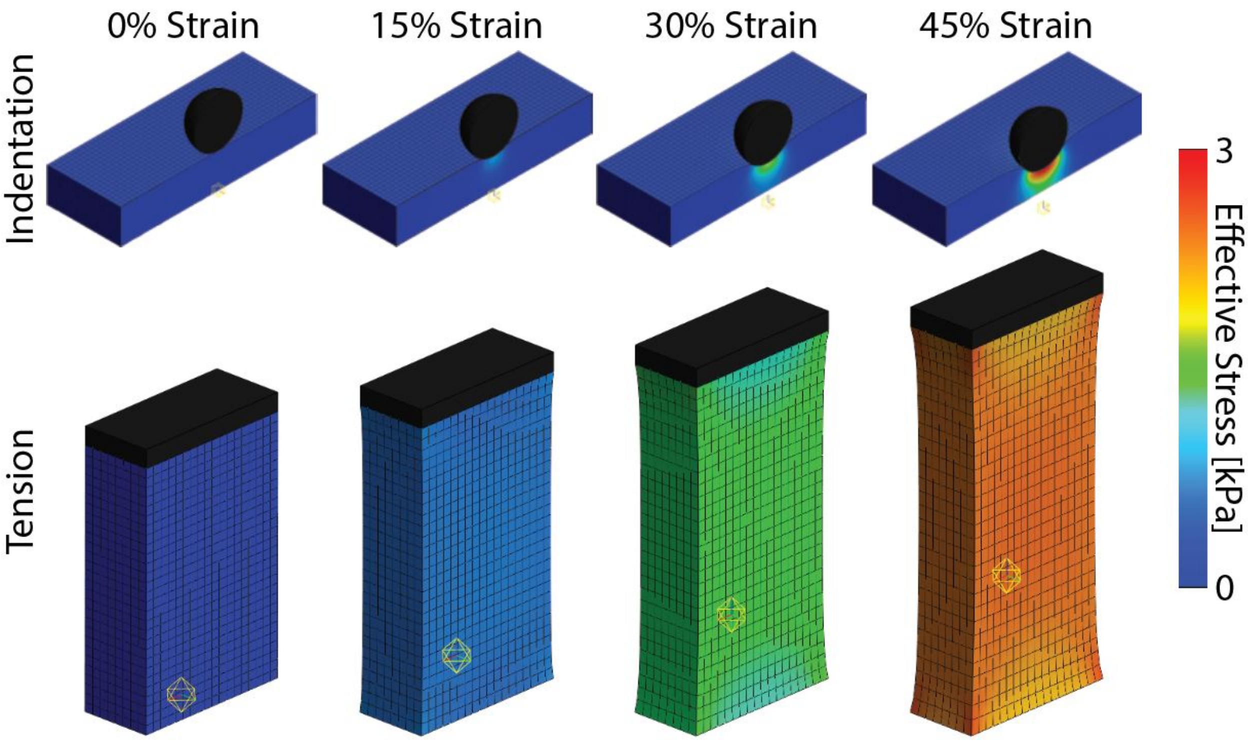

Fig. 1. Representative inverse finite element analysis model of indentation and tension tests on uterine tissue. Geometry in indentation and tension tests are not to scale. For scale, the indentor in the indentation tests had a diameter of 6mm, and the sample in the tension tests shown had an undeformed length of 14mm.

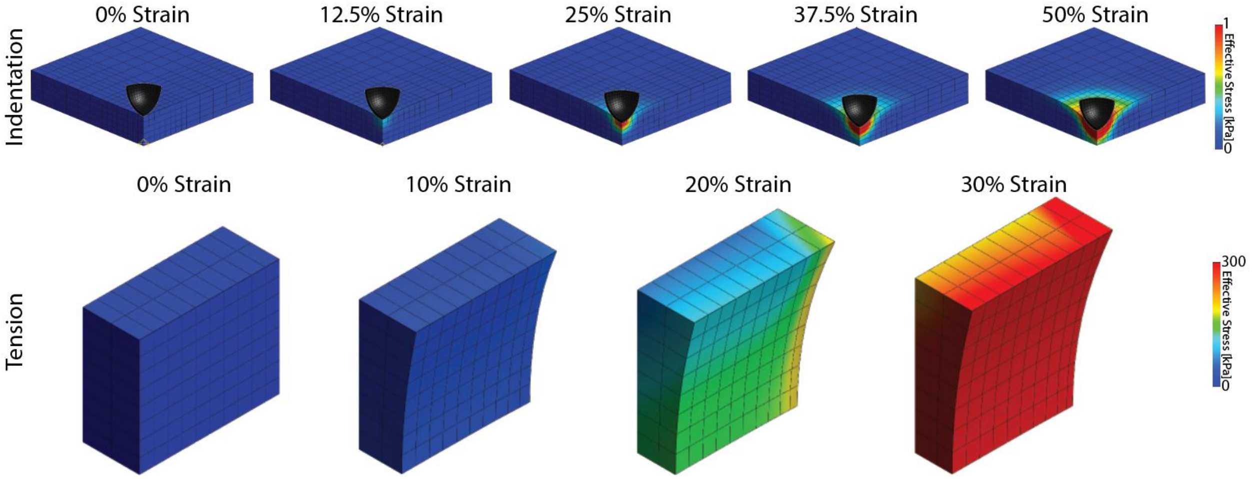

Fig. 2. Representative inverse finite element analysis model of indentation and tension tests on cervical tissue. Geometry in indentation and tension tests are not to scale. The number and level of tensile ramp-holds were not the same across cervical samples.

*FIGURES*

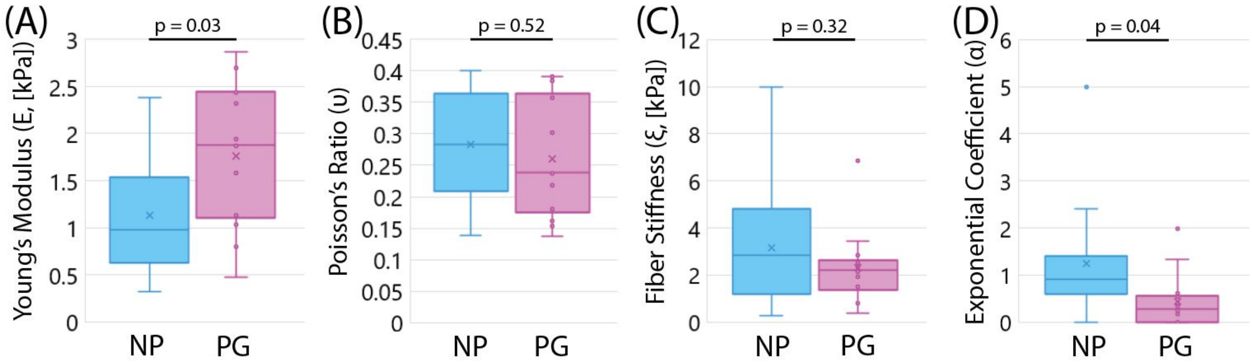

Fig. 3. Material properties of the human uterus (A) Young's modulus, (B) Poisson's ratio, (C) fiber stiffness, and (D) exponential coefficient for non-pregnant (NP) and pregnant (PG) uterine tissues from inverse finite element analysis, with p-values from Welch's t-tests displayed.

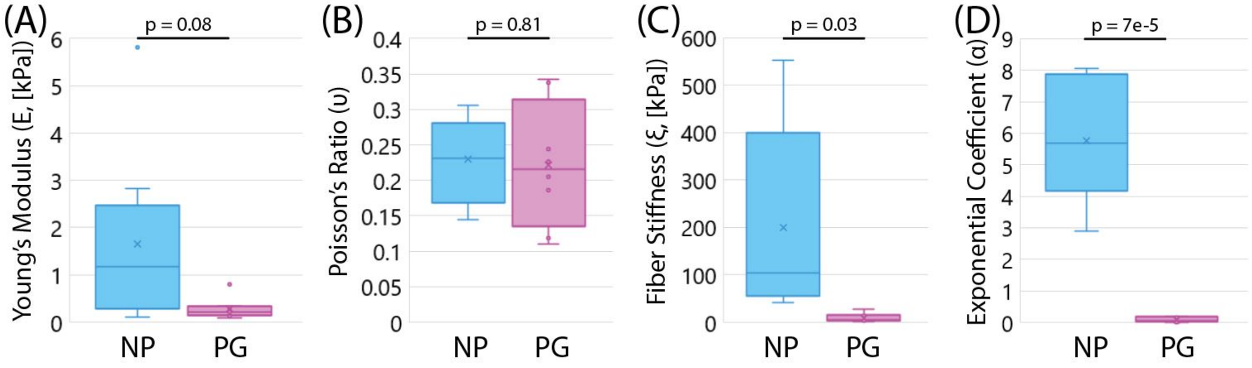

Fig. 4. Boxplots of (A) Young's modulus, (B) Poisson's ratio, (C) fiber stiffness, and (D) exponential coefficient for non-pregnant (NP) and pregnant (PG) cervical tissues from inverse finite element analysis, with p-values from Welch's t-tests displayed.

| Patient | Sample | NP/PG | $E$ [kPa] | $\nu$ | $\xi$ [kPa] | $\alpha$ | Error |
|---|---|---|---|---|---|---|---|
| 1 | 1 | NP | 0.68 | 0.263 | 10.00 | 0.827 | 0.300 |
| 1 | 2 | NP | 0.73 | 0.400 | 3.38 | 0.645 | 0.1851 |
| 2 | 1 | NP | 0.47 | 0.179 | 2.30 | 1.066 | 0.170 |
| 2 | 2 | NP | 2.26 | 0.263 | 0.51 | 1.590 | 0.194 |
| 3 | 1 | NP | 1.38 | 0.348 | 4.57 | 0.392 | 0.168 |
| 3 | 2 | NP | 2.38 | 0.372 | 3.69 | 0.972 | 0.167 |
| 4 | 1 | NP | 0.40 | 0.302 | 1.33 | 2.407 | 0.241 |
| 4 | 2 | NP | 0.32 | 0.361 | 5.53 | 1.335 | 0.261 |
| 5 | 1 | NP | 1.05 | 0.332 | 0.80 | 0.000 | 0.124 |
| 5 | 2 | NP | 1.20 | 0.218 | 0.28 | 0.880 | 0.161 |
| 6 | 1 | NP | 0.90 | 0.128 | 5.74 | 0.814 | 0.235 |
| 6 | 2 | NP | 0.70 | 0.250 | 3.37 | 5.000 | 0.235 |
| 7 | 1 | NP | 1.16 | 0.123 | 1.39 | 0.461 | 0.228 |
| 7 | 2 | NP | 1.38 | 0.392 | 1.32 | 0.930 | 0.249 |
| NP Average ± STDEV | | | 1.13 ± 0.68 | 0.283 ± 0.090 | 3.16 ± 2.67 | 1.237 ± 1.226 | - |
| 8 | 1 | PG | 2.32 | 0.182 | 2.14 | 0.304 | 0.199 |
| 8 | 2 | PG | 2.44 | 0.218 | 2.33 | 0.166 | 0.172 |
| 9 | 1 | PG | 0.80 | 0.240 | 2.52 | 0.396 | 0.177 |
| 9 | 2 | PG | 1.13 | 0.358 | 6.86 | 0.000 | 0.199 |
| 10 | 1 | PG | 1.03 | 0.391 | 1.91 | 0.000 | 0.154 |
| 10 | 2 | PG | 1.94 | 0.302 | 1.51 | 0.541 | 0.154 |
| 11 | 1 | PG | 0.47 | 0.391 | 3.44 | 0.000 | 0.294 |
| 11 | 2 | PG | 1.15 | 0.161 | 2.17 | 0.324 | 0.294 |
| 12 | 1 | PG | 2.87 | 0.153 | 0.81 | 1.337 | 0.228 |
| 12 | 2 | PG | 1.58 | 0.385 | 2.22 | 0.243 | 0.139 |
| 13 | 1 | PG | 1.89 | 0.305 | 2.85 | 0.000 | 0.192 |
| 13 | 2 | PG | 2.45 | 0.237 | 0.87 | 0.254 | 0.158 |
| 14 | 1 | PG | 2.70 | 0.137 | 0.38 | 1.974 | 0.216 |
| 14 | 2 | PG | 1.87 | 0.180 | 2.37 | 0.617 | 0.245 |
| PG Average ± STDEV | | | 1.76 ± 0.75 | 0.260 ± 0.094 | 2.31 ± 1.55 | 0.440 ± 0.565 | - |

Table 1. Inverse finite element analysis results of tension/indentation data from inverse finite element analysis on non-pregnant (NP) and pregnant (PG) uterine samples.

| Patient | Location | NP/PG | $E$ [kPa] | $\nu$ | $\xi$ [kPa] | $\alpha$ | Error |
|---|---|---|---|---|---|---|---|
| 1 | EO | NP | 1.38 | 0.232 | 205.16 | 4.135 | 0.182 |
| 1 | IO | NP | 5.81 | 0.306 | 47.71 | 8.055 | 0.143 |
| 2 | EO | NP | 0.17 | 0.282 | 122.40 | 4.258 | 0.428 |
| 2 | IO | NP | 1.18 | 0.221 | 465.74 | 2.882 | 0.307 |
| 3 | EO | NP | 0.65 | 0.151 | 87.14 | 8.054 | 0.467 |
| 3 | IO | NP | 2.83 | 0.230 | 553.29 | 5.582 | 0.264 |
| 4 | EO | NP | 0.10 | 0.276 | 41.48 | 5.804 | 0.394 |
| 4 | IO | NP | 1.16 | 0.144 | 78.85 | 7.352 | 0.218 |
| NP Average ± STDEV | | | 1.66 ± 1.88 | 0.230 ± 0.059 | 200.22 ± 199.00 | 5.765 ± 1.938 | - |
| 5 | EO | PG | 0.21 | 0.342 | 4.39 | 0.045 | 0.228 |
| 5 | IO | PG | 0.79 | 0.244 | 3.16 | 0.184 | 0.489 |
| 6 | EO | PG | 0.15 | 0.226 | 1.52 | 0.046 | 1.492 |
| 6 | IO | PG | 0.23 | 0.110 | 28.44 | 0.057 | 0.630 |
| 7 | EO | PG | 0.09 | 0.338 | 15.83 | 0.000 | 0.571 |
| 7 | IO | PG | 0.34 | 0.186 | 4.58 | 0.000 | 0.531 |
| 8 | EO | PG | 0.13 | 0.118 | 2.79 | 0.148 | 0.217 |
| 8 | IO | PG | 0.30 | 0.205 | 13.06 | 0.185 | 0.282 |
| PG Average ± STDEV | | | 0.28 ± 0.22 | 0.221 ± 0.087 | 9.22 ± 9.34 | 0.083 ± 0.078 | - |

Table 2. Inverse finite element analysis results of tension/indentation data from inverse finite element analysis on non-pregnant (NP) and pregnant (PG) cervical samples.

| Material parameter | Non-pregnant | Pregnant |
|---|---|---|
| Young's Modulus ($E$, [kPa]) | 1.9 | 1.7 |
| Poisson's Ratio ($\nu$) | 0.40 | 0.33 |
| Fiber stiffness modulus ($\xi$, [kPa]) | 0.24 | 0.35 |

Table 3. Approximate average uterine material parameter values from Fang et al., found via inverse finite element analysis on spherical indentation data [1].

| Material parameter | Non-pregnant | Pregnant |
|---|---|---|
| Young's Modulus ($E$, [kPa]) | 2 | 0.65 |
| Poisson's Ratio ($\nu$) | 0.3 | 0.3 |
| Fiber stiffness modulus ($\xi$, [kPa]) | 190 | 0.7 |

Table 4. Cervix material parameter values from Myers et al., found via inverse finite element analysis on uni-axial tension and compression data [9].

| Material parameter | Mean | Standard Deviation |
|---|---|---|
| Young's Modulus ($E$, [kPa]) | 3.20 | 3.30 |
| Poisson's Ratio ($\nu$) | 0.22 | 0.13 |
| Fiber stiffness modulus ($\xi$, [kPa]) | 3.59 | 5.51 |
| Exponential coefficient ($\alpha$) | 5.7 | 2.6 |

Table 5. Mean and standard deviation of cervical material parameter values from Shi et al., found via inverse finite element analysis on spherical indentation data [7].

| Material parameter | Assumed second trimester value |
|---|---|
| Young's modulus ($E$, [kPa]) | 1.5 |
| Poisson's ratio ($\nu$) | 0.27 |
| Fiber stiffness modulus ($\xi$, [kPa]) | 2.7 |
| Exponential coefficient ($\alpha$) | 0.74 |

Table 6. Uterine material parameter values for use in pregnant human reproductive anatomy models in the second trimester.

| Material parameter | Assumed second trimester value |
|---|---|
| Young's modulus ($E$, [kPa]) | 1.0 |
| Poisson's ratio ($\nu$) | 0.225 |
| Fiber stiffness modulus ($\xi$, [kPa]) | 104.7 |
| Exponential coefficient ($\alpha$) | 3 |

Table 7. Cervical material parameter values for use in pregnant human reproductive anatomy models in the second trimester.
