## Appendix B for "A finite element model of pregnancy derived from maternal sonography: effect of uterine and cervical structural properties on cervical mechanical loading"

### Appendix B: Approach to calculate stretch in the proximal cervix and mesh size convergence study results

**CONTENTS**

**1 Calculating local cylindrical stretch in the proximal cervix** **2**

**2 Mesh size convergence study** **6**

2.1 Methods . . . . . . 6

2.2 Results . . . . . . 6

### 1 CALCULATING LOCAL CYLINDRICAL STRETCH IN THE PROXIMAL CERVIX

A custom MATLAB (Mathworks, Natick, MA) script was developed to calculate the local cylindrical stretch at each node on the proximal cervix. The script then calculated the differences in the radial, longitudinal, and circumferential stretch fields between models to understand how model inputs affect the cervical response to membrane pressurization. First, the initial position (i.e., reference configuration) of each node on the proximal cervix face was exported from FEBio studio, along with the right stretch tensor components in the FEBio coordinate system in the final timestep, with the right stretch tensor (**U**) calculated by $\mathbf{F} = \mathbf{R} \cdot \mathbf{U}$, where **F** is the deformation gradient and **R** is the rotation tensor (fig. 1). These data were imported into MATLAB.

Within MATLAB, a series of orthogonal basis transformations on the position vector and stretch tensor were performed for each proximal cervix node (fig. 2). First, the origin was translated from the middle of the uterine cavity to the position of the cervical internal os (IO) (eq. 1):

$$\mathbf{X} = \begin{bmatrix} \mathrm{X_{FEBio}} \\ \mathrm{Y_{FEBio}} \\ \mathrm{Z_{FEBio}} \end{bmatrix} - \begin{bmatrix} \mathrm{IO_X} \\ \mathrm{IO_Y} \\ \mathrm{IO_Z} \end{bmatrix} \tag{1}$$

where **X** is the nodal position with the coordinate system origin placed at the cervical IO, $(\mathrm{X_{FEBio}}, \mathrm{Y_{FEBio}}, \mathrm{Z_{FEBio}})$ is the original nodal position with the coordinate system origin from FEBio, and $(\mathrm{IO_X}, \mathrm{IO_Y}, \mathrm{IO_Z})$ is the position of the cervical IO with the coordinate system from FEBio. Then, the coordinate system was rotated 90 degrees counterclockwise about the X-axis with orthogonal tensor $\mathbf{Q}_{\mathrm{Intermediate}}$, so the rotated Y-axis would be towards the left side of the anatomy.

$$\mathbf{Q}_{\mathrm{Intermediate}} = \begin{bmatrix} 1 & 0 & 0 \\ 0 & 0 & -1 \\ 0 & 1 & 0 \end{bmatrix} \tag{2}$$

Finally, the basis was rotated around the new Y-axis so that the Z-axis aligned with the cervical canal and the X-axis pointed to the anterior cervix. To do this, the vector pointing from the cervical external os (EO) to cervical IO ($\mathbf{c}_{\mathrm{FEBio}}$) was defined in the FEBio coordinate system (eq. 3):

$$\mathbf{c}_{\mathrm{FEBio}} = \begin{bmatrix} \mathrm{IO_X} \\ \mathrm{IO_Y} \\ \mathrm{IO_Z} \end{bmatrix} - \begin{bmatrix} \mathrm{EO_X} \\ \mathrm{EO_Y} \\ \mathrm{EO_Z} \end{bmatrix} \tag{3}$$

where $(\mathrm{EO_X}, \mathrm{EO_Y}, \mathrm{EO_Z})$ is the position of the cervical EO with the coordinate system from FEBio. The unit vector in the direction from the cervical EO to cervical IO ($\mathbf{n}_{\mathrm{c,FEBio}}$) was then found with equation 4:

$$\mathbf{n}_{\text{c,FEBio}} = \frac{\mathbf{c}_{\text{FEBio}}}{\sqrt{\mathbf{c}_{\text{FEBio}} \cdot \mathbf{c}_{\text{FEBio}}}} \tag{4}$$

An orthogonal basis transformation was then applied to $\mathbf{n}_{\text{c,FEBio}}$ to be in the coordinate system with the intermediate coordinate system to get $\mathbf{n}_{\text{c,Intermediate}}$ via equation 5:

$$\mathbf{n}_{\text{c,Intermediate}} = \mathbf{Q}_{\text{Intermediate}} \cdot \mathbf{n}_{\text{c,FEBio}} \tag{5}$$

Then, the angle between the Z-axis in the intermediate coordinate system and cervical canal direction ($\alpha_{\text{c}}$) was found using equation 6:

$$\alpha_{\text{c}} = \cos^{-1}(\mathbf{n}_{\text{c,Intermediate}} \cdot \mathbf{n}_{\text{Z,Intermediate}}) \tag{6}$$

where $\mathbf{n}_{\text{Z,Intermediate}}$ is the unit vector in the Z direction in the intermediate coordinate system. With $\alpha_{\text{c}}$ defined, the orthogonal tensor $\mathbf{Q}_{\text{Cartesian}}$ was defined by equation 7:

$$\mathbf{Q}_{\text{Cartesian}} = \begin{bmatrix} \cos\alpha_{\text{c}} & 0 & \sin\alpha_{\text{c}} \\ 0 & 1 & 0 \\ -\sin\alpha_{\text{c}} & 0 & \cos\alpha_{\text{c}} \end{bmatrix} \tag{7}$$

Therefore, the final nodal position ($\mathbf{X}_{\text{Cartesian}}$) basis transformation is given by equation 8

$$\mathbf{X}_{\text{Cartesian}} = \mathbf{Q}_{\text{Cartesian}} \cdot \mathbf{Q}_{\text{Intermediate}} \cdot \mathbf{X} \tag{8}$$

With the Cartesian coordinate system transformed to align with the cervical geometry, the Cartesian coordinates were then converted to cylindrical coordinates through equation 9:

$$\mathbf{X}_{\text{Cylindrical}} = \begin{bmatrix} \text{r} \\ \theta \\ \text{z} \end{bmatrix} = \begin{bmatrix} \sqrt{\text{X}^2_{\text{Cartesian}} + \text{Y}^2_{\text{Cartesian}}} \\ \tan^{-1} \frac{\text{Y}_{\text{Cartesian}}}{\text{X}_{\text{Cartesian}}} \\ \text{Z} \end{bmatrix} \tag{9}$$

With the cylindrical position coordinates established, the cylindrical stretch tensor ($\mathbf{U}_{\text{Cylindrical}}$) was found using equation 10:

$$\mathbf{U}_{\text{Cylindrical}} = \mathbf{Q}_{\text{Cylindrical}} \cdot \mathbf{Q}_{\text{Cartesian}} \cdot \mathbf{Q}_{\text{Intermediate}} \cdot \mathbf{U}_{\text{FEBio}} \cdot \mathbf{Q}^{\text{T}}_{\text{Intermediate}} \cdot \mathbf{Q}^{\text{T}}_{\text{Cartesian}} \cdot \mathbf{Q}^{\text{T}}_{\text{Cylindrical}} \tag{10}$$

where $\mathbf{U}_{\text{FEBio}}$ was the right stretch in the FEBio coordinate system and $\mathbf{Q}_{\text{Cylindrical}}$ was given by equation 11:

$$\mathbf{Q}_{\text{Cylindrical}} = \begin{bmatrix} \cos\theta & \sin\theta & 0 \\ -\sin\theta & \cos\theta & 0 \\ 0 & 0 & 1 \end{bmatrix} \tag{11}$$

With the cylindrical coordinate system established, the curvature of the proximal cervix face was examined, and it was found that a loacl definition of the radial, circumferential, and normal directions were needed to define the proximal cervix face (fig. 3). The proximal cervix face had to be divided into radial segments to find the local surface radial, circumferential, and normal directions (fig. 3). The number of radial segments was set to 50 after a sensitivity study showed this number of segments to yield the most consistent results. In each segment, a bivariate second-degree polynomial was fit to the $(\text{r}, \theta, \text{z})$ nodal positions, fitting z as a function of r and $\theta$, yielding equation 12 for each radial segment.

$$\text{z}(\text{r}, \theta) = a_N \text{r}^2 \cos^2\theta + b_N \text{r}^2 \sin^2\theta + c_N \text{r}^2 \cos\theta \sin\theta + d_N \text{r} \cos\theta + e_N \text{r} \sin\theta + f_N \tag{12}$$

where $a_N$, $b_N$, $c_N$, $d_N$, $e_N$, and $f_N$ are the best-fit coeffients for second-degree polynomial in radial slice $N$.

With eq. 12, the surface radial and circumferential directions could be calculated using the local slope ($m_{\text{r}}$ and $m_\theta$, respectively) given the nodal position $(\text{r}, \theta, \text{z})$ using equations 13 and 14:

$$m_{\text{r}}(\text{r}, \theta) = \frac{\partial \text{z}(\text{r}, \theta)}{\partial \text{r}} = 2a_N \text{r} \cos^2\theta + \sin\theta(2b_N \text{r} \sin\theta + e_N) + \cos\theta(2c_N \text{r} \sin\theta + d_N) \tag{13}$$

$$m_\theta(\text{r}, \theta) = \frac{\partial \text{z}(\text{r}, \theta)}{\partial \theta} = \text{r} \cos\theta(2\text{r}(b_N - a_N) \sin\theta + e_N) - \text{r} \sin\theta(c_N \text{r} \sin\theta + d_N) + c_N \text{r}^2 \cos^2\theta \tag{14}$$

The local slope could then be used to define the local surface tangent vectors $\mathbf{v}_{\text{local radial}}(\text{r}, \theta) = [1, 0, m_{\text{r}}(\text{r}, \theta)]$ and $\mathbf{v}_{\text{local circumferential}}(\text{r}, \theta) = [0, 1, m_\theta(\text{r}, \theta)]$. Thus, the unit vector in the direction of the local surface radial tangent vector ($\mathbf{r}'$) and local surface circumferential tangent ($\boldsymbol{\theta}'$) can be calculated using equation 15 and 16:

$$\mathbf{r}'(\mathrm{r},\theta) = \frac{\mathbf{v}_{\text{local radial}}(\mathrm{r},\theta)}{\sqrt{\mathbf{v}_{\text{local radial}}(\mathrm{r},\theta) \cdot \mathbf{v}_{\text{local radial}}(\mathrm{r},\theta)}} \tag{15}$$

$$\boldsymbol{\theta}'(\mathrm{r},\theta) = \frac{\mathbf{v}_{\text{local circumferential}}(\mathrm{r},\theta)}{\sqrt{\mathbf{v}_{\text{local circumferential}}(\mathrm{r},\theta) \cdot \mathbf{v}_{\text{local circumferential}}(\mathrm{r},\theta)}} \tag{16}$$

The local surface normal vector ($\mathbf{z}'$) was found by taking the cross product between $\mathbf{r}'$ and $\boldsymbol{\theta}'$ ($\mathbf{z}'(\mathrm{r},\theta) = \mathbf{r}'(\mathrm{r},\theta) \times \boldsymbol{\theta}'(\mathrm{r},\theta)$). With these unit vectors established, the local surface radial stretch ($\lambda_{\mathrm{r}'}$) was calculated using equation 17, the local surface circumferential stretch ($\lambda_{\theta}$) was calculated using equation 18, and the local surface normal stretch was calculated using equation 19. Spatial heat maps were then generated for stretch in the local surface radial, circumferential, and normal directions using the $(\mathrm{r},\theta)$ position data.

$$\lambda_{\mathrm{r}'} = \mathbf{r}' \cdot \mathbf{U}_{\text{Cylindrical}} \cdot \mathbf{r}' \tag{17}$$

$$\lambda_{\theta} = \boldsymbol{\theta} \cdot \mathbf{U}_{\text{Cylindrical}} \cdot \boldsymbol{\theta} \tag{18}$$

$$\lambda_{\mathrm{z}'} = \mathbf{z}' \cdot \mathbf{U}_{\text{Cylindrical}} \cdot \mathbf{z}' \tag{19}$$

To compare proximal cervix stretch fields between models, stretch difference heat maps were generated. A target grid was created to generate the difference heat maps, with the inner boundary of the grid set by the model with the larger inner boundary and the outer boundary of the grid set by the model with the smaller outer boundary. The target grid was necessary to spatially match the nodes between models, as meshes varied between models. The target grid was divided into radial and circumferential sections, and the nodes from each model closest to the target grid points were selected for comparison. Stretch differences were found by subtracting the first model's stretch values from the second model's stretch values at the nodes closest to the target grid points. These stretch differences were then plotted with the target grid position to generate the spatial difference heat maps.

#### 2 MESH SIZE CONVERGENCE STUDY

##### 2.1 Methods

A meshing discretization scheme was determined for use in patient-specific second-trimester models of pregnant anatomy. A mesh size convergence study was undertaken to determine the optimal mesh size for accurate results. The study focused on the element size used in the cervix and fetal membrane, as these are most likely to have the largest effect on proximal cervix stretch results. Hypermesh 2021.1 (Altair, Troy, MI) was used to discretize the geometry, and the settings given in table 1 were used for mesh refinement. The resulting proximal cervix face meshes are shown in fig. 4, with all quadratic tetrahedral elements in the cervix. The desired convergence criterion was a 5% or less difference between models for the variables of interest. The variables of interest were Green-Lagrange strain in the anterior, left, and posterior proximal cervix region and the proximal cervix heat maps. The percent difference was calculated using equation 20:

$$\text{error} = \frac{E_{coarser} - E_{finer}}{E_{coarser}} \tag{20}$$

where $E_{coarser}$ is the Green-Lagrange strain in the model with the coarser mesh, and $E_{finer}$ is the Green-Lagrange strain in the model with the finer mesh. The right stretch tensor (**U**) is related to the Green-Lagrange strain tensor (**E**) by equation 21 [1]:

$$\mathbf{E} = \frac{1}{2}(\mathbf{U}^2 - \mathbf{I}) \tag{21}$$

where **I** is the identity tensor. The Green-Lagrange strain was used as the variable for convergence rather than stretch values, which are naturally larger and would produce artificially small errors. For example, if $\lambda_{\mathrm{r}',coarser} = 1.1$ and $\lambda_{\mathrm{r}',finer} = 1.2$, the error given by equation 20 would be -0.09, or a 9% error. For those values of stretch, the corresponding Green-Lagrange strain would be $E_{\mathrm{r}',coarser} = 0.11$ and $E_{\mathrm{r}',finer} = 0.22$, producing an error of -1.10, or 110%. Therefore, given the form of equation 20, Green-Lagrange strain is a more appropriate variable of interest for the mesh size convergence study.

After good convergence was found for the cervix mesh, a mesh size convergence study was performed for the fetal membrane. The fetal membrane was meshed with a single-layer quadratic hexahedral scheme following the settings in table 2. The resulting fetal membrane meshes are in fig. 5. Several finer meshes were attempted in the fetal membrane but did not reach full convergence and are not included.

##### 2.2 Results

Time to full convergence for each mesh size convergence study model with varied meshing scheme in the cervix is given in tab. 3. Computation time generally increased with increasing mesh size, with the largest changes occurring after the model with 108017 cervical elements.

The surface radial, circumferential, and normal strains in the anterior, left, and posterior portions changed slightly with the different mesh sizes (fig. 6). In the local radial direction, strain increased slightly in all analyzed portions with increasing number of cervical elements (<1% strain).

The local circumferential strain stayed rather constant in all analyzed portions with increasing number of cervical elements (¡0.5% strain). In the local normal direction, strain decreased in all analyzed portions with increasing number of cervical elements (¡1% strain). All strains in all analyzed portions stayed rather constant with models that had greater than or equal to 78506 elements,

In the local radial direction, strain did not reach less than 5% error across the entire cervical face for any of the mesh sizes attempted (fig. 7). However, the area that did not reach less than 5% error was the region with the smallest strains, that closest to the cervical IO ($<0.05$). The rest of the proximal cervical face reached less than 5% error in all but 4 points in the model with 108017 cervical elements (as seen compared to the model with 150643 cervical elements).

In the local circumferential direction, strain did not reach less than 5% error across the entire cervical face for any of the mesh sizes attempted (fig. 8). However, the area that did not reach less than 5% error was the region with the smallest strains, which was in the mid-stroma to uterocervical junction ($<0.03$). The cervical IO and anterior/posterior regions reached less than 5% error in the model with 108017 cervical elements (as seen compared to the model with 150643 cervical elements).

In the surface normal direction, strain did reach less than 5% error across the entire cervical face within the mesh sizes attempted (fig. 9). Convergence to less than 5% error in strain was reached by the model with 31403 cervical elements (as seen compared to the model with 41788 cervical elements).

Time to full convergence for each mesh size convergence study model with a varied meshing scheme in the fetal membrane is given in the tab. 4. Computation time generally increased with increasing mesh size.

For the local radial, circumferential, and normal strains in the anterior, left, and posterior portions of the proximal cervix, little change was seen with increasing number of elements in the fetal membrane (fig. 10). Within the tested range, strain was considered converged in all portions and directions for the tested mesh sizes.

In the local radial direction, less than 5% error in strain was not reached across the proximal cervix face for the fetal membrane mesh sizes tested (fig. 11). However, the only area that did not fully converge was that nearest the cervical IO, where the surface radial tangent strains are the smallest. For the rest of the proximal cervix face, surface radial tangent strain converged with less than 5% error in strain by the model with 1843 elements in the fetal membrane (as compared to the model with 2283 elements in the fetal membrane).

In the local circumferential direction, less than 5% error in strain was not reached across the proximal cervix face for the fetal membrane mesh sizes tested (fig. 12). However, the area that did not converge is that with the smallest strains (left and right mid-stroma to uterocervical junction). The area with the higher levels of circumferential strain (near the cervical IO and in the anterior/posterior proximal cervix) converged with less than 5% error in strain by the model with 2283 fetal membrane elements (as compared to the model with 3107 elements fetal membrane).

In the local normal direction, less than 5% error in strain was reached within the fetal membrane mesh sizes tested (fig. 13). Less than 5% error in surface normal strain was reached across the proximal cervix face by the model with 1843 elements in the fetal membrane (as compared to the model with 2283 elements in the fetal membrane).

#### LIST OF FIGURES

1 The initial position coordinates and final stretch tensor for all nodes on the proximal cervix surface (highlighted yellow) were exported from FEBio Studio and imported in MATLAB for cylindrical stretch analysis. . . . . . . . . . . . . . . . . . . . . . . . . 10
2 The cylindrical coordinate system was established through a series of coordinate transformations. First, the coordinate system was translated so that the origin aligned with the cervical IO coordinate. Second, the coordinate system was rotated 90 degrees counter-clockwise (CCW) about the X axis, such that the Y axis was oriented to the left. Third, the coordinate system was rotated about Y such that Z was aligned with the cervical canal, with X in the direction of the anterior cervix and Y in the direction of the left cervix. Finally, X and Y were converted to polar coordinates, r and $\theta$. . . . . . . . . . . . . . . . . . . . . . . . . . . . . . . . . 11
3 To create local cylindrical coordinates, the proximal cervix face (highlighted blue) was split up into many radial sections (purple). For each section, a second-degree bivariate polynomial was fit to the nodal position data (yellow). Using this polynomial, the loca surface radial (r'), circumferential ($\theta$'), and normal (z') directions were defined at each node on the proximal surface. . . . . . . . . . . . . . . . . . . . . . . . . 12
4 Resulting discretization in the proximal cervix for mesh settings given in tab. 1. . . . 13
5 Resulting discretization in the fetal membrane for mesh settings given in tab. 2. . . . 14
6 Mean and standard deviation of the surface (Radial) tangent, Circumferential, and surface normal (Longitudinal) stretch in the anterior, left, and posterior proximal cervix face with the number of elements in the cervix. . . . . . . . . . . . . . . . . . 15
7 Heat maps of surface radial tangent stretch with the increasing number of elements in the cervix. . . . . . . . . . . . . . . . . . . . . . . . . . . . . . . . . . . . . . 16
8 Heat maps of surface circumferential stretch with the increasing number of elements in the cervix. . . . . . . . . . . . . . . . . . . . . . . . . . . . . . . . . . . . . . 17
9 Heat maps of surface normal stretch with increasing the number of elements in the cervix. . . . . . . . . . . . . . . . . . . . . . . . . . . . . . . . . . . . . . . . . . 18
10 Mean and standard deviation of the surface (Radial) tangent, Circumferential, and surface normal (Longitudinal) stretch in the anterior, left, and posterior proximal cervix face with the number of elements in the fetal membrane. . . . . . . . . . . . . 19
11 Heat maps of surface radial tangent stretch with the increasing number of elements in the fetal membrane. . . . . . . . . . . . . . . . . . . . . . . . . . . . . . . . . . 20
12 Heat maps of surface circumferential stretch with the increasing number of elements in the fetal membrane. . . . . . . . . . . . . . . . . . . . . . . . . . . . . . . . . . 21
13 Heat maps of surface normal stretch with the increasing number of elements in the fetal membrane. . . . . . . . . . . . . . . . . . . . . . . . . . . . . . . . . . . . . 22

*FIGURES*

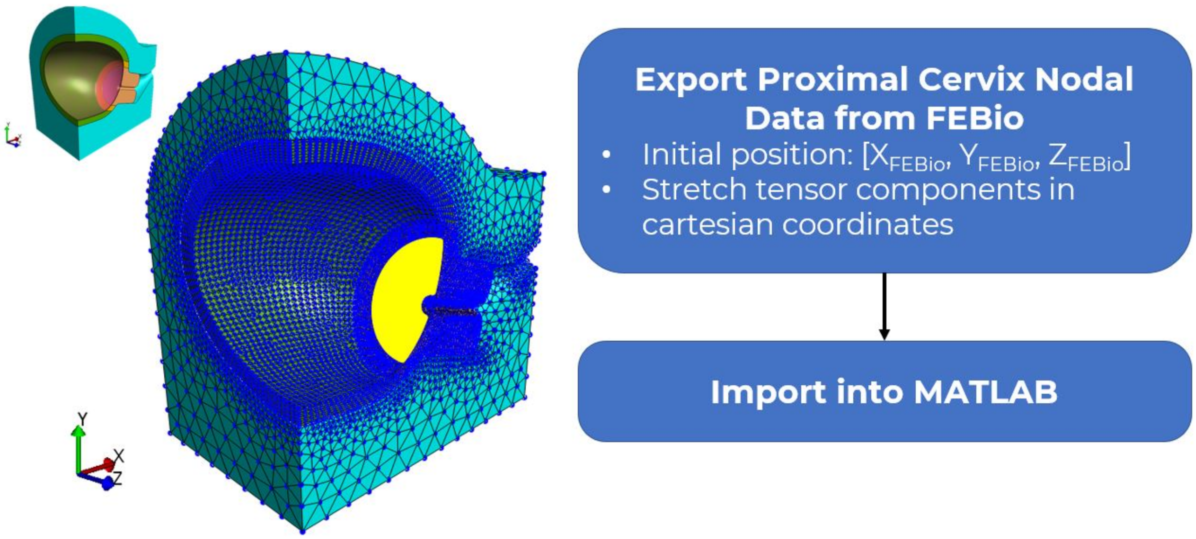

Fig. 1. The initial position coordinates and final stretch tensor for all nodes on the proximal cervix surface (highlighted yellow) were exported from FEBio Studio and imported in MATLAB for cylindrical stretch analysis.

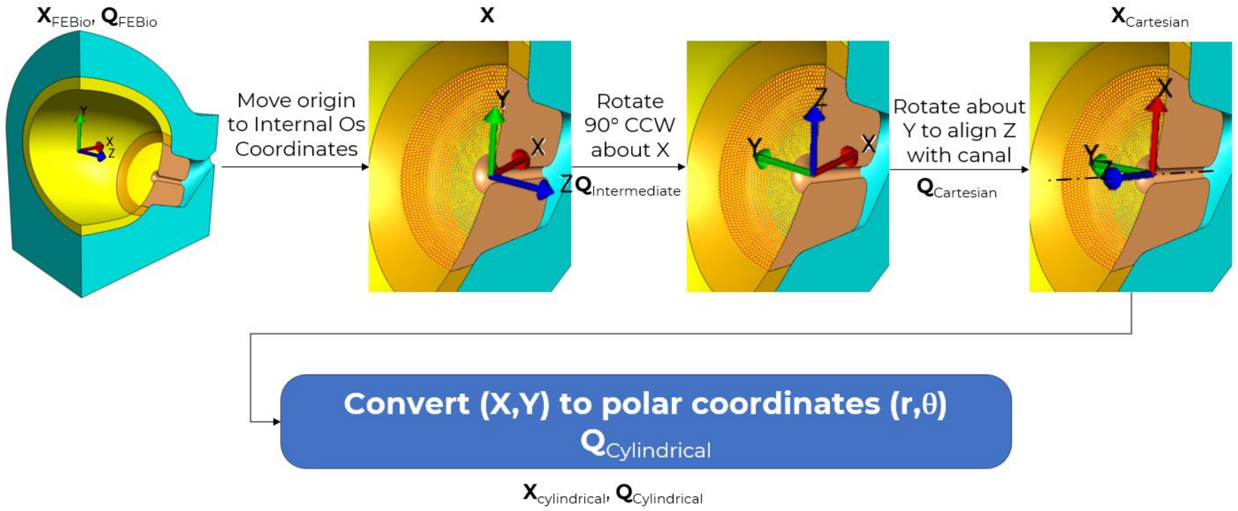

Fig. 2. The cylindrical coordinate system was established through a series of coordinate transformations. First, the coordinate system was translated so that the origin aligned with the cervical IO coordinate. Second, the coordinate system was rotated 90 degrees counter-clockwise (CCW) about the X axis, such that the Y axis was oriented to the left. Third, the coordinate system was rotated about Y such that Z was aligned with the cervical canal, with X in the direction of the anterior cervix and Y in the direction of the left cervix. Finally, X and Y were converted to polar coordinates, r and $\theta$.

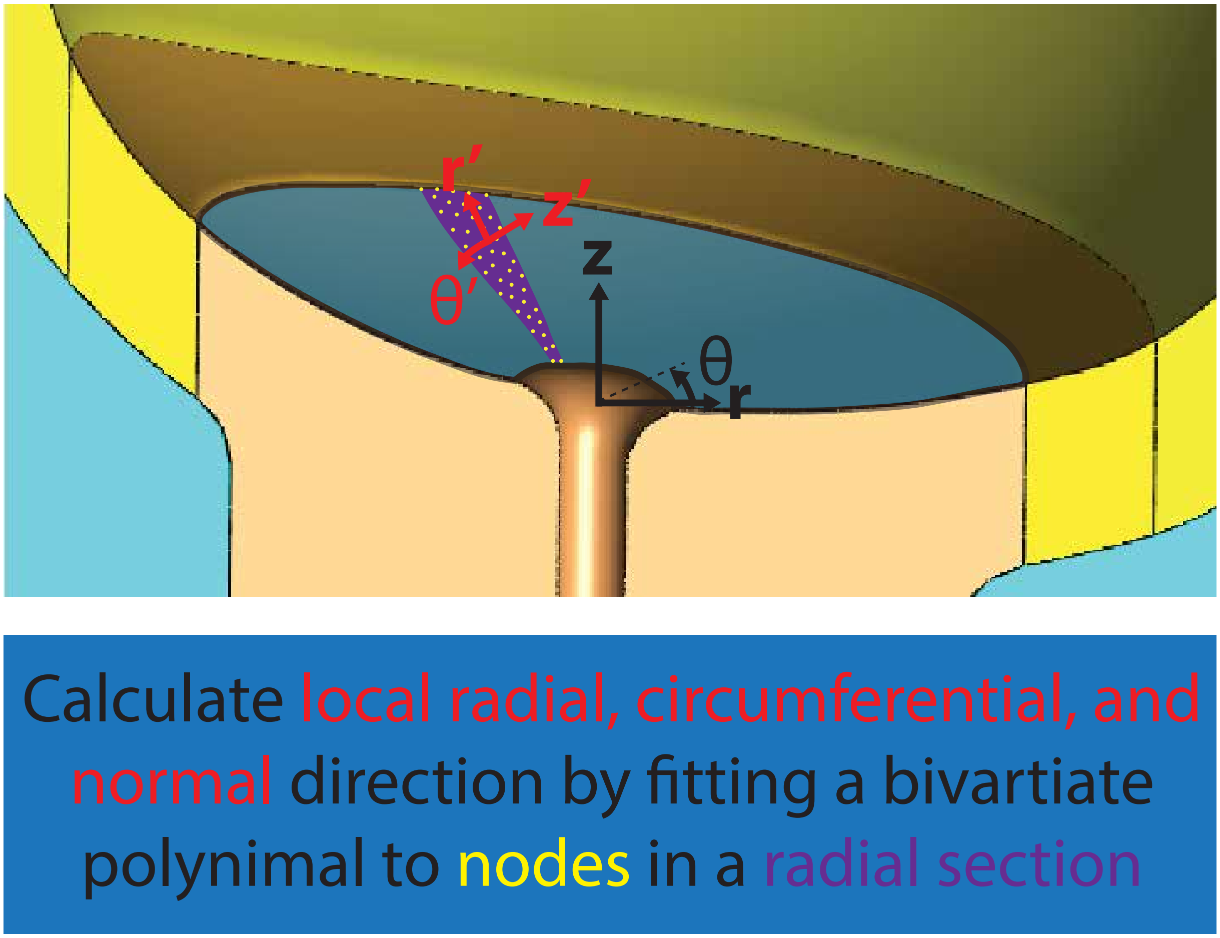

Fig. 3. To create local cylindrical coordinates, the proximal cervix face (highlighted blue) was split up into many radial sections (purple). For each section, a second-degree bivariate polynomial was fit to the nodal position data (yellow). Using this polynomial, the loca surface radial (r’), circumferential ($\theta$’), and normal (z’) directions were defined at each node on the proximal surface.

### FIGURES

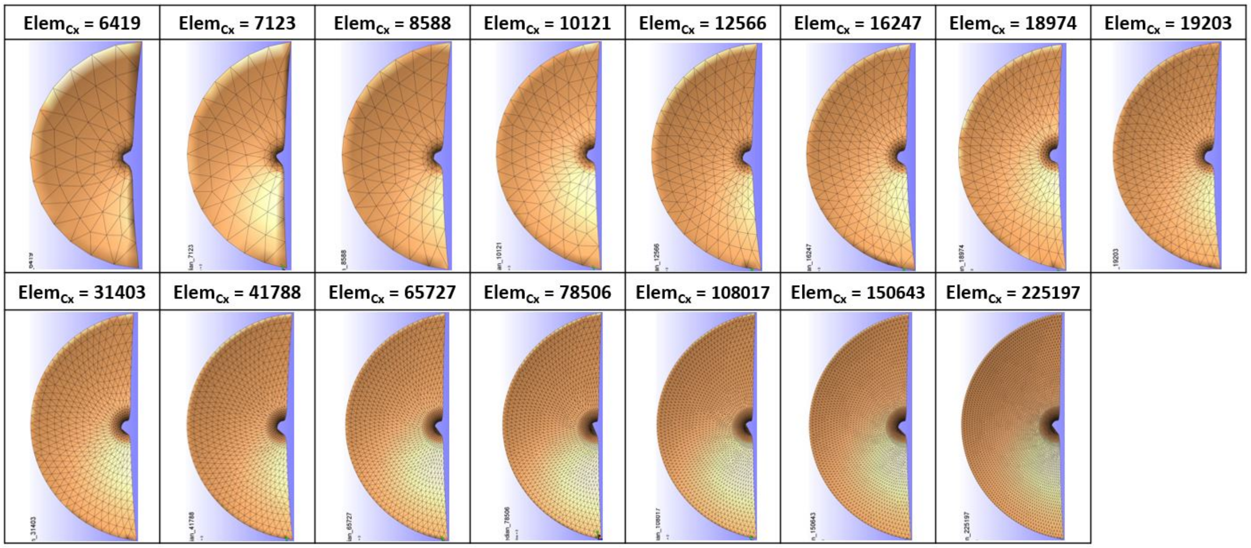

Fig. 4. Resulting discretization in the proximal cervix for mesh settings given in tab. 1.

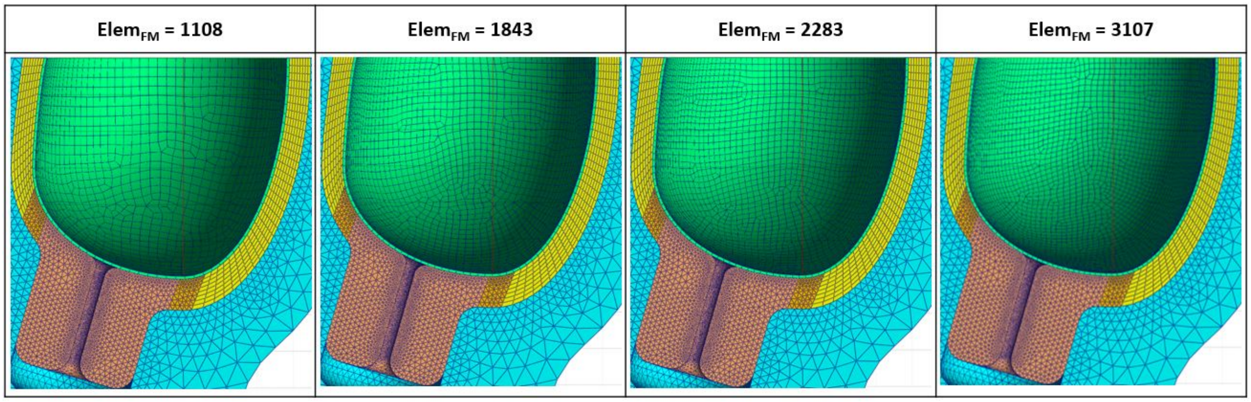

Fig. 5. Resulting discretization in the fetal membrane for mesh settings given in tab. 2.

*FIGURES*

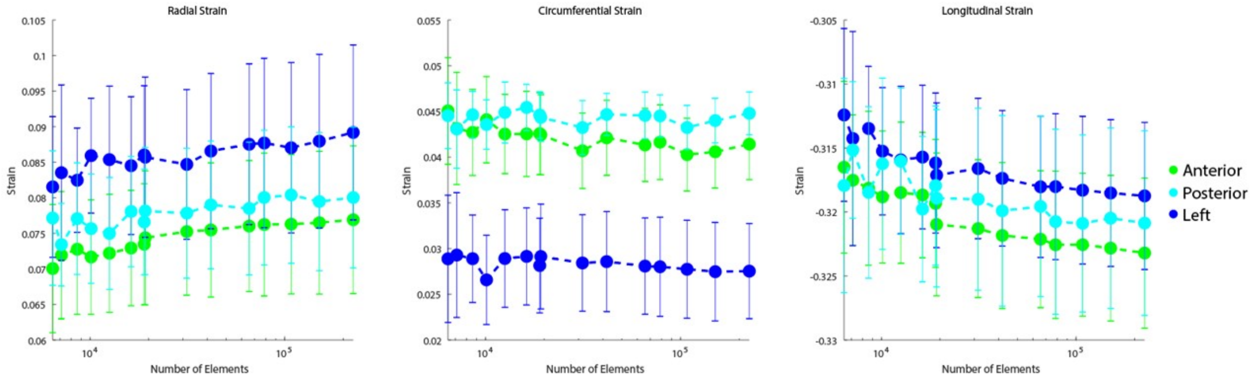

Fig. 6. Mean and standard deviation of the surface (Radial) tangent, Circumferential, and surface normal (Longitudinal) stretch in the anterior, left, and posterior proximal cervix face with the number of elements in the cervix.

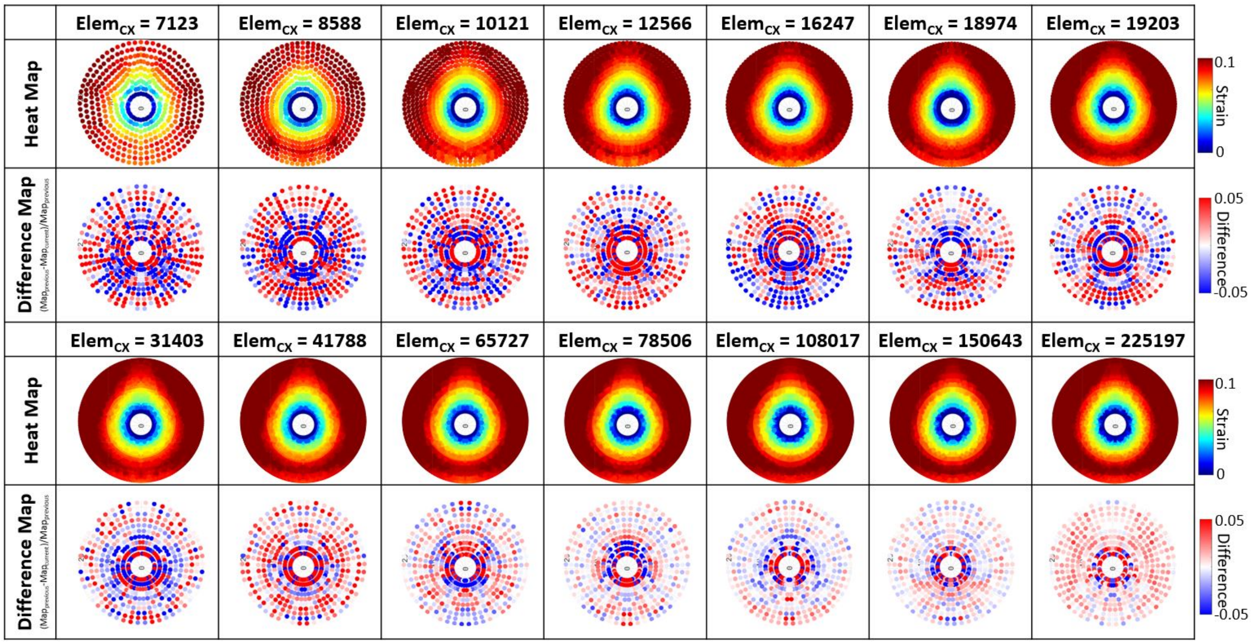

Fig. 7.  Heat maps of surface radial tangent stretch with the increasing number of elements in the cervix.

*FIGURES*

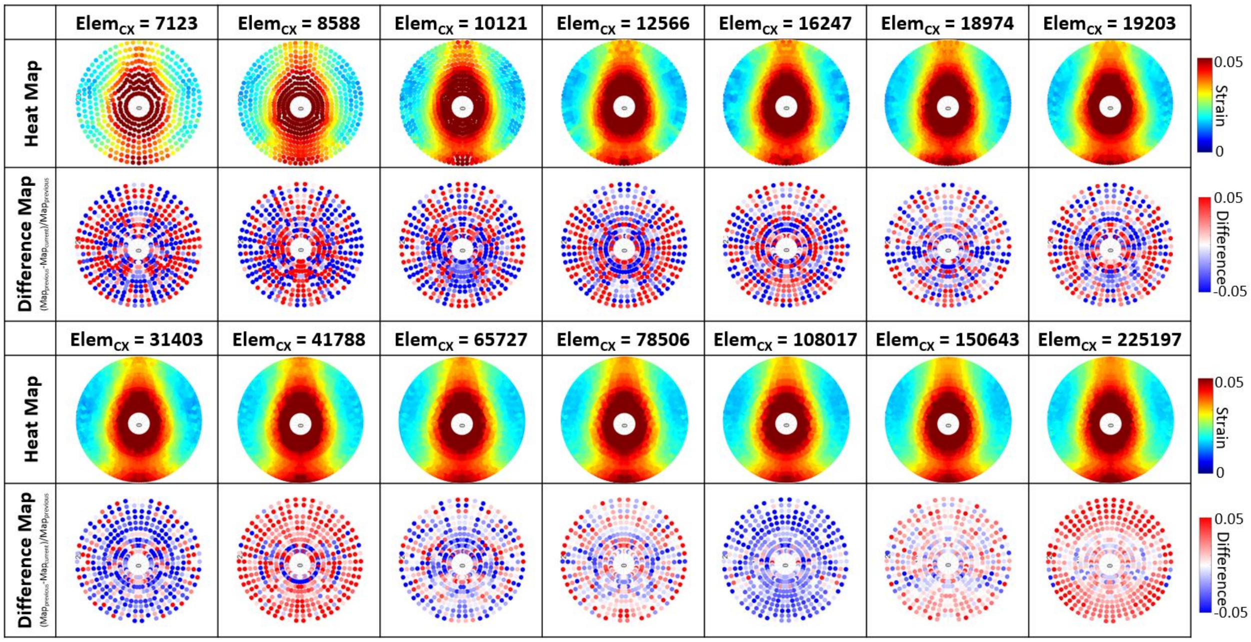

Fig. 8. Heat maps of surface circumferential stretch with the increasing number of elements in the cervix.

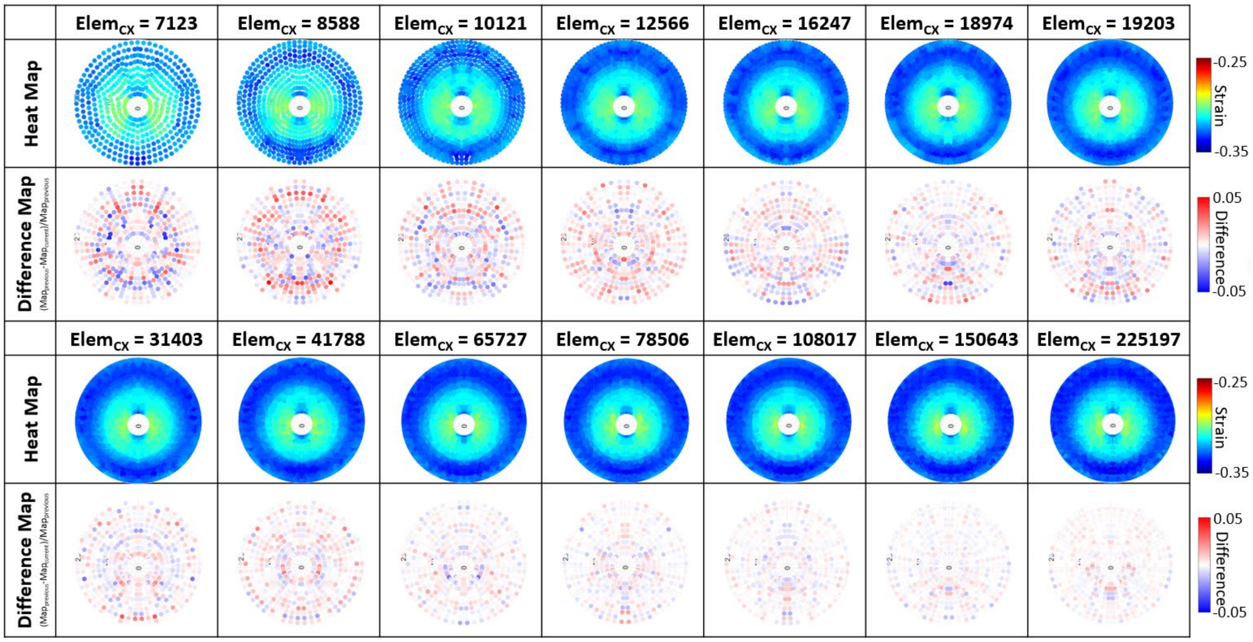

Fig. 9. Heat maps of surface normal stretch with increasing the number of elements in the cervix.

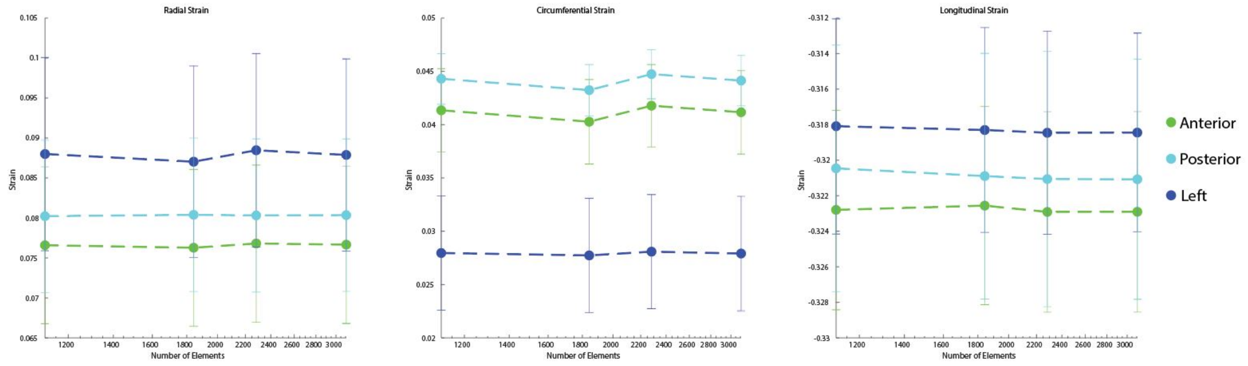

Fig. 10. Mean and standard deviation of the surface (Radial) tangent, Circumferential, and surface normal (Longitudinal) stretch in the anterior, left, and posterior proximal cervix face with the number of elements in the fetal membrane.

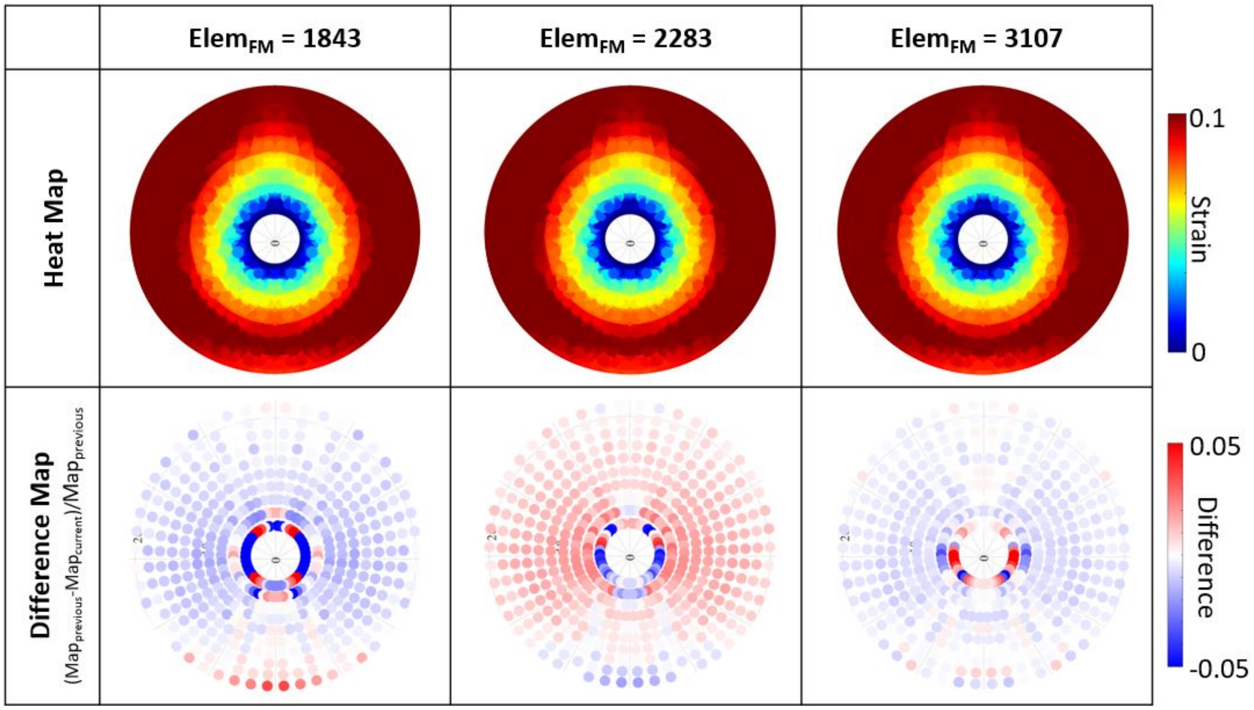

Fig. 11. Heat maps of surface radial tangent stretch with the increasing number of elements in the fetal membrane.

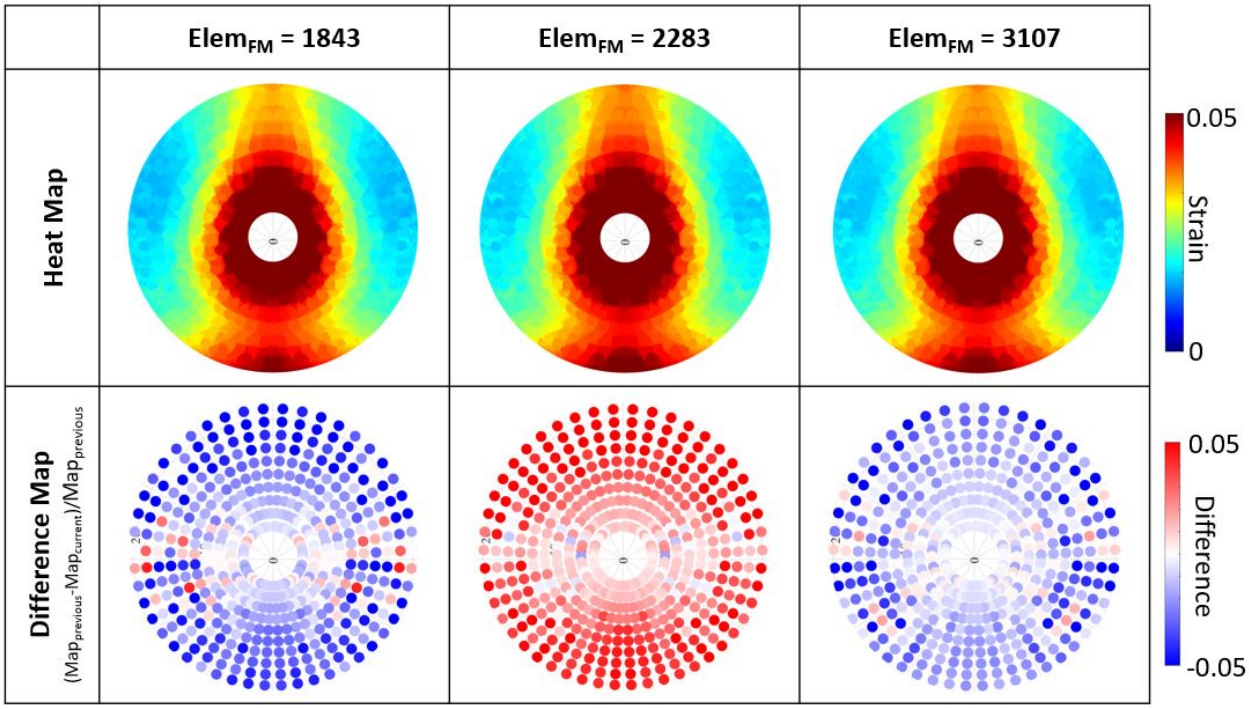

Fig. 12. Heat maps of surface circumferential stretch with the increasing number of elements in the fetal membrane.

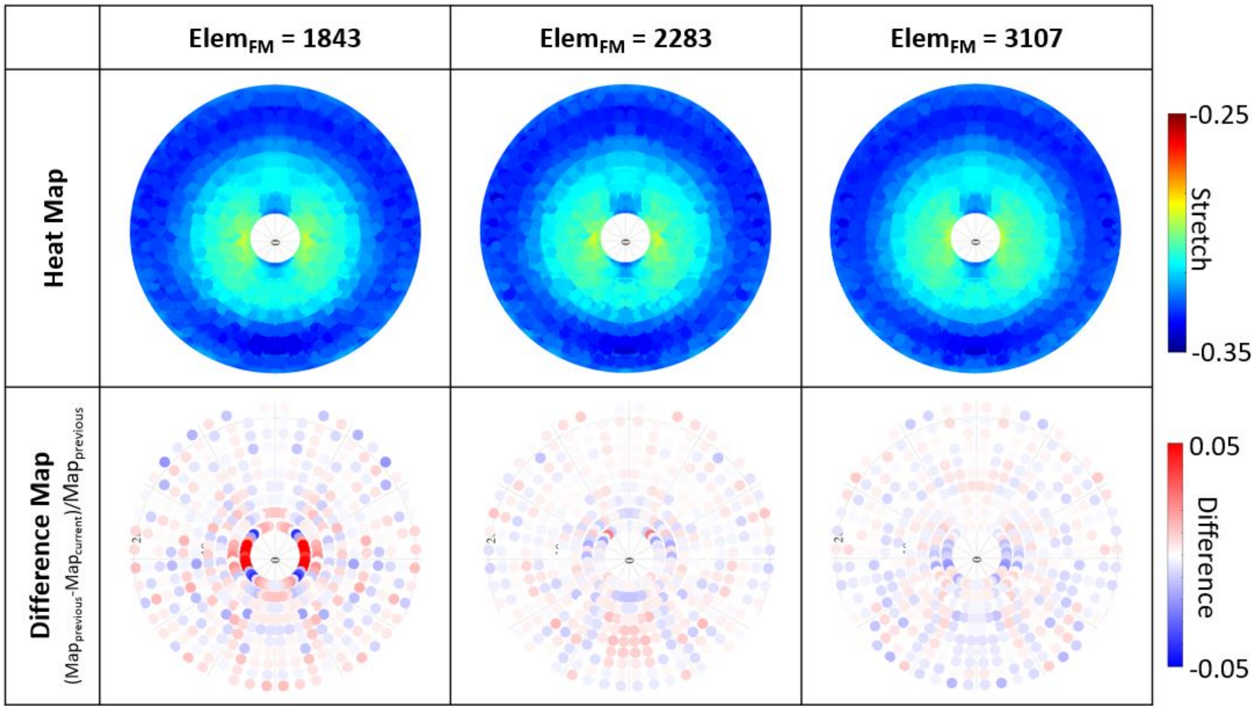

Fig. 13. Heat maps of surface normal stretch with the increasing number of elements in the fetal membrane.

| Canal size | Face size | Element size | Number of cervix elements |
|---|---|---|---|
| 1.8 | 4.5 | 9 | 6419 |
| 1.6 | 4 | 8 | 7123 |
| 1.4 | 3.5 | 7 | 8588 |
| 1.2 | 3 | 6 | 10121 |
| 1 | 2.5 | 5 | 12566 |
| 0.9 | 2.25 | 4.5 | 16247 |
| 0.8 | 2 | 4 | 18974 |
| 0.7 | 1.75 | 3.5 | 19203 |
| 0.6 | 1.5 | 3 | 31403 |
| 0.5 | 1.25 | 2.5 | 41788 |
| 0.4 | 1 | 2 | 65727 |
| 0.35 | 0.875 | 1.75 | 78506 |
| 0.3 | 0.75 | 1.5 | 108017 |
| 0.25 | 0.625 | 1.25 | 150643 |
| 0.2 | 0.5 | 1 | 225197 |

Table 1. Mesh settings used in the cervix mesh size convergence study, with the canal size applied to the cervical canal face, the face size applied to the proximal cervix face, and the element size applied across the cervical geometry, though local refinement in element size was allowed. The number of cervix elements is the resulting number of elements in the cervix with the corresponding mesh settings.

| Element size | Number of fetal membrane elements |
|---|---|
| 2.5 | 1108 |
| 2 | 1843 |
| 1.75 | 2283 |
| 1.5 | 3107 |

Table 2. Mesh settings used in the fetal membrane mesh size convergence study with the resulting number of elements in the fetal membrane. The element size was applied to the fetal membrane face, and the element thickness in all cases was assigned as 1.

| Number of cervix elements | Computation time (min) |
|---|---|
| 6419 | 50 |
| 7123 | 59 |
| 8588 | 45 |
| 10121 | 53 |
| 12566 | 59 |
| 16247 | 62 |
| 18974 | 72 |
| 19203 | 76 |
| 31403 | 92 |
| 41788 | 109 |
| 65727 | 219 |
| 78506 | 167 |
| 108017 | 238 |
| 150643 | 383 |
| 225197 | 636 |

Table 3. Computation run time to model completion with changing number of elements in the cervix.

| Number of fetal membrane elements | Computation time (min) |
|---|---|
| 1108 | 226 |
| 1843 | 238 |
| 2283 | 316 |
| 3107 | 372 |

Table 4. Computation run time to model completion with changing number of elements in the fetal membrane.
