## Appendix C for "A finite element model of pregnancy derived from maternal sonography: effect of uterine and cervical structural properties on cervical mechanical loading"

### Appendix C: Proximal cervix stretch with varied material property values and maternal sonographic dimensions

#### LIST OF FIGURES


1 Proximal cervix heat maps of actual and used median geometry for stretch and difference from used median geometry. . . . 3
2 Proximal cervix heat maps of varying cervix $E$ for stretch and difference from 1.0 kPa. 4
3 Proximal cervix heat of maps varying cervix $\nu$ for stretch and difference from 0.225. 5
4 Proximal cervix heat maps of varying cervix $\xi$ for stretch and difference from 5 kPa. . 6
5 Proximal cervix heat maps of varying cervix $\alpha$ for stretch and difference from 3.00. . 7
6 Proximal cervix heat maps of varying uterus $E$ for stretch and difference from 1.5 kPa. 8
7 Proximal cervix heat maps of varying uterus $\nu$ for stretch and difference from 0.27. . 9
8 Proximal cervix heat maps of varying uterus $\xi$ for stretch and difference from 2.7 kPa. 10
9 Proximal cervix heat maps of varying uterus $\alpha$ for stretch and difference from 0.74. . 11
10 Proximal cervix heat maps of varying UD1 for stretch and difference from median (120.70 mm). . . . 12
11 Proximal cervix heat maps of varying UD2 for stretch and difference from median (28.95 mm). . . . 13
12 Proximal cervix heat maps of varying UD3 for stretch and difference from median (38.08 mm). . . . 14
13 Proximal cervix heat maps of varying UD3a for stretch and difference from median (33.32 mm). . . . 15
14 Proximal cervix heat maps of varying UD1a for stretch and difference from median (90.52 mm). . . . 16
15 Proximal cervix heat maps of varying UD1b for stretch and difference from median (30.17 mm). . . . 17
16 Proximal cervix heat maps of varying PCO for stretch and difference from median (22.01 mm). . . . 18
17 Proximal cervix heat maps of varying UT1 for stretch and difference from median (6.48 mm). . . . 19
18 Proximal cervix heat maps of varying UT2 for stretch and difference from median (5.76 mm). . . . 20
19 Proximal cervix heat maps of varying UT3 for stretch and difference from median (7.31 mm). . . . 21
20 Proximal cervix heat maps of varying CL for stretch and difference from median (32.31). . . . 22
21 Proximal cervix heat maps of varying CD1 for stretch and difference from median (33.43 mm). . . . 23
22 Proximal cervix heat maps of varying CD2 for stretch and difference from median (3.11 mm). . . . 24
23 Proximal cervix heat maps of varying AUCA for stretch and difference from median (81.18°). . . . 25
24 Proximal cervix heat maps of varying FM thickness for stretch and difference from used value (0.79 mm). . . . 26

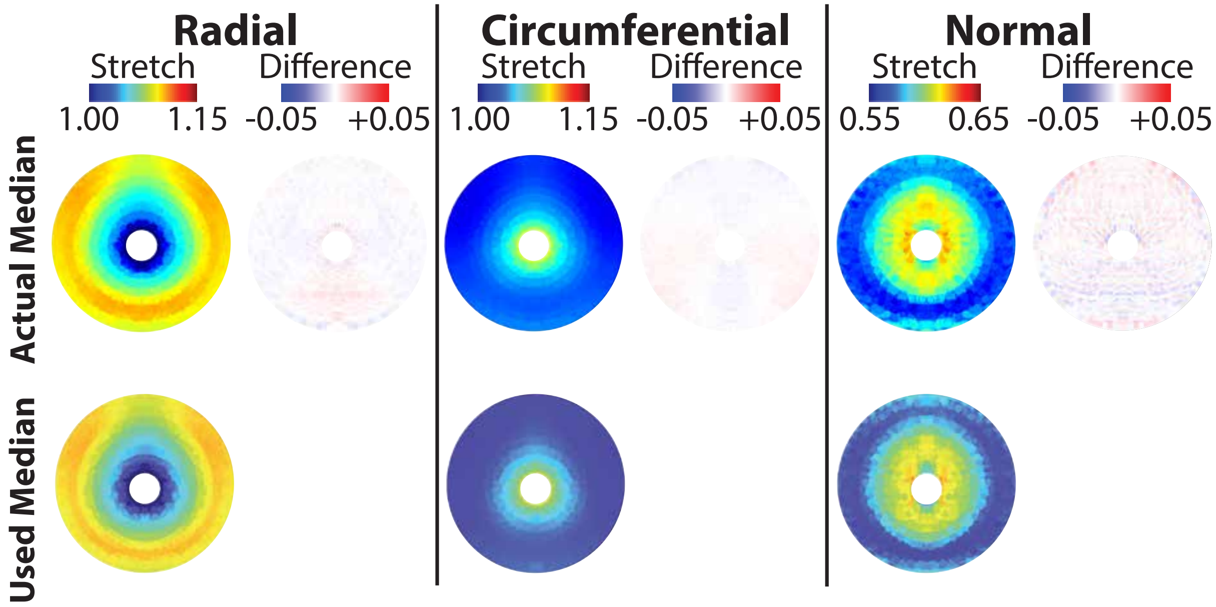


Fig. 1. Proximal cervix heat maps of actual and used median geometry for stretch and difference from used median geometry.

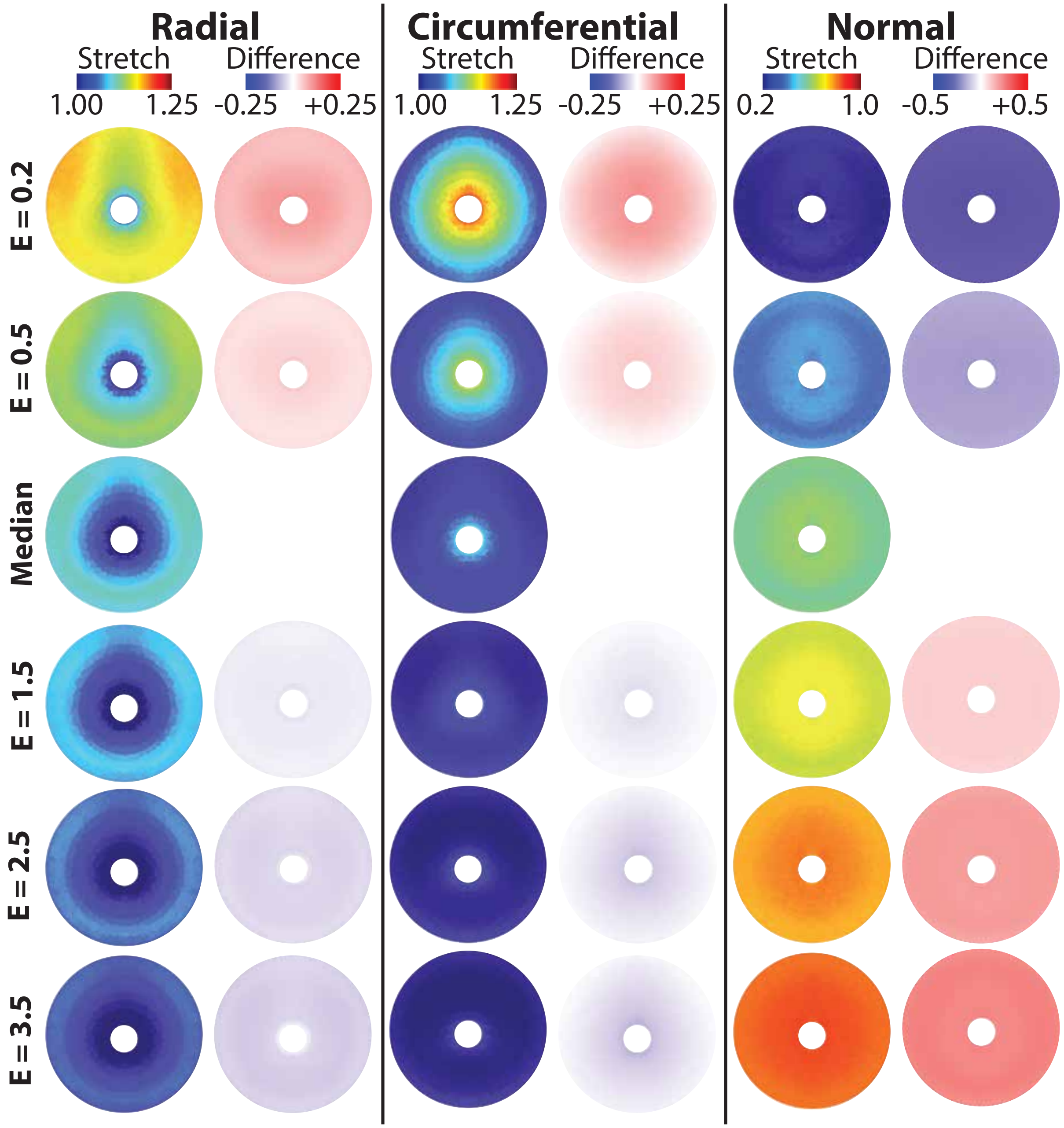


Fig. 2. Proximal cervix heat maps of varying cervix $E$ for stretch and difference from 1.0 kPa.

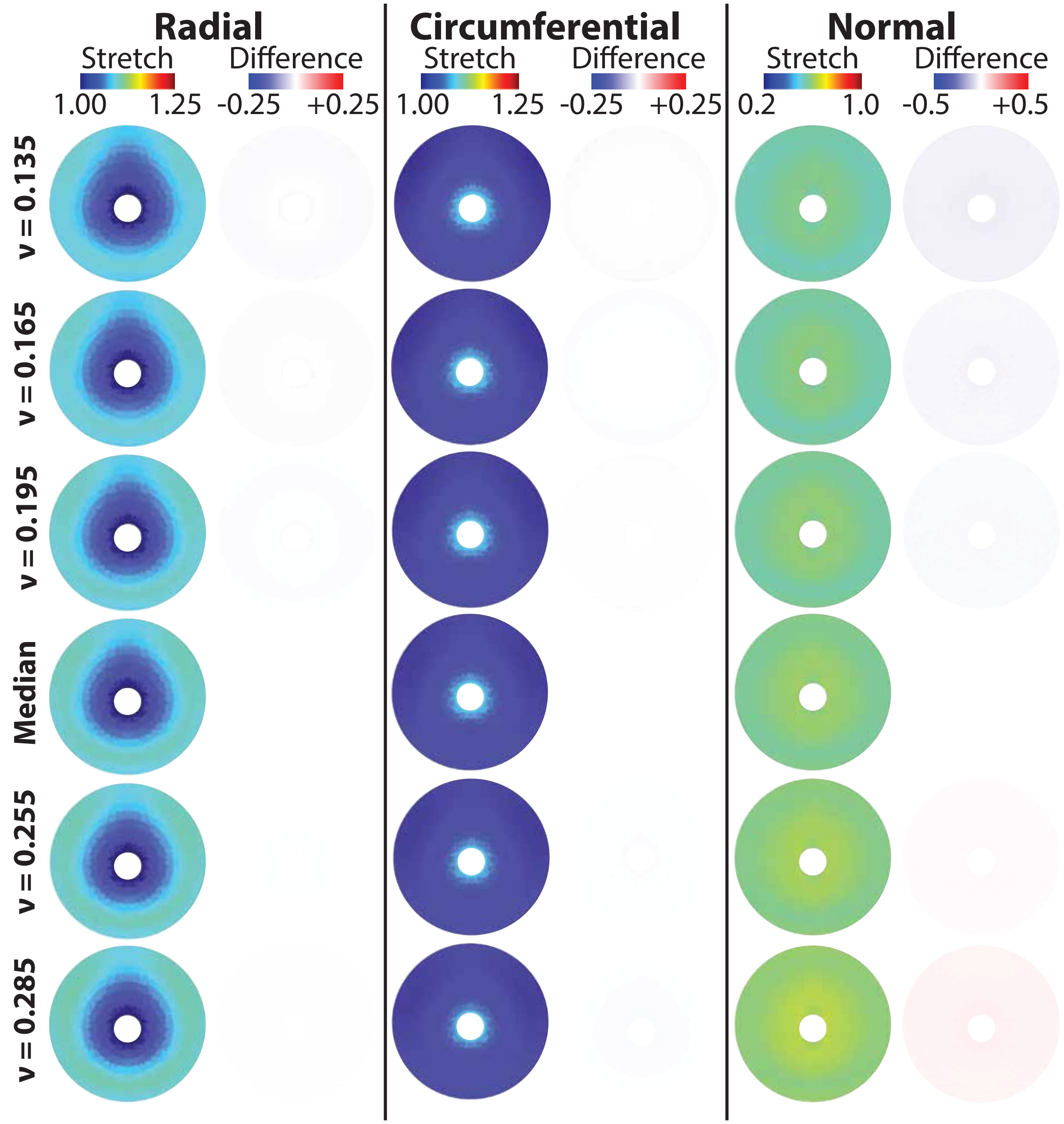


Fig. 3. Proximal cervix heat of maps varying cervix $\nu$ for stretch and difference from 0.225.

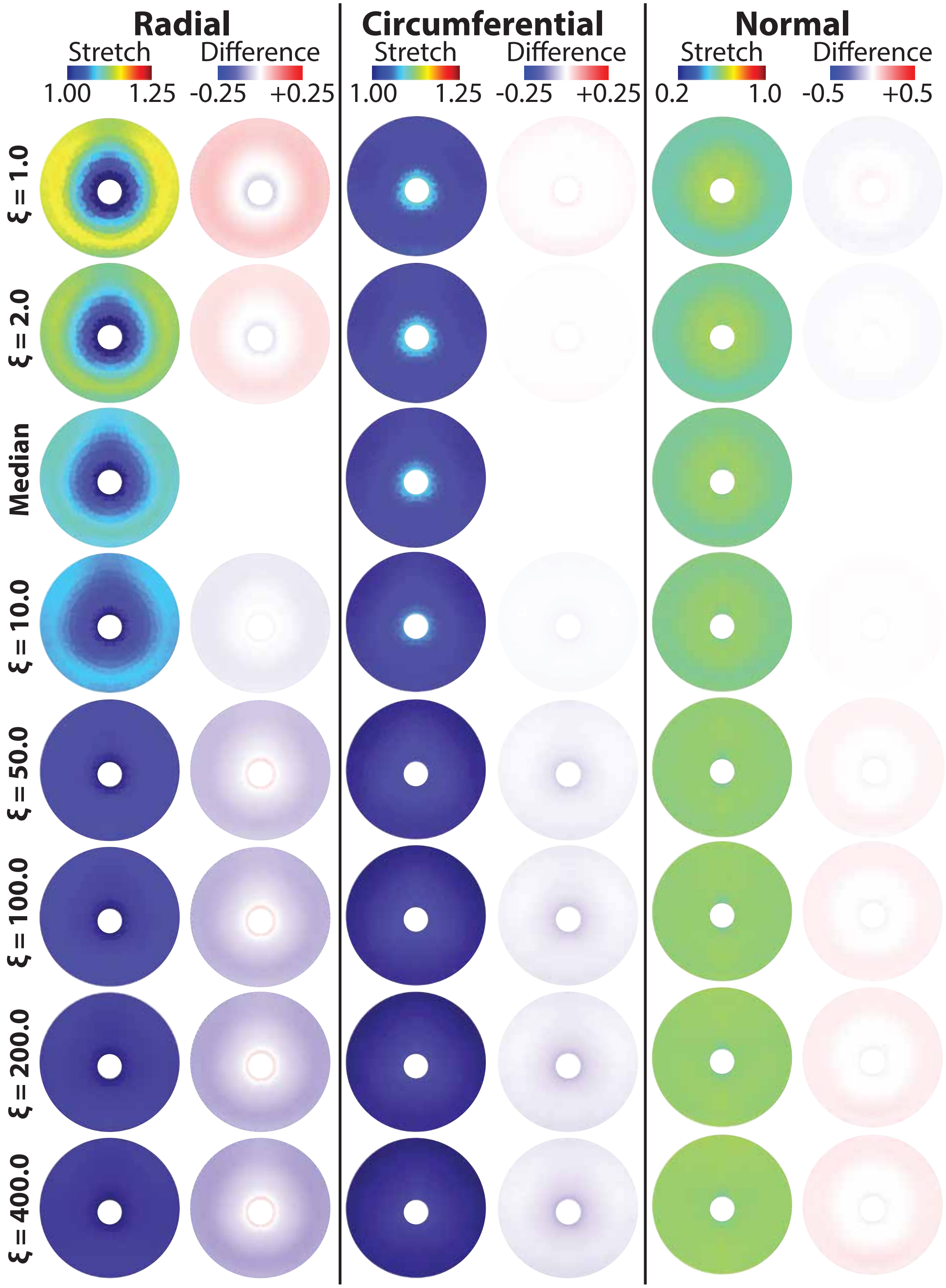


Fig. 4. Proximal cervix heat maps of varying cervix $\xi$ for stretch and difference from 5 kPa.

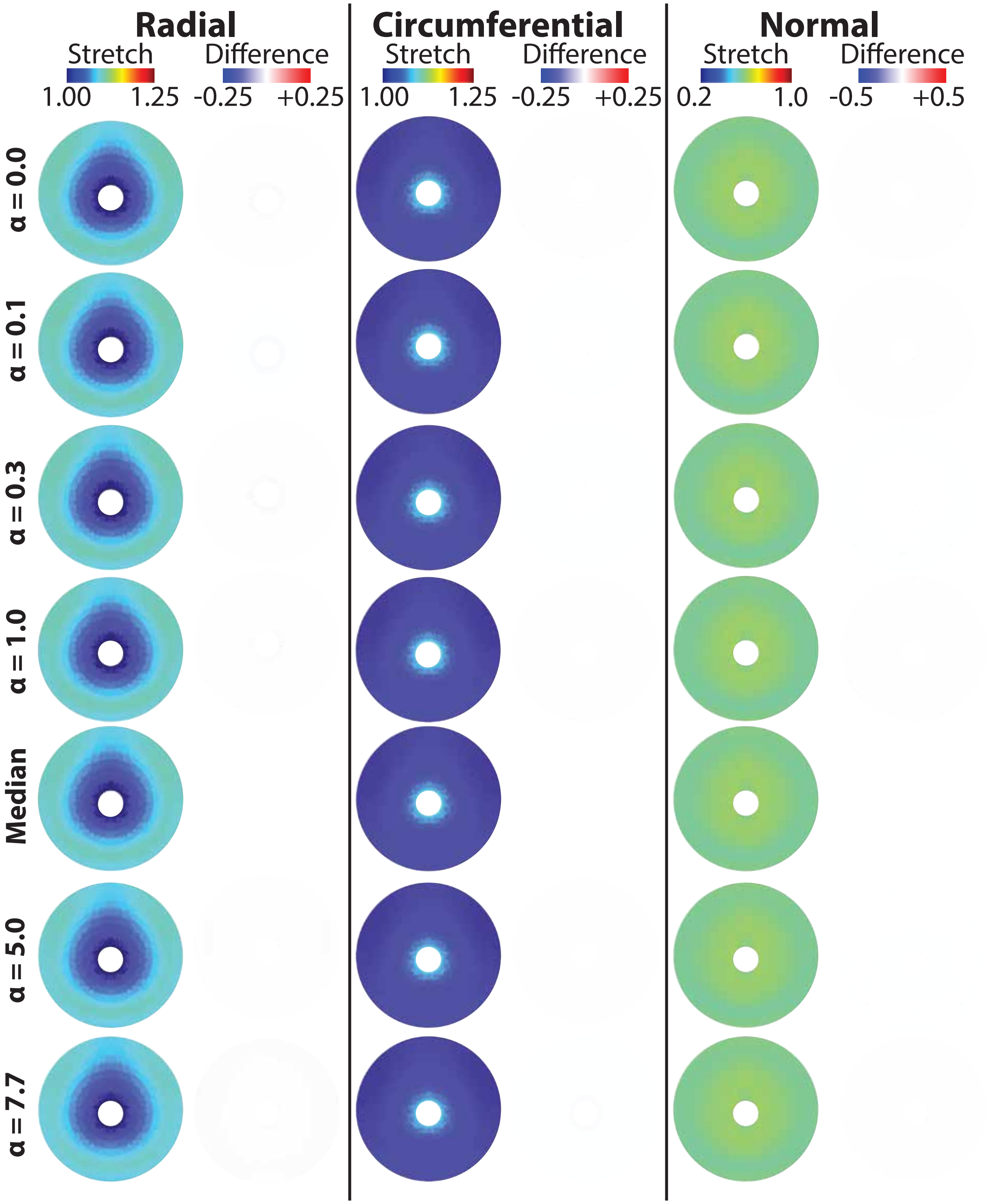


Fig. 5. Proximal cervix heat maps of varying cervix $\alpha$ for stretch and difference from 3.00.

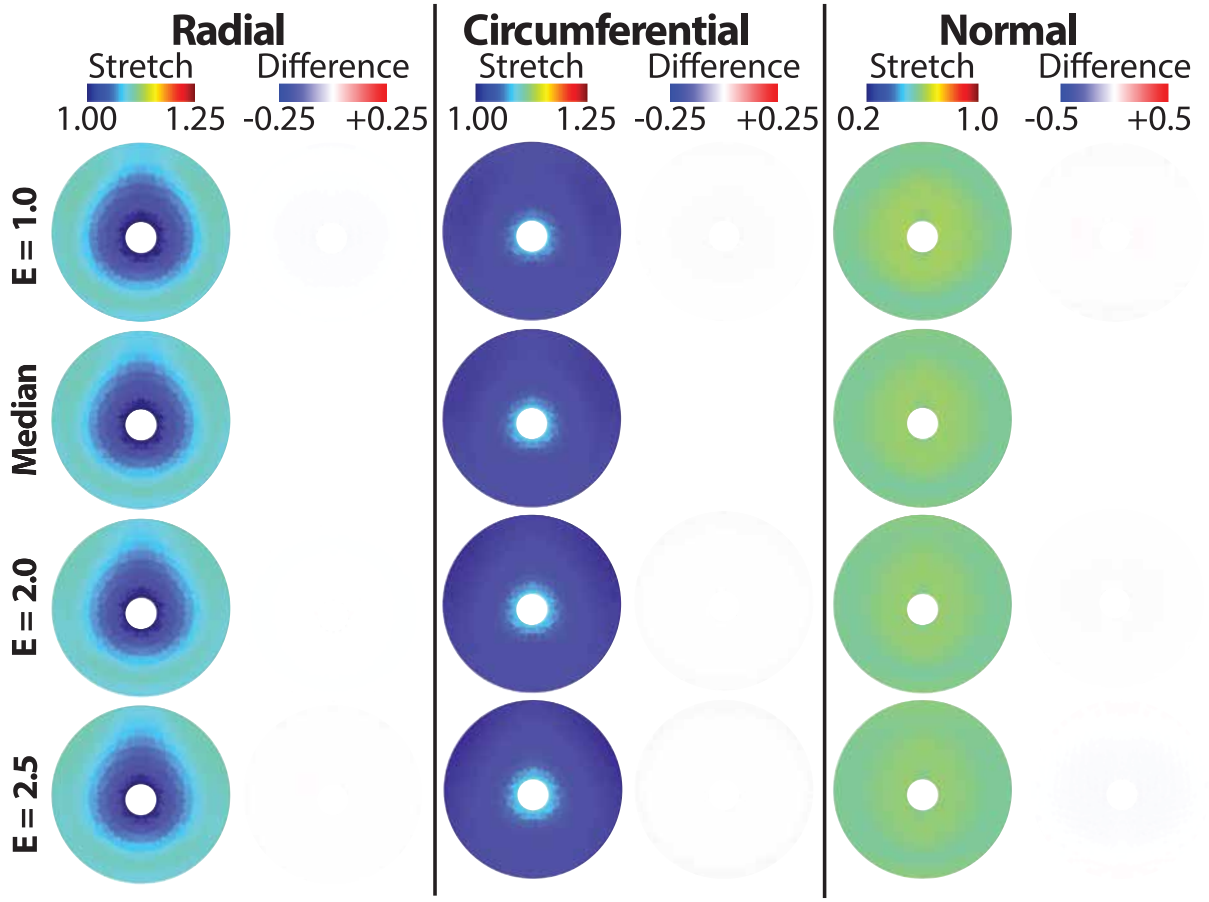


Fig. 6. Proximal cervix heat maps of varying uterus $E$ for stretch and difference from 1.5 kPa.

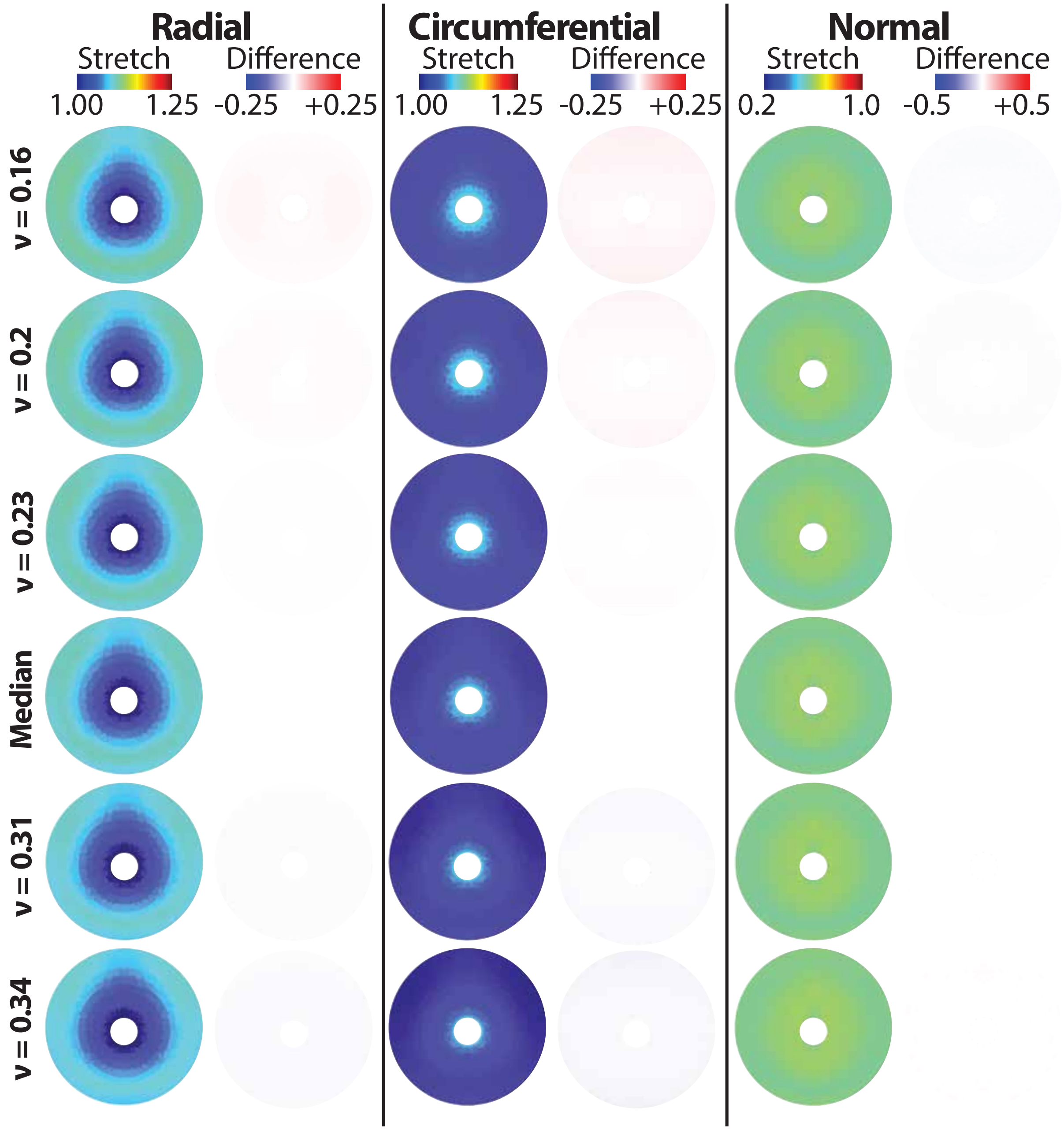


Fig. 7. Proximal cervix heat maps of varying uterus $\nu$ for stretch and difference from 0.27.

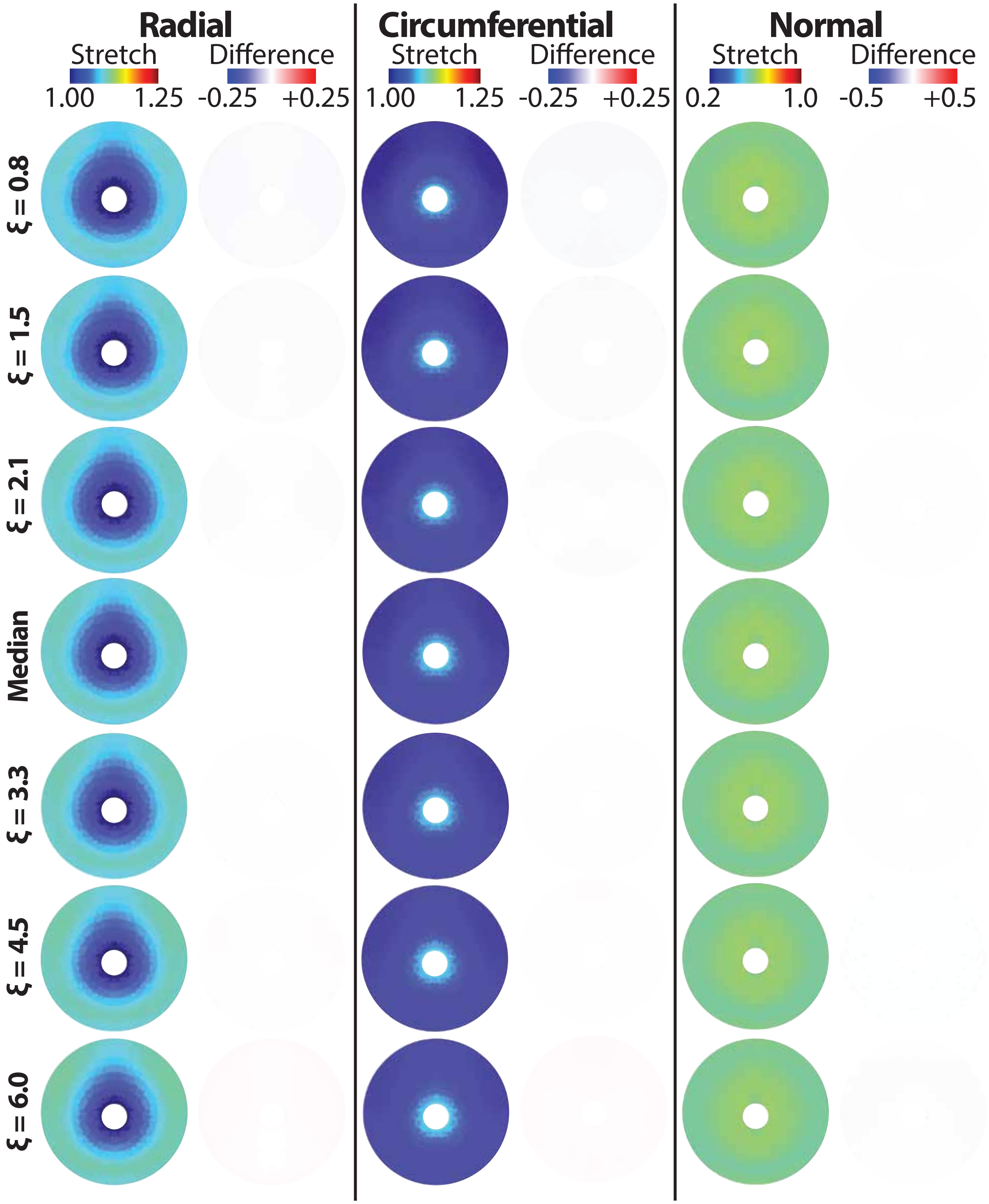


Fig. 8. Proximal cervix heat maps of varying uterus $\xi$ for stretch and difference from 2.7 kPa.

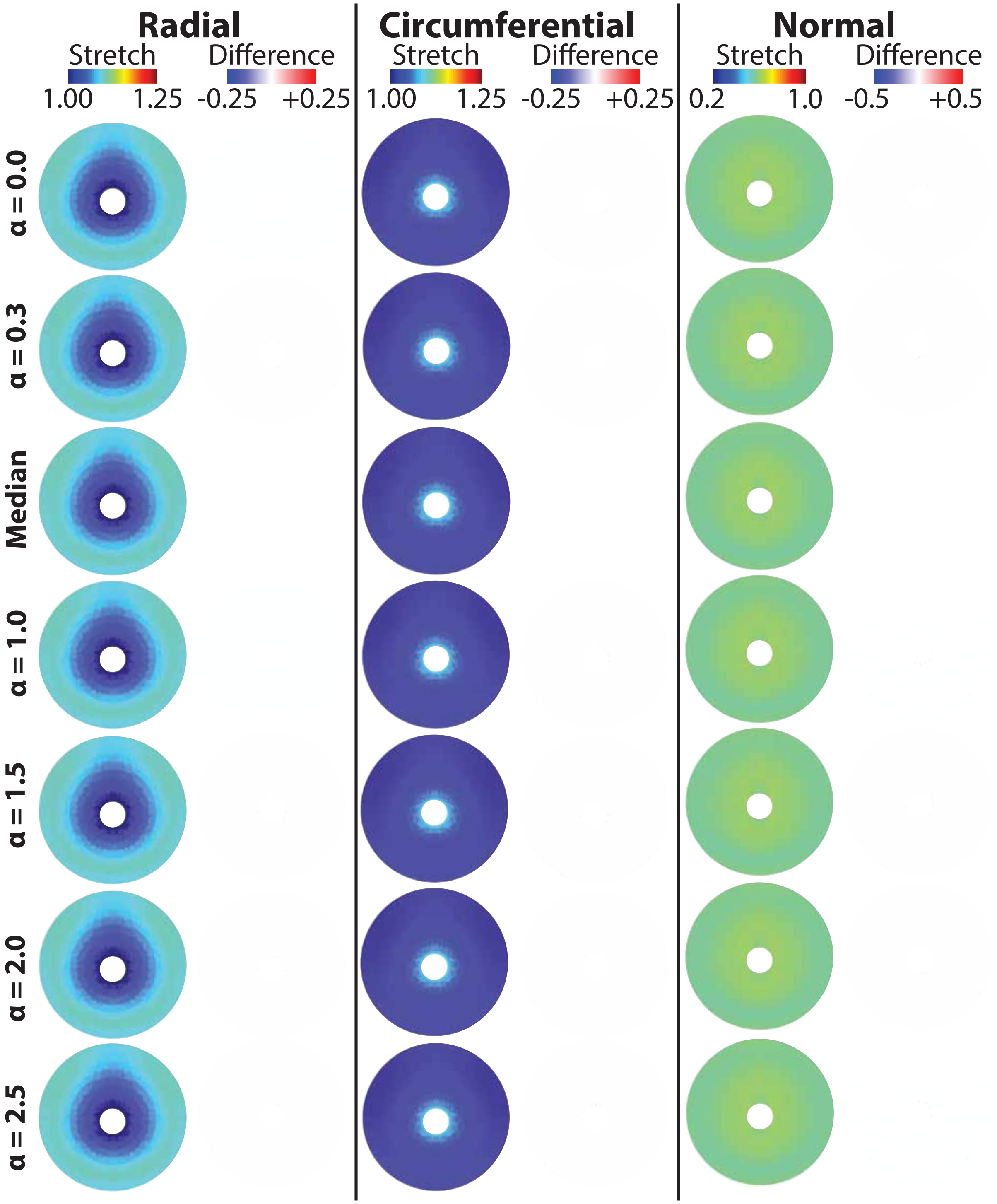


Fig. 9. Proximal cervix heat maps of varying uterus $\alpha$ for stretch and difference from 0.74.

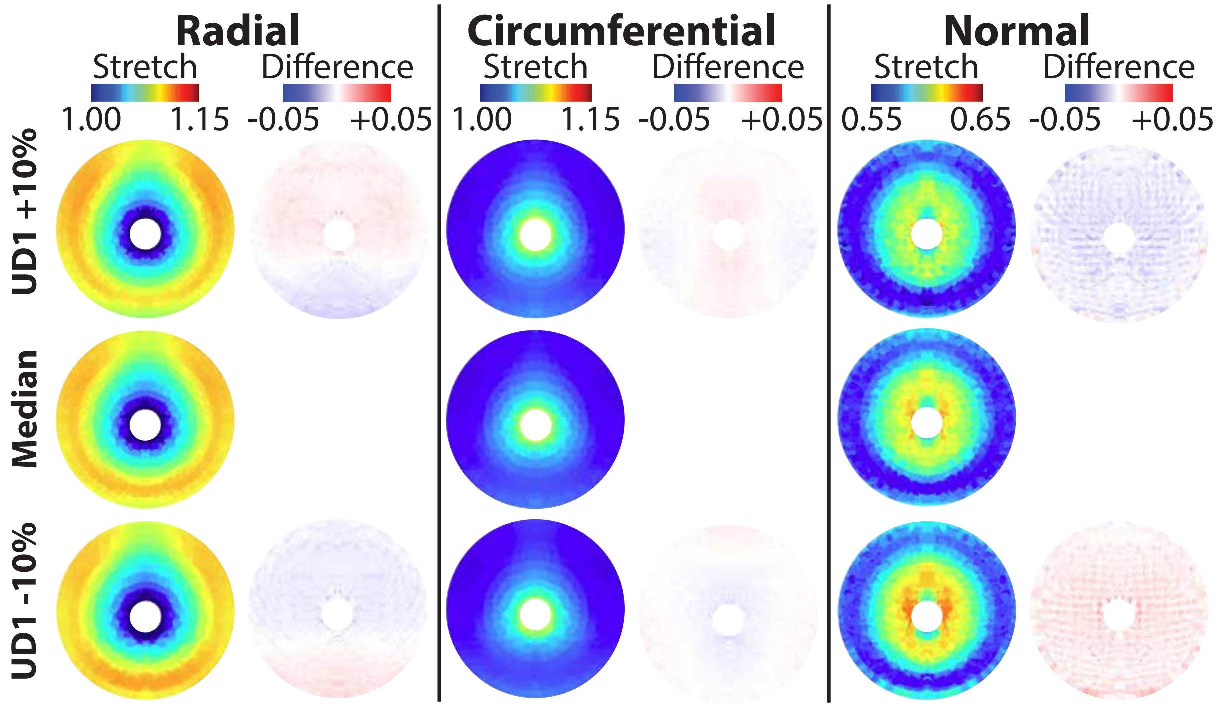


Fig. 10. Proximal cervix heat maps of varying UD1 for stretch and difference from median (120.70 mm).

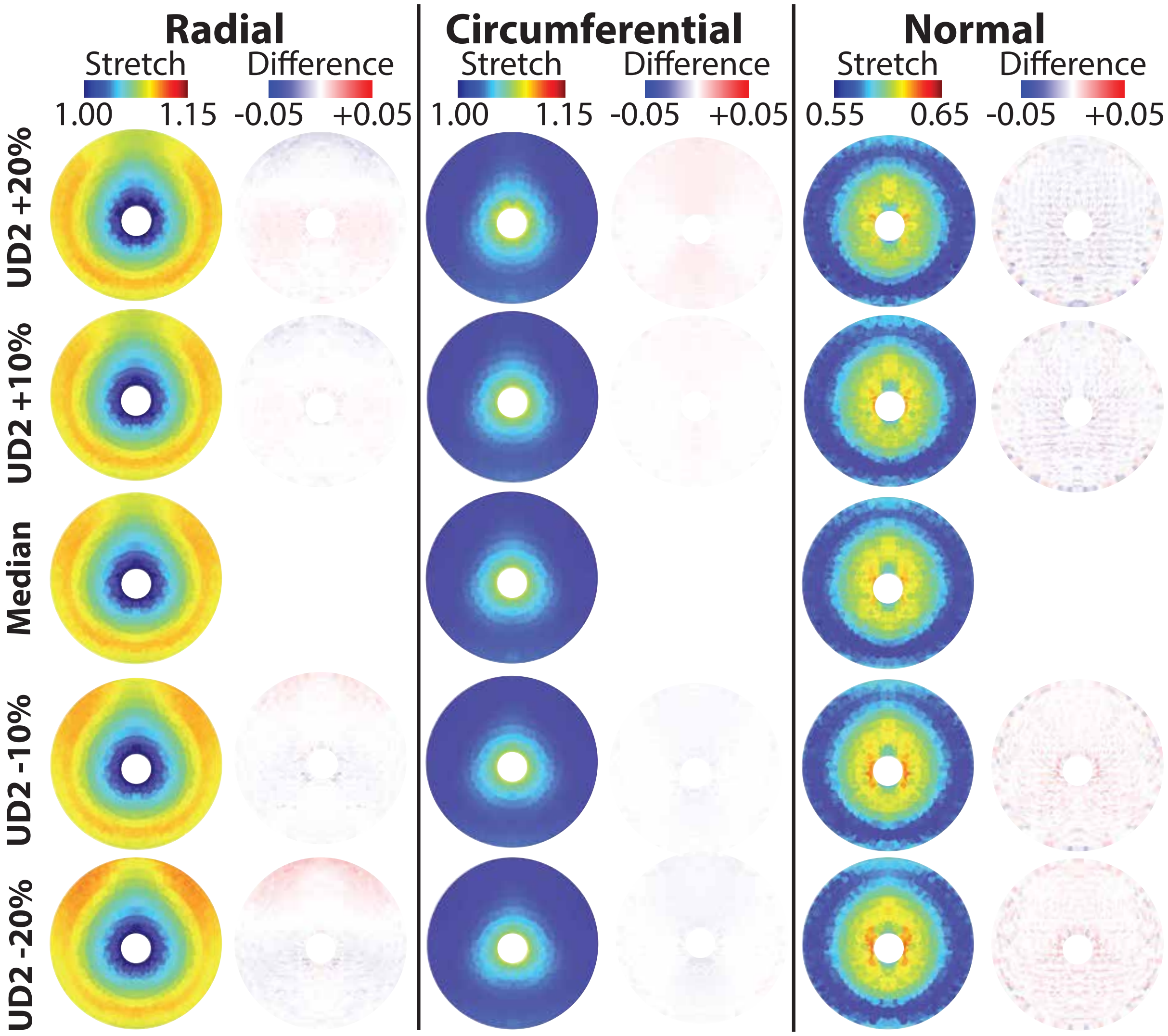


Fig. 11. Proximal cervix heat maps of varying UD2 for stretch and difference from median (28.95 mm).

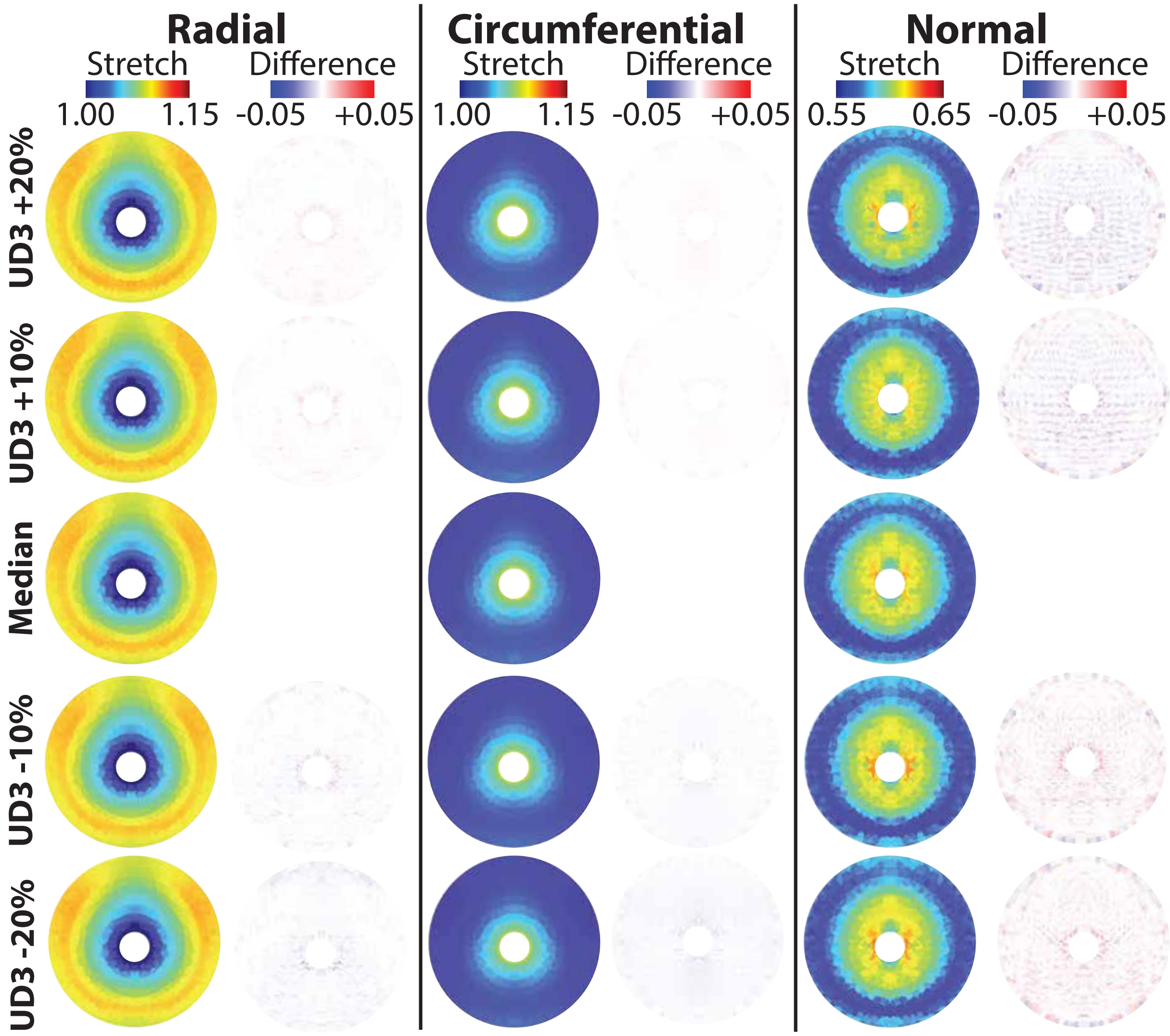


Fig. 12. Proximal cervix heat maps of varying UD3 for stretch and difference from median (38.08 mm).

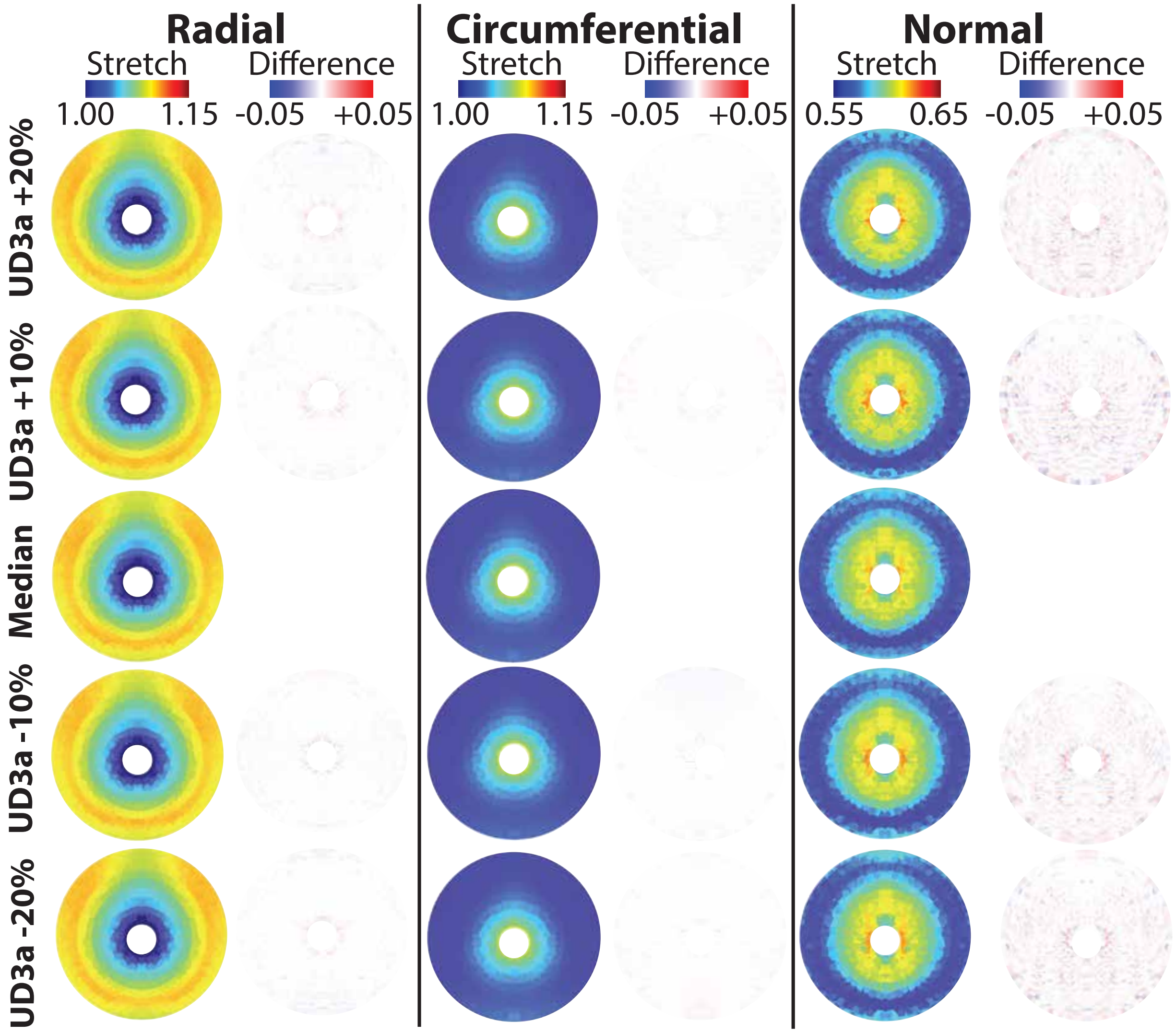


Fig. 13. Proximal cervix heat maps of varying UD3a for stretch and difference from median (33.32 mm).

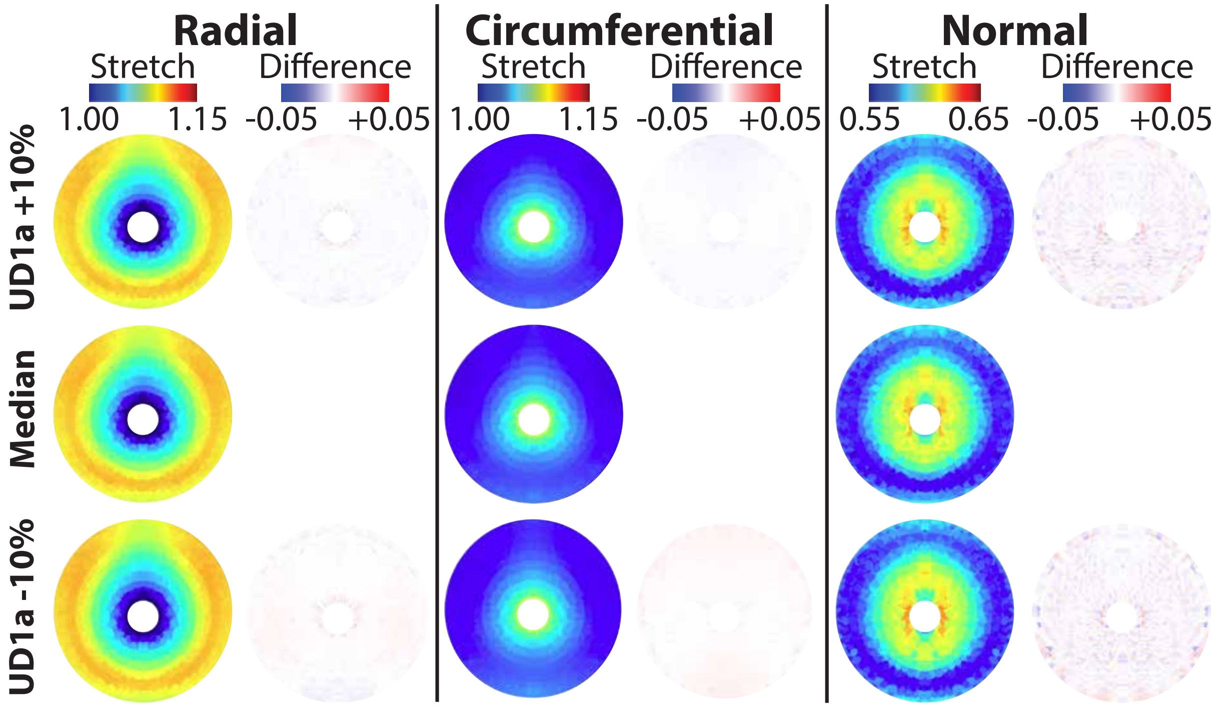


Fig. 14. Proximal cervix heat maps of varying UD1a for stretch and difference from median (90.52 mm).

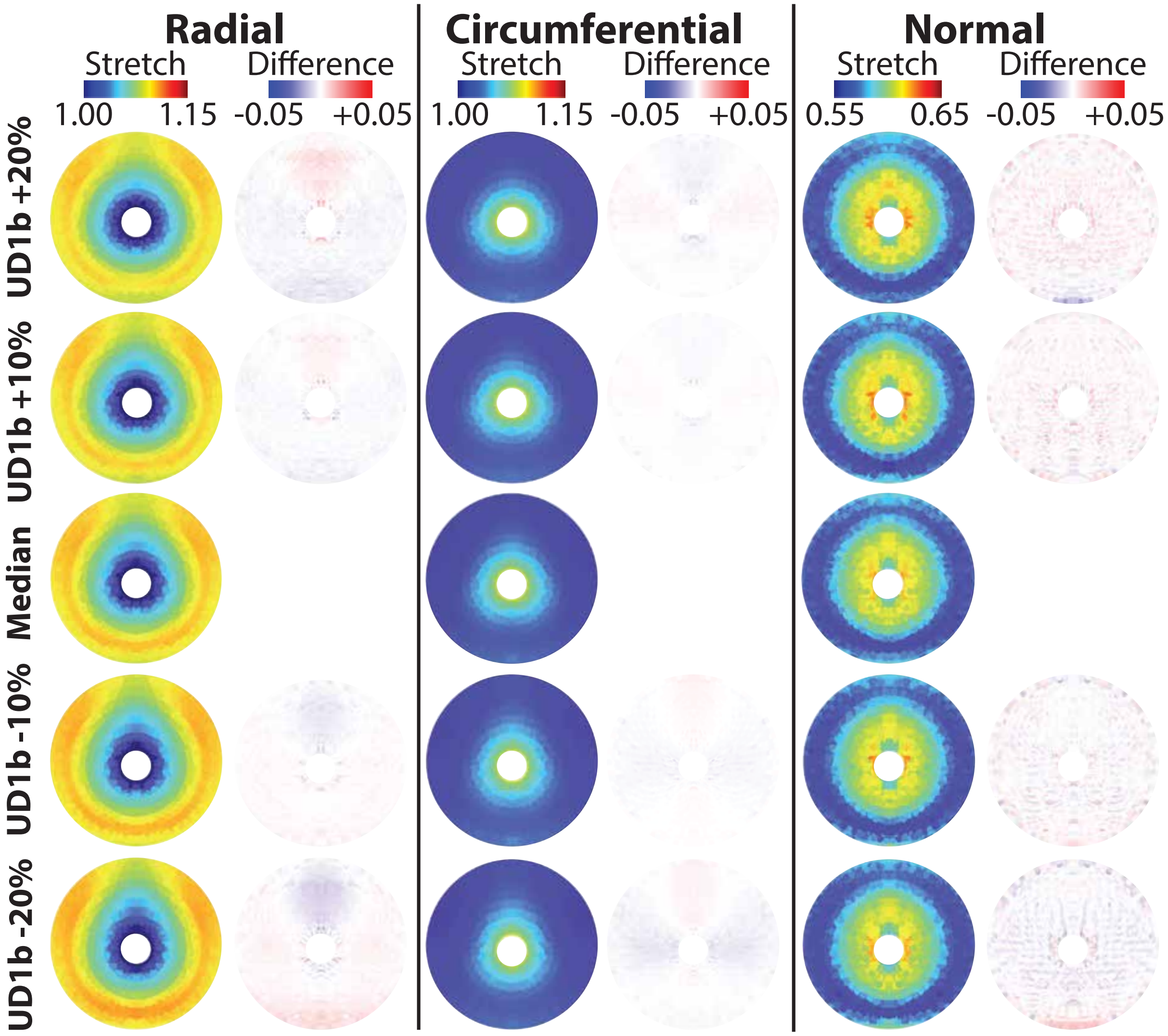


Fig. 15. Proximal cervix heat maps of varying UD1b for stretch and difference from median (30.17 mm).

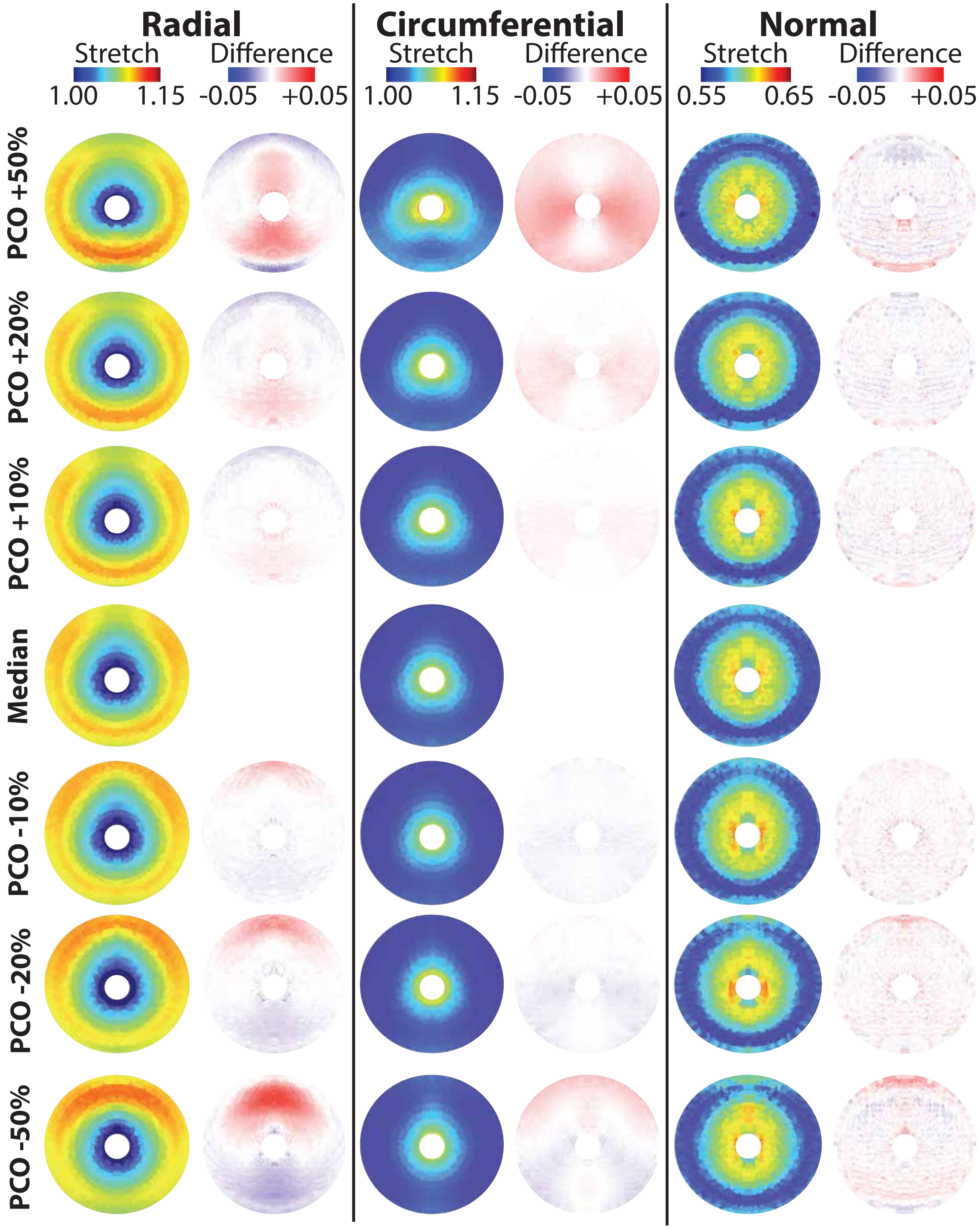


Fig. 16. Proximal cervix heat maps of varying PCO for stretch and difference from median (22.01 mm).

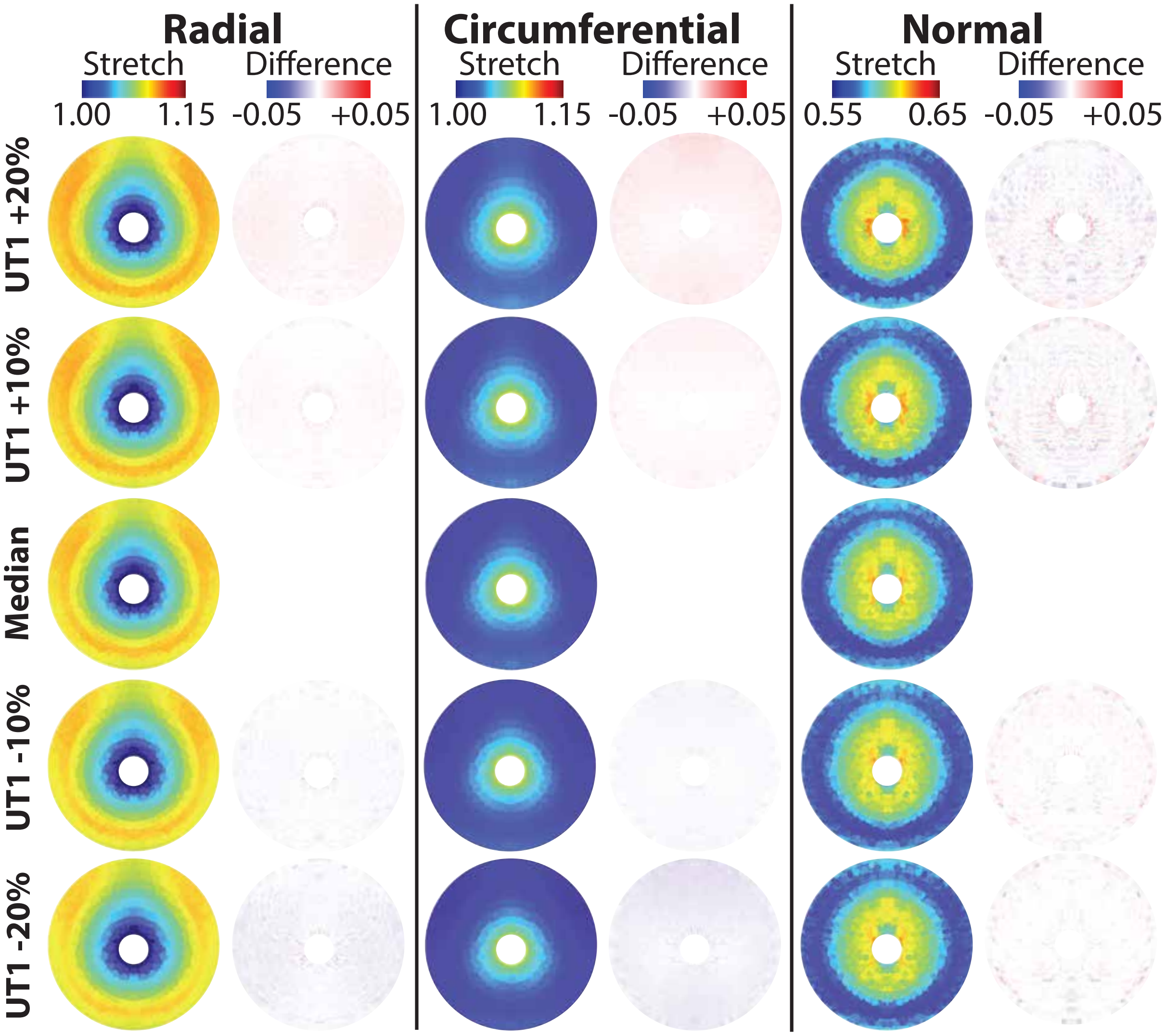


Fig. 17. Proximal cervix heat maps of varying UT1 for stretch and difference from median (6.48 mm).

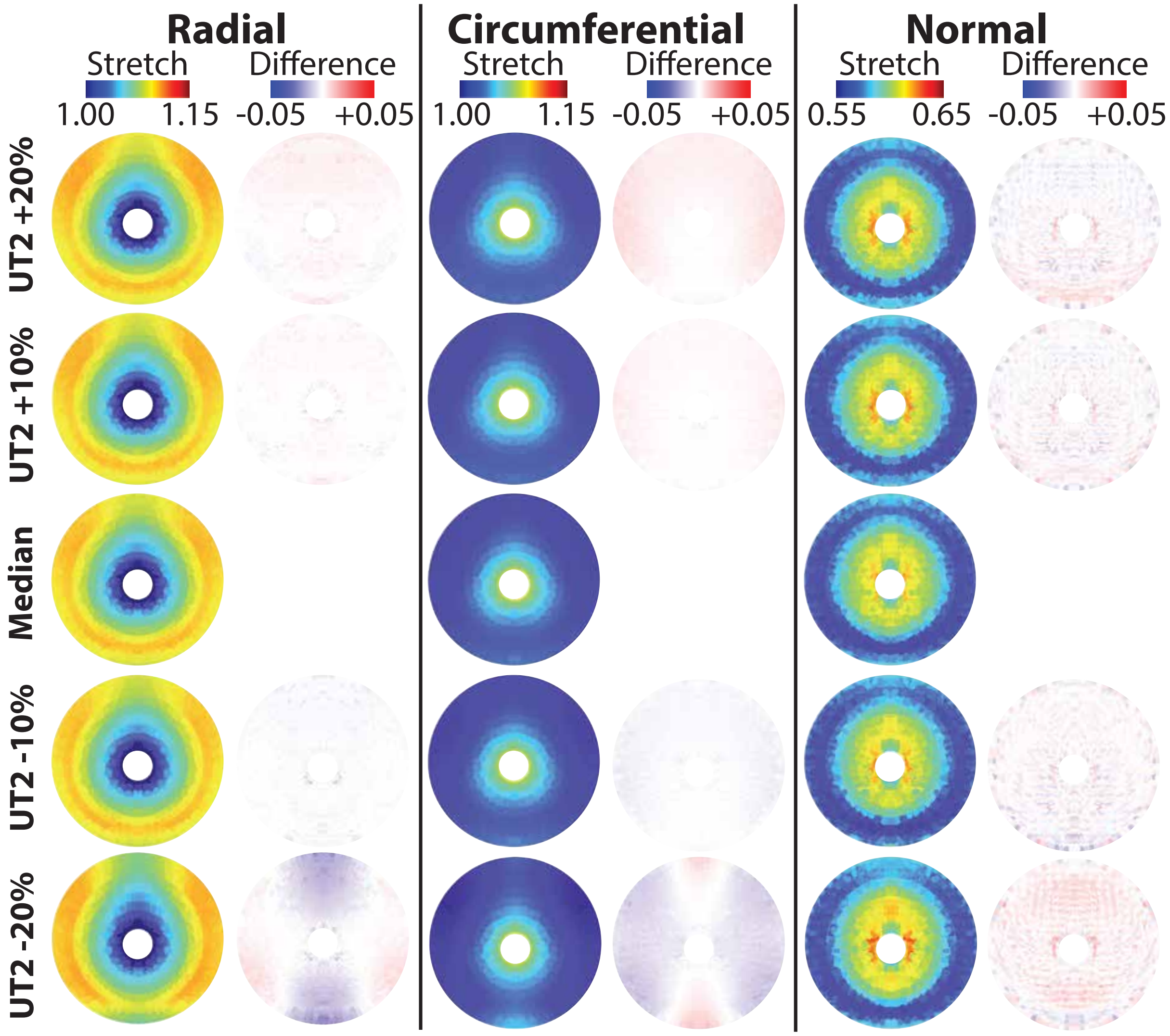


Fig. 18. Proximal cervix heat maps of varying UT2 for stretch and difference from median (5.76 mm).

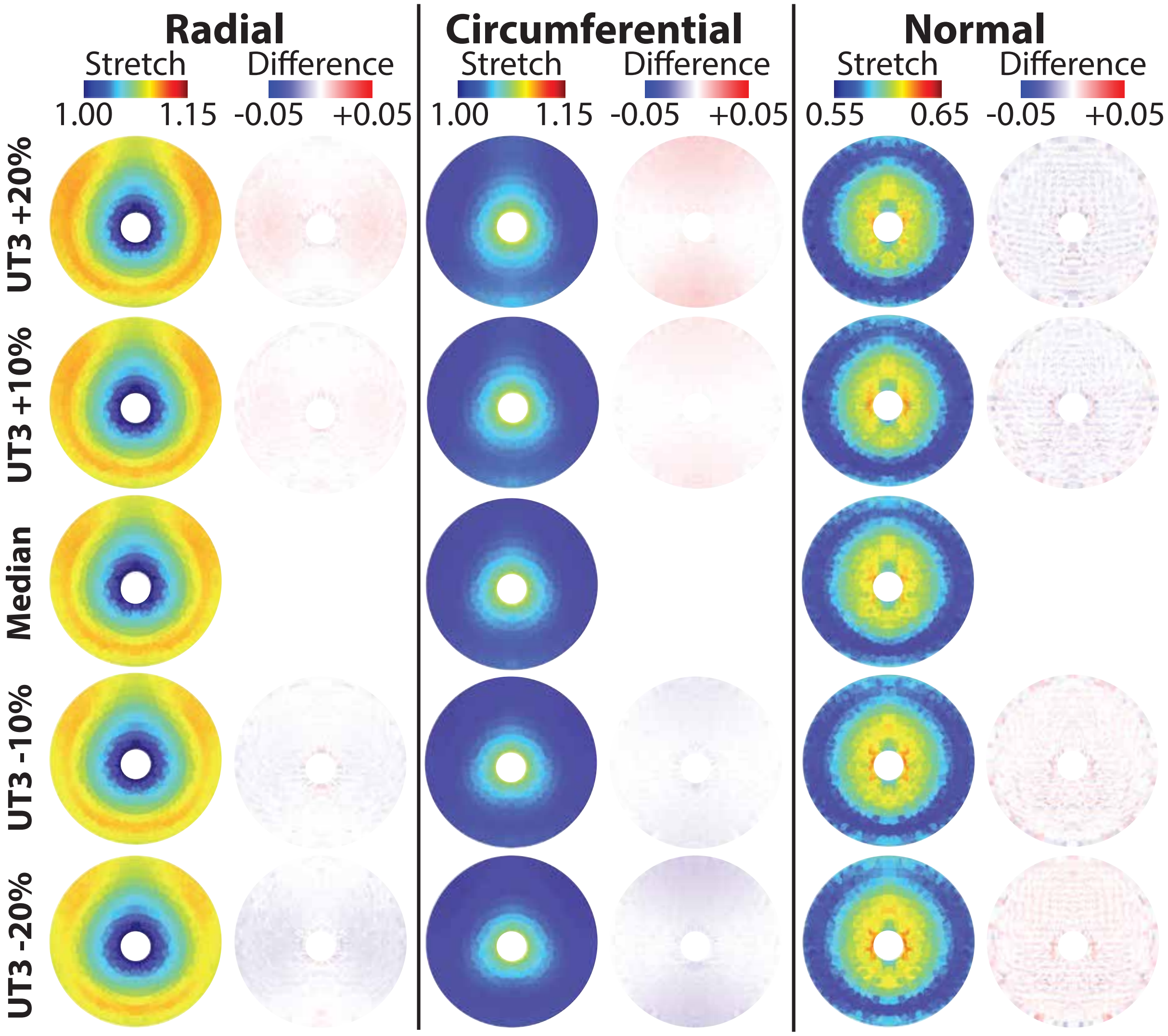


Fig. 19. Proximal cervix heat maps of varying UT3 for stretch and difference from median (7.31 mm).

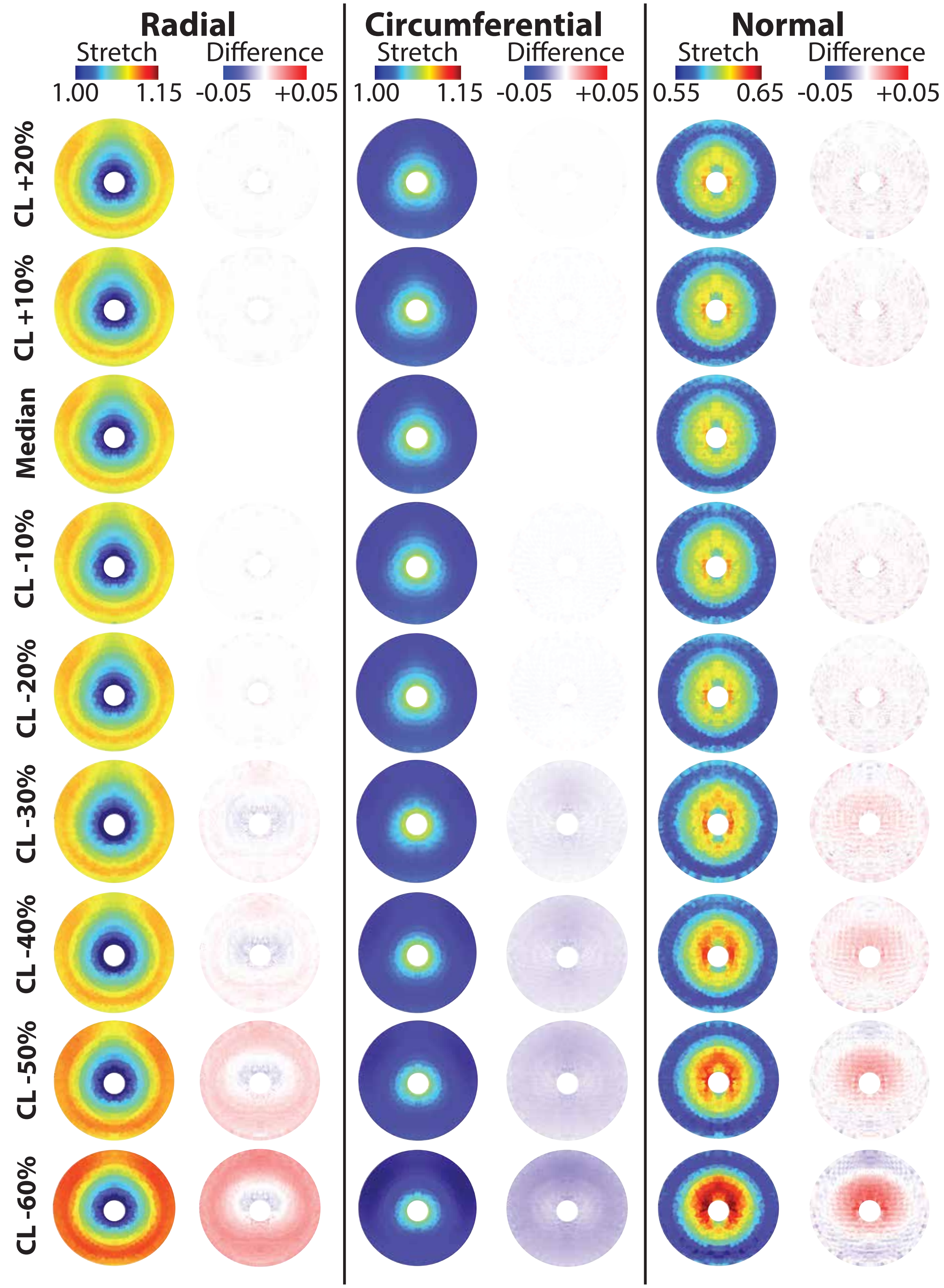


Fig. 20. Proximal cervix heat maps of varying CL for stretch and difference from median (32.31).

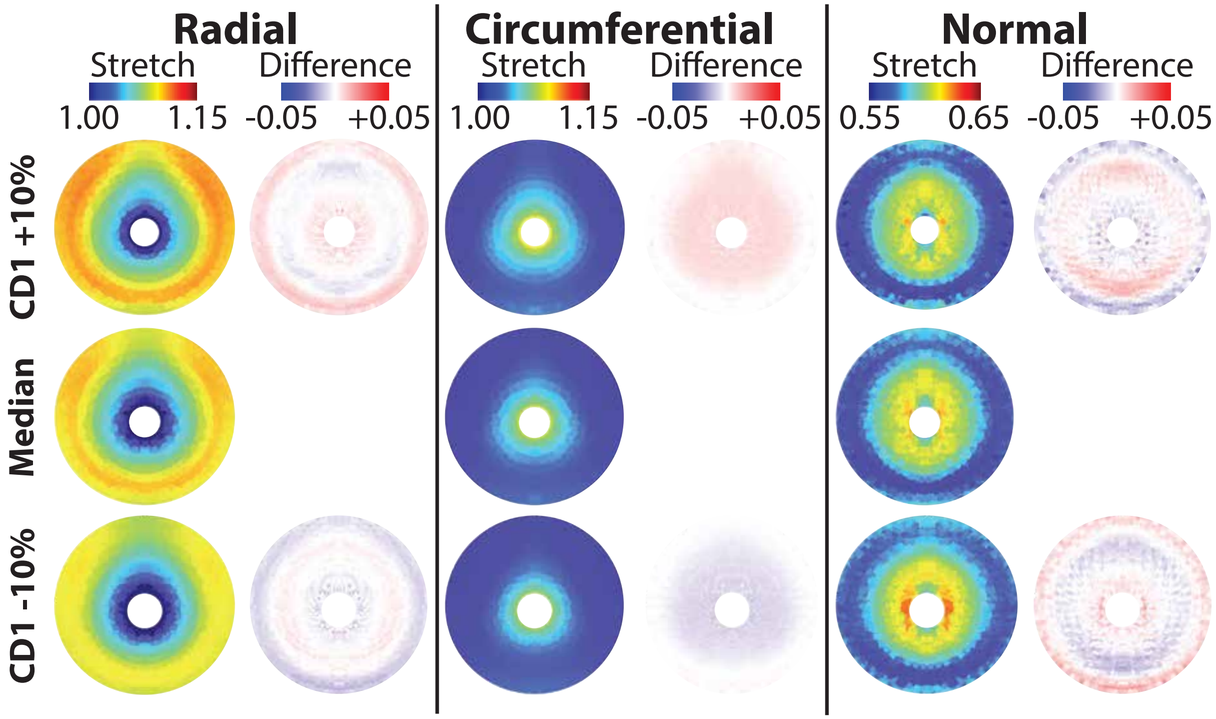


Fig. 21. Proximal cervix heat maps of varying CD1 for stretch and difference from median (33.43 mm).

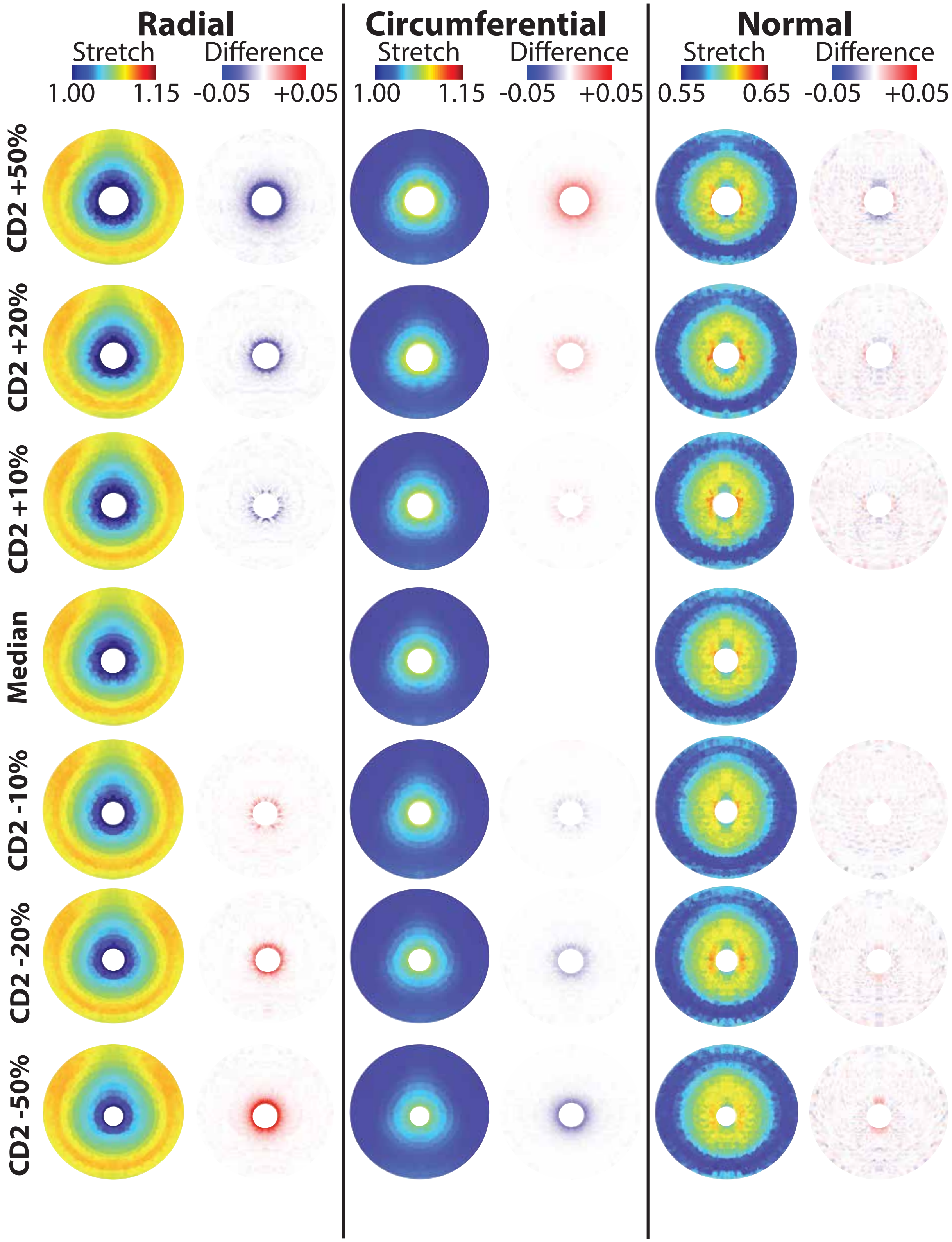


Fig. 22. Proximal cervix heat maps of varying CD2 for stretch and difference from median (3.11 mm).

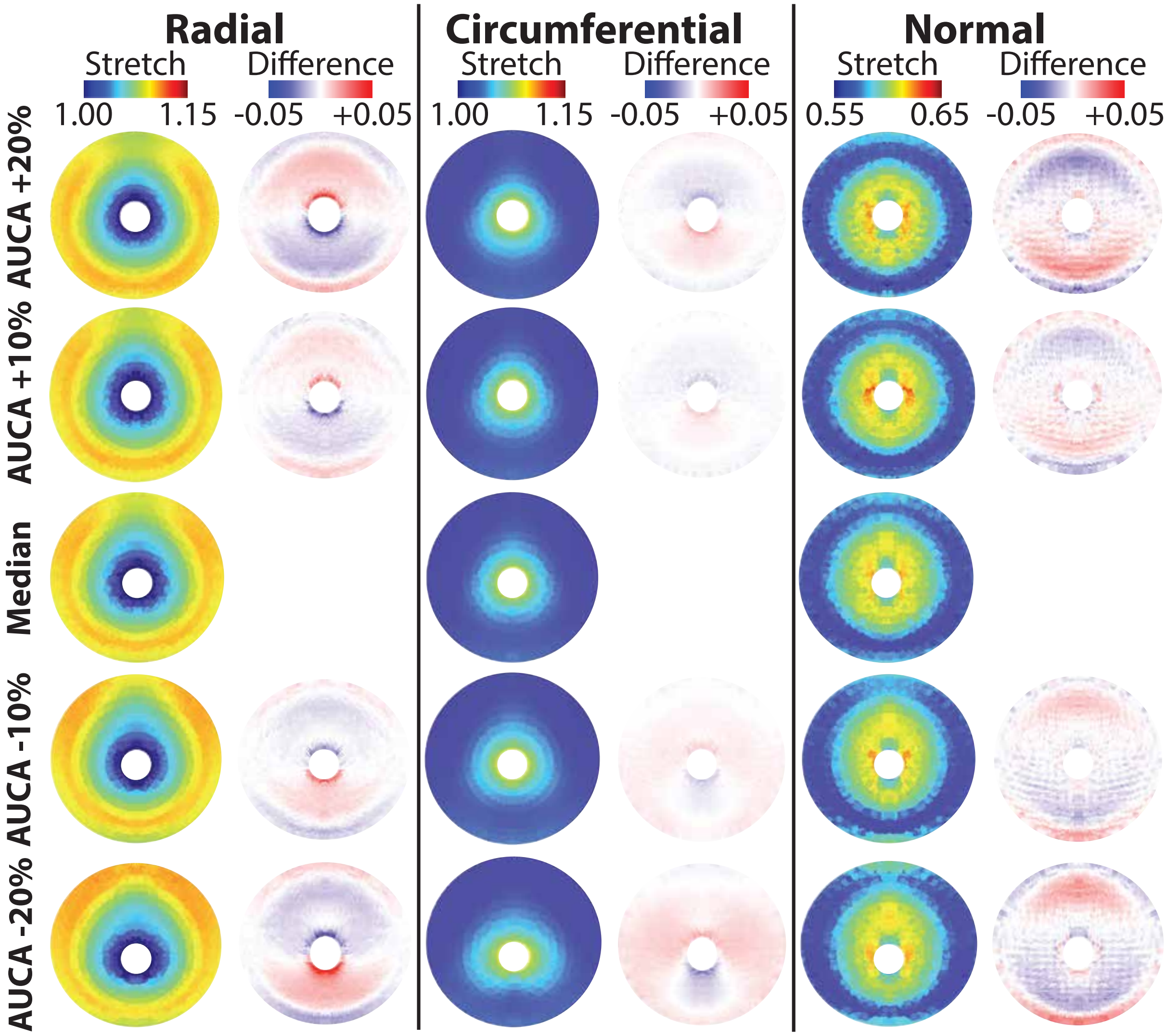


Fig. 23. Proximal cervix heat maps of varying AUCA for stretch and difference from median (81.18°).

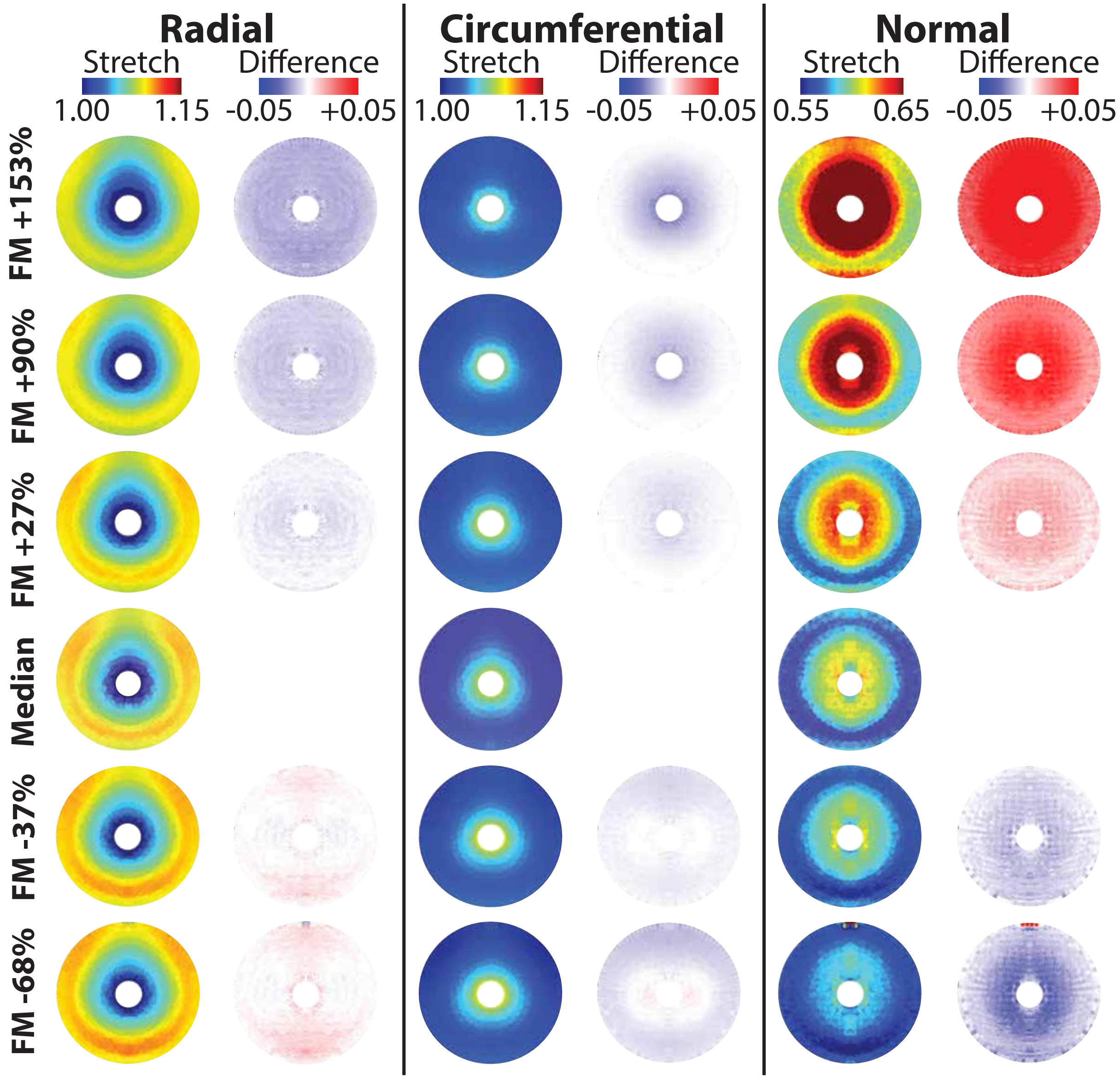


Fig. 24. Proximal cervix heat maps of varying FM thickness for stretch and difference from used value (0.79 mm).

#### LIST OF TABLES

1 Slope and $R^2$ of linear fits to stretch against the normalized material property value, where normalized material property value indicates value's position between the lower and upper bound tested for the given material property (Tab. 4). Stretch was analyzed in the anterior, left, and posterior ROIs of the proximal cervix mid-stroma in the surface radial, circumferential, and normal directions. The maximum first principal stretch ($\lambda_1$) was analyzed in the cervical internal os (IO) and in the fetal membrane (FM). Asterisks indicate significance level: *P* $>$.05 ( ), *P* = .01-.05 (*), *P* = .001-.01(**), *P* $<$.001 (***). Level of slope significance is based on the *t*-statistic where $t = m/\text{SE}(m)$, where $m$ is the slope and SE$(m)$ is the standard error of the slope. The *P*-value does not necessarily give significance of slope value magnitude from 0. . . . . 28

2 Slope and $R^2$ of linear fits to stretch against percent change in ultrasonic dimensions, where percent change indicates the percentage change from the median ultrasonic dimension value (Tab. 5). Stretch was analyzed in the anterior, left, and posterior ROIs of the proximal cervix mid-stroma in the surface radial, circumferential, and normal direction. The maximum first principal stretch ($\lambda_1$) was analyzed in the cervical internal os (IO) and in the fetal membrane (FM). Asterisks indicate significance level: *P* $>$.05 ( ), *P* = .01-.05 (*), *P* = .001-.01(**), *P* $<$.001 (***). Level of slope significance is based on the *t*-statistic where $t = m/\text{SE}(m)$, where $m$ is the slope and SE$(m)$ is the standard error of the slope. The *P*-value does not necessarily give significance of slope value magnitude from 0. . . . . 29

Table 1. Slope and $R^2$ of linear fits to stretch against the normalized material property value, where normalized material property value indicates value's position between the lower and upper bound tested for the given material property (Tab. 4). Stretch was analyzed in the anterior, left, and posterior ROIs of the proximal cervix mid-stroma in the surface radial, circumferential, and normal directions. The maximum first principal stretch ($\lambda_1$) was analyzed in the cervical internal os (IO) and in the fetal membrane (FM). Asterisks indicate significance level: $P >$.05 ( ), $P$ = .01-.05 (*), $P$ = .001-.01(**), $P <$.001 (***). Level of slope significance is based on the $t$-statistic where $t = m/\mathrm{SE}(m)$, where $m$ is the slope and $\mathrm{SE}(m)$ is the standard error of the slope. The $P$-value does not necessarily give significance of slope value magnitude from 0.

| Material property | Anterior stretch slope*$10^{-3}$ ($R^2$) | | | Left stretch slope*$10^{-3}$ ($R^2$) | | |
|---|---|---|---|---|---|---|
| Direction | Rad | Circ | Norm | Rad | Circ | Norm |
| Cervix E | -1.08* (0.73) | -1.09* (0.65) | 5.71* (0.80) | -1.21* (0.75) | -0.95* (0.64) | 5.88* (0.80) |
| Cervix $\nu$ | 0.09*** (1.00) | -0.02 (0.50) | 0.57*** (0.98) | 0.09*** (0.99) | -0.03** (0.88) | 0.60*** (0.99) |
| Cervix $\xi$ | -0.66* (0.46) | -0.23** (0.68) | 0.24* (0.51) | -0.83* (0.44) | -0.21** (0.71) | 0.31* (0.52) |
| Cervix $\alpha$ | -0.01** (0.83) | 0.00 (-0.20) | -0.00 (-0.20) | -0.02** (0.86) | 0.00 (-0.10) | 0.00* (0.53) |
| Uterus E | 0.04** (0.99) | 0.05* (0.88) | -0.10* (0.94) | 0.06* (0.95) | 0.07** (0.98) | -0.22* (0.91) |
| Uterus $\nu$ | -0.16*** (0.99) | -0.23*** (0.99) | 0.16*** (0.99) | -0.20*** (0.99) | -0.16*** (0.99) | 0.11*** (1.00) |
| Uterus $\xi$ | 0.10*** (0.99) | 0.15*** (0.987) | -0.10*** (0.98) | 0.13*** (0.97) | 0.10*** (0.99) | -0.10*** (0.98) |
| Uterus $\alpha$ | 0.00 (-0.20) | -0.00 (-0.20) | 0.00 (-0.19) | -0.00 (-0.20) | 0.00 (-0.20) | -0.00 (-0.20) |

| Material property | Posterior stretch slope*$10^{-3}$ ($R^2$) | | | Cervical IO stretch slope*$10^{-3}$ ($R^2$) |
|---|---|---|---|---|
| Direction | Rad | Circ | Norm | Max $\lambda_1$ |
| Cervix E | -1.10* (0.79) | -0.99* (0.66) | 5.71** (0.81) | -2.27** (0.90) |
| Cervix $\nu$ | 0.08*** (1.00) | -0.01 (0.47) | 0.56*** (0.98) | -0.35*** (0.98) |
| Cervix $\xi$ | -0.83* (0.47) | -0.26** (0.66) | 0.35* (0.52) | -1.91* (0.53) |
| Cervix $\alpha$ | -0.02*** (0.98) | 0.00 (-0.18) | 0.00 (0.24) | -0.28*** (1.00) |
| Uterus E | 0.05* (0.95) | 0.05* (0.93) | -0.13* (0.89) | 0.33* (0.94) |
| Uterus $\nu$ | -0.16*** (0.99) | -0.22*** (0.99) | 0.15*** (0.99) | -0.16*** (1.00) |
| Uterus $\xi$ | 0.07*** (0.99) | 0.12*** (0.97) | -0.06*** (0.97) | 0.12*** (0.98) |
| Uterus $\alpha$ | -0.00 (-0.18) | -0.00 (-0.20) | 0.00 (-0.17) | -0.00 (-0.19) |

*TABLES*

Table 2. Slope and $R^2$ of linear fits to stretch against percent change in ultrasonic dimensions, where percent change indicates the percentage change from the median ultrasonic dimension value (Tab. 5). Stretch was analyzed in the anterior, left, and posterior ROIs of the proximal cervix mid-stroma in the surface radial, circumferential, and normal direction. The maximum first principal stretch ($\lambda_1$) was analyzed in the cervical internal os (IO) and in the fetal membrane (FM). Asterisks indicate significance level: $P >$.05 ( ), $P$ = .01-.05 (*), $P$ = .001-.01(**), $P <$.001 (***). Level of slope significance is based on the $t$-statistic where $t = m/\mathrm{SE}(m)$, where $m$ is the slope and $\mathrm{SE}(m)$ is the standard error of the slope. The $P$-value does not necessarily give significance of slope value magnitude from 0.

| Dimension | Anterior stretch slope*$10^{-3}$ ($R^2$) | | | Left stretch slope*$10^{-3}$ ($R^2$) | | |
|---|---|---|---|---|---|---|
| Direction | Rad | Circ | Norm | Rad | Circ | Norm |
| UD1 | 0.27* (1.00) | 0.15 (0.81) | -0.32* (1.00) | 0.28 (0.97) | -0.01 (-0.74) | -0.43* (1.00) |
| UD2 | -0.06*** (0.98) | 0.14** (0.94) | -0.08** (0.95) | 0.13*** (0.98) | 0.00 (-0.27) | -0.09** (0.97) |
| UD3 | 0.03 (0.45) | 0.06** (0.93) | -0.06** (0.91) | 0.03* (0.77) | 0.05* (0.78) | -0.07** (0.95) |
| UD4 | 0.07 (0.89) | -0.98 (0.98) | 0.44* (0.99) | -1.02 (0.98) | -0.03 (0.69) | 0.70* (1.00) |
| UD3a | -0.03 (0.24) | 0.01 (-0.22) | 0.00 (-0.30) | 0.01 (-0.26) | -0.02 (-0.04) | -0.00 (-0.09) |
| UD1a | 0.01 (-0.36) | -0.13* (1.00) | 0.02 (0.83) | -0.11 (0.87) | -0.01 (0.93) | -0.01 (0.51) |
| UD3b | -0.72*** (0.99) | 0.03 (0.60) | -0.08** (0.97) | 0.23*** (0.98) | -0.80*** (1.00) | -0.08** (0.92) |
| UD1b | 0.30*** (0.99) | -0.12*** (0.98) | 0.04** (0.93) | -0.12*** (0.98) | 0.13** (0.96) | 0.09*** (0.99) |
| PCO | -0.18 (0.14) | -0.03 (0.07) | -0.07*** (0.92) | -0.00 (-0.11) | 0.18* (0.68) | -0.01 (0.38) |
| UT1 | 0.09*** (1.00) | 0.22*** (1.00) | -0.06*** (0.99) | 0.17*** (0.99) | 0.11*** (1.00) | -0.02** (0.91) |
| UT2 | 0.30* (0.75) | 0.09 (0.46) | -0.16 (0.67) | 0.00 (-0.32) | 0.23** (0.95) | -0.03 (0.15) |
| UT3 | 0.06*** (1.00) | 0.30*** (1.00) | -0.14*** (1.00) | 0.26*** (1.00) | 0.05** (0.96) | -0.12*** (1.00) |
| UT4 | -0.12* (0.59) | -0.26*** (1.00) | 0.19*** (0.97) | -0.25*** (0.97) | -0.08** (0.77) | 0.19*** (0.98) |
| CL | -0.09* (0.44) | 0.21** (0.78) | -0.12** (0.77) | -0.07 (0.14) | 0.15** (0.73) | -0.11*** (0.78) |
| CD1 | 0.22 (0.96) | 0.37 (0.94) | 0.10 (0.98) | 0.32* (0.99) | 0.34* (1.00) | -0.18** (1.00) |
| CD2 | 0.02** (0.78) | -0.01 (-0.02) | -0.04*** (0.97) | 0.02** (0.81) | -0.00 (-0.12) | -0.04*** (0.98) |
| AUCA | -0.09** (0.92) | -0.00 (-0.24) | 0.34*** (0.99) | -0.07** (0.84) | -0.02* (0.68) | 0.32*** (0.99) |

| Dimension | Posterior stretch slope*$10^{-3}$ ($R^2$) | | | Cervical IO stretch slope*$10^{-3}$ ($R^2$) |
|---|---|---|---|---|
| Direction<br>Max 1st Prin | Rad | Circ | Norm | Max 1st Prin |
| UD1 | -0.35* (0.99) | 0.23 (0.93) | -0.25* (1.00) | 0.27 (0.47) |
| UD2 | 0.03** (0.96) | 0.13** (0.96) | -0.11*** (0.99) | 0.25 (0.54) |
| UD3 | 0.06*** (0.98) | 0.10** (0.94) | -0.08* (0.89) | 0.19* (0.83) |
| UD4 | 0.22 (0.81) | -0.92 (0.97) | 0.30* (1.00) | -1.46 (0.96) |
| UD3a | 0.00 (-0.32) | -0.02 (-0.06) | -0.02 (0.41) | -0.06 (-0.20) |
| UD1a | 0.00 (-0.92) | -0.12 (0.86) | 0.01 (0.43) | 0.00 (-1.00) |
| UD3b | -0.62*** (0.98) | -0.06* (0.83) | -0.22** (0.96) | -0.06 (-0.29) |
| UD1b | -0.11*** (0.98) | -0.04 (0.67) | 0.02 (0.61) | -0.19 (0.26) |
| PCO | 0.32*** (0.98) | 0.02 (0.08) | 0.01 (-0.06) | 0.12 (0.21) |
| UT1 | 0.08*** (0.99) | 0.17*** (1.00) | -0.03* (0.82) | 0.16 (0.47) |
| UT2 | 0.27 (0.66) | -0.03 (-0.09) | -0.15 (0.48) | 0.10 (-0.00) |
| UT3 | 0.03* (0.77) | 0.29*** (1.00) | -0.13*** (0.99) | 0.35*** (0.98) |
| UT4 | -0.11*** (0.96) | -0.10*** (0.96) | 0.10*** (0.99) | -0.35*** (0.97) |
| CL | -0.17* (0.46) | 0.16** (0.74) | -0.01 (-0.13) | 0.29*** (0.78) |
| CD1 | 0.29 (0.98) | 0.24 (0.91) | 0.35** (1.00) | 1.05 (0.94) |
| CD2 | -0.00 (-0.18) | 0.01 (0.29) | -0.04*** (0.90) | 0.35*** (0.97) |
| AUCA | -0.16 (0.50) | 0.10** (0.96) | 0.19 (0.26) | -0.69* (0.72) |
| FM thickness | -0.11*** (1.00) | -0.00 (-0.23) | 0.34*** (1.00) | -0.50*** (0.94) |
